# LRCQ: phenome-wide heritability enrichment of genomic loci from GWAS summary statistics

**DOI:** 10.64898/2026.11.02.763731

**Authors:** Lane Guolan Chen, Thuan Quoc Thach, Pak C Sham

**Affiliations:** Department of Psychiatry, Li Ka Shing Faculty of Medicine, The University of Hong Kong, Hong Kong SAR, China; Centre for PanorOmic Sciences, Li Ka Shing Faculty of Medicine, The University of Hong Kong, Hong Kong SAR, China; State Key Laboratory of Brain and Cognitive Sciences, The University of Hong Kong, Hong Kong SAR, China

## Abstract

Some genetic variants are associated with many traits, but a variant’s GWAS statistics are also inflated by the effects of variants in LD with it and by confounding shared by all variants in a study. Existing measures of pervariant pleiotropy, such as counts of significant associations or multi-trait association tests, do not separate these sources. Stratified LD score regression separates them for one trait at a time and only through predefined annotations. We present LRCQ (LD score regression on *χ*^2^ across many GWAS), which estimates for each locus its phenome-wide heritability enrichment: the variance of its causal effects across a set of traits, relative to an average locus. LRCQ regresses squared Z-scores from > 30 GWAS jointly on squared LD, with pertrait LDSC intercepts and heritabilities fitted in earlier stages. Its pooled least-squares solution has a closed form whose cost does not grow with the number of traits. Its standard errors account for sample overlap and genetic correlation between GWAS. Squared Z-scores identify enrichment at the resolution of LD clumps, not single SNPs, so LRCQ reports clump-level estimates (“omni-loci”). These estimates can be aggregated over any functional annotation or gene set chosen after the analysis.

In simulations with real UK Biobank LD and 30 to 1,000 traits, LRCQ was unbiased for its clump-level estimand. When GWAS share samples its test statistics are strongly right-skewed: normal-reference p-values were up to 10-fold anti-conservative at *P* < 0.001. P-values matched to the exact third cumulant restored type-I error and FDR control, and matching the fourth cumulant as well kept the far tail within 1.6-fold of nominal down to *P* = 10^−4^. Score-type tests, which evaluate the variance under the null hypothesis, kept power at the most enriched loci. Model-based standard errors were calibrated once the enrichment estimates plugged into them kept the polygenic background. It ranked enriched loci better than mean *χ*^2^, LD-normalised *χ*^2^, multi-trait omnibus tests and significant-association counts in 13 of 16 scenarios. Its main vulnerability was a misspecified LDSC intercept. Applied to 444 Pan-UK Biobank European-ancestry GWAS, LRCQ found strongly non-uniform phenome-wide enrichment: the SD of true clump-mean enrichment was 8.3, against a genome mean of 1. At a tag-level FDR of 5%, 881 LD clumps were enriched above the uniform expectation; they fall in 638 distinct loci, including FTO, AXIN2, HBS1L-MYB, CST3, CELSR2-SORT1, PCSK9 and MC4R. These ranked only moderately with mean *χ*^2^ and poorly with counts of significant associations. Gene-level phenome-wide enrichment rose with loss-of-function constraint. Phenome-wide heritability was 3-5-fold enriched in coding, conserved, 5′ UTR and promoter sequence, and depleted in repressed chromatin. LRCQ turns hundreds of biobank GWAS into an LD-corrected map of where phenome-wide genetic variance lies, and complements its companion method LRCP, which estimates genetic-effect correlations between loci.

## Introduction

Most common variants associated with complex traits are associated with more than one trait. Pleiotropy is the rule rather than the exception in the human genome (Sivakumaran et al. 2011; Solovieff et al. 2013; Pickrell et al. 2016; Visscher & Yang 2016; Chesmore et al. 2018; Watanabe et al. 2019). Biobank-scale GWAS of hundreds of phenotypes in the same individuals now make it possible to study pleiotropy across much of the phenome (Bycroft et al. 2018; Karczewski et al. 2025). One simple observation from such data motivates this work. The GWAS test statistics of some variants are inflated far beyond their LD-expected level, and consistently across very different traits. This pattern is what the omnigenic model predicts for variants with broad regulatory consequences (Boyle et al. 2017; Liu et al. 2019). But inflation of a variant’s *χ*^2^ statistics across traits has at least three sources: larger causal effects of the variant itself on many traits, the effects of other variants it tags through linkage disequilibrium (LD), and confounding such as population stratification or cryptic relatedness that inflates all statistics of a GWAS. Separating the first source from the other two requires a model.

For a single trait, this separation is well developed. LD score regression (LDSC) uses the dependence of *χ*^2^ on LD score to distinguish polygenic signal from confounding (Bulik-Sullivan et al. 2015a). Stratified LDSC partitions heritability across functional annotations (Finucane et al. 2015; Gazal et al. 2017). Likelihood and Bayesian methods refine the same idea (Speed et al. 2017; Zeng et al. 2018; Ning et al. 2020). All of these methods either assume that per-SNP effect variance is constant, or let it vary only through a small number of annotations or a fixed function of allele frequency and LD. All are fitted one trait at a time. Per-SNP prior variances used in functionally informed fine-mapping are likewise trait-specific and limited by annotations (Weissbrod et al. 2020). Cross-trait extensions estimate genetic covariances between traits, summed over SNPs or over LD blocks (Bulik-Sullivan et al. 2015b; Shi et al. 2017; Werme et al. 2022). They describe how traits share genetics, not how strongly a given variant acts across traits.

Existing per-variant measures of pleiotropy do not fill this gap. Counts of genome-wide significant associations (Watanabe et al. 2019) depend on sample size, heritability and the density of the trait collection. They are also inflated for variants that merely tag pleiotropic neighbours. Multi-trait association tests, such as CPASSOC (Zhu et al. 2015), metaUSAT (Ray & Boehnke 2018) and PLEIO (Lee et al. 2021), and per-variant pleiotropy scores such as HOPS (Jordan et al. 2019), test whether a variant is associated with any or many traits. They do not estimate an interpretable quantity, and they do not remove the contribution of LD. Multivariate shrinkage and multi-trait fine-mapping (Urbut et al. 2019; Foley et al. 2021; Arvanitis et al. 2022; Zou et al. 2026) give per-variant, per-trait posterior effects. They are applied to selected loci with sparse causal signals, and they do not scale to hundreds of traits genome-wide.

Here we introduce LRCQ, a method that estimates for every SNP its *phenome-wide heritability enrichment w*: the variance of the SNP’s causal effect across a collection of traits, relative to an average SNP, so that the genome-wide mean of *w* is 1. LRCQ is built on a random-effects model for the SNP-by-trait effect matrix with a matrix-normal second-moment structure, in the tradition of multivariate linear mixed models (Stephens 2013; Zhou & Stephens 2014; Furlotte & Eskin 2015). Unlike those models, it targets the SNP-side covariance rather than the trait-side covariance, and it needs only GWAS summary statistics and an LD reference. Under this model the expected *χ*^2^ of each SNP in each trait is a known linear function of *w* through squared LD, scaled by the trait’s sample size and heritability and shifted by its LDSC intercept. LDSC is the special case *w* ≡ 1, and stratified LDSC the case where *w* is constrained to the span of annotations. LRCQ instead lets *w* vary freely across the genome and makes this estimable by pooling dozens to hundreds of traits. Its companion method, LRCP, uses Z-score cross-products within the same framework to estimate the genetic-effect correlation between pairs of SNPs (Chen et al. 2026).

We show how much of *w* GWAS summary statistics can identify, and with what precision. At HapMap3 density, squared Z-scores identify *w* only up to the resolution of LD clumps, so LRCQ reports clump-level enrichment (“omni-loci”). Z-scores of biobank traits are strongly correlated through shared samples and genetic correlation, so the effective number of traits is much smaller than their count. We derive standard errors that account for this, exact for Gaussian Z-scores, and a skewness correction for the test statistics, which are far from normal when the effective number of traits is small. Because per-locus precision is limited, aggregating LRCQ estimates over functional annotations, genes and phenotype categories is where most of its power lies; these aggregates are unbiased and need no annotation model. In simulations with real UK Biobank LD and 30 to 1,000 traits, LRCQ was unbiased for its clump-level estimand and, with moment-matched p-values, held type-I error and FDR. In 13 of 16 scenarios it ranked enriched loci better than mean *χ*^2^, LD-normalised *χ*^2^, significant-association counts and a multi-trait omnibus test. Its main vulnerability was a misspecified LDSC intercept. Applied to 444 Pan-UK Biobank traits, LRCQ finds strongly non-uniform phenome-wide enrichment, with 881 enriched clumps in 638 loci. Phenome-wide heritability is concentrated in coding, conserved and promoter sequence and near loss-of-function-intolerant genes. Which loci rank highest depends on the trait collection, while these functional patterns do not.

## Results

### Some loci are associated with many traits beyond what LD predicts

We first asked whether the pattern that motivated this work is visible in Pan-UK Biobank data. For each HapMap3 SNP *k* and each of 444 European-ancestry GWAS, 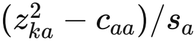 estimates the LD-weighted heritability around SNP *k* in trait *a*. When every SNP contributes equally (**w** ≡ **1**), its expectation equals the SNP’s LD score. Pooled across traits, this quantity rose with LD score as LDSC predicts, but with a long right tail of SNPs carrying far more signal than their LD score implies (Fig. 1a,b). Whether this tail reflects SNPs that are themselves pleiotropic, their LD neighbours, or noise shared through overlapping samples cannot be read from the raw statistics (see “Why naive replication fails in a single biobank” below). LRCQ is designed to separate these sources.

**Figure 1.**
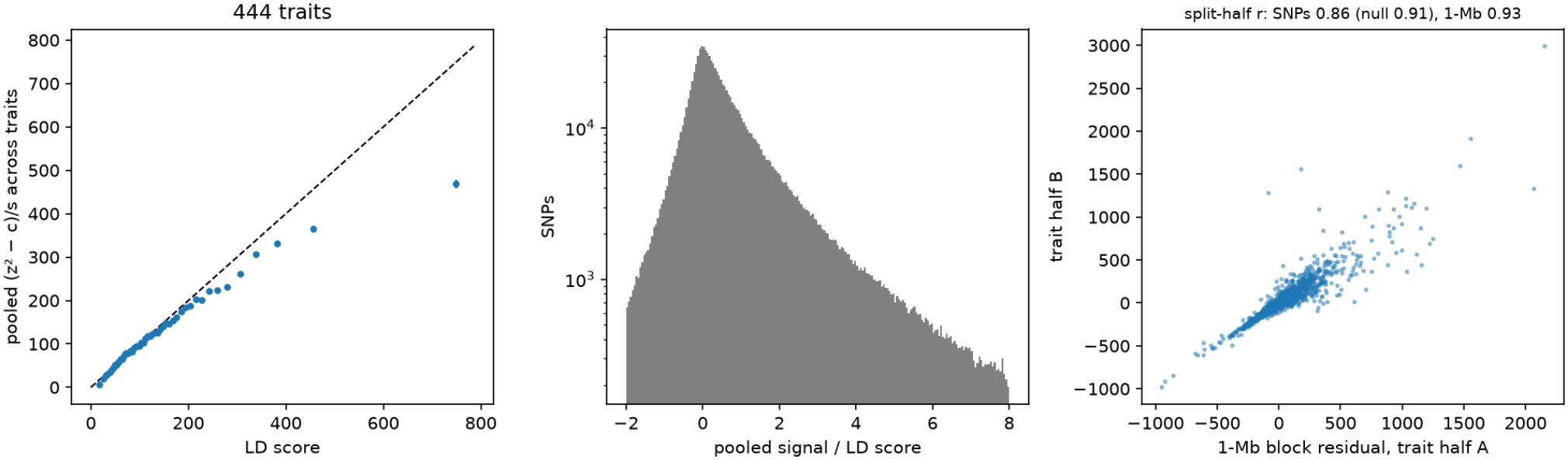
Some loci are associated with many traits beyond their LD. (a) Per-SNP pooled signal 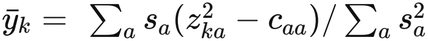 across 444 Pan-UKB traits, averaged within 50 LD-score bins (95% CI). The dashed line is the LDSC expectation under uniform enrichment. (b) Distribution of pooled signal divided by LD score (log scale). (c) Split-half replication: residuals of 1-Mb block means after regressing out LD score, for two random halves of the traits. The title gives the mean SNP-level and block-level correlation over 20 splits, and the exact null correlation under uniform enrichment.

### LRCQ in brief

LRCQ models the SNP-by-trait matrix of causal effects as random, with a SNP covariance shared by all traits up to scale (Methods). The phenome-wide enrichment *w*_*s*_ of SNP *s* is its causal-effect variance across the traits analysed relative to an average SNP, so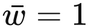. Each trait’s expected *χ*^2^ at SNP *k* is its LDSC intercept plus 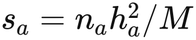 times (**Dw**)_*k*_, where **D** holds squared LD. Given genome-wide heritabilities and intercepts (Stages 1-2), Stage 3 solves this linear system for **w** by pooling all traits, one LD window at a time. The pooled solution has a closed form that costs one window-sized solve however many traits are analysed (Theorem 3.1). Standard errors account for sample overlap and genetic correlation between traits (Theorem 3.10), and p-values account for the skewness of the estimates (Methods).

### Per-SNP enrichment is identified only at LD-clump resolution

At HapMap3 density, squared LD is numerically singular. In a representative 3-Mb window of UK Biobank insample LD (1,106 SNPs), the condition number of **D** was about 5 × 10^13^, and the variance inflation of an exact per-SNP solve, (**D**^−1^)_*ii*_, had median 13 and 90th percentile 364. In simulation, the exact solve gave estimates with SD 46 and correlation 0.17 with the truth (Supplementary Table S1). The split of enrichment between SNPs in tight LD is not identifiable from Z-scores. The data identify its total. Simulations of the per-SNP estimator show the same limit from another angle: per-SNP OLS lost accuracy as the genome-wide SNP count grew, from *R*^2^ 0.97 with the truth at 10^4^ SNPs to 0.46 at 10^7^ (Methods). This is because the per-SNP signal 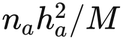 shrinks while sampling noise does not.

We therefore estimate one parameter per LD-pruned tag SNP (*r*^2^ < 0.5), using the responses of all SNPs. This leaves 273 tags in the example window, with median variance inflation 1.41. The estimand is the enrichment of the tag’s LD clump, *W*_*t*_ (Methods); we call tags with large *W*_*t*_ omni-loci. Ridge regularisation of the per-SNP solve is the alternative. It spreads enrichment evenly within clumps, and we use it as a sensitivity analysis.

### Simulations with real LD

We simulated summary statistics for *q* = 300 traits with UK Biobank-scale samples (*n* = 3 × 10^5^) over three 3-Mb windows of real UK Biobank LD (3,178 HapMap3 SNPs). In the baseline, 2% of SNPs had non-zero log-normal enrichment, concentrated in a synthetic four-level annotation. Sixteen scenarios varied one factor at a time (Methods; Table 1).

**Table 1.** Simulation summary. Per scenario: Pearson r, slope, bias and RMSE of OLS; WLS and equal-weight r; SD of standardised errors (background-filled and method-A plug-ins); type-I error at null tags; coverage; AUC (top 10%) for LRCQ and comparators; Method D discoveries, FDP and power.

| Scenario | OLS *r* | OLS slope | OLS bias | OLS RMSE | WLS *r* | Equal-weight *r* | SD std. error (bg-filled) | SD std. error (method A) | Type-I at 5% | Coverage 95% (bg-filled) | Coverage 95% (method A) | AUC10 LRCQ | AUC10 mean χ<sup>2</sup> | AUC10 LD-norm. χ<sup>2</sup> | AUC10 omnibus | AUC10 no. significant | Method D discoveries | Method D FDP | Method D power |
|---|---|---|---|---|---|---|---|---|---|---|---|---|---|---|---|---|---|---|---|
| Baseline | 0.984 | 1.009 | 0.006 | 3.90 | 0.970 | 0.978 | 1.043 | 1.145 | 0.036 | 0.946 | 0.927 | 0.974 | 0.921 | 0.879 | 0.921 | 0.725 | 45.0 | 0.004 | 0.244 |
| *q* = 30 | 0.847 | 0.959 | -0.059 | 12.36 | 0.849 | 0.800 | 1.059 | 1.180 | 0.046 | 0.946 | 0.922 | 0.859 | 0.880 | 0.818 | 0.880 | 0.588 | 26.8 | 0.029 | 0.137 |
| *q* = 100 | 0.949 | 1.007 | 0.014 | 7.05 | 0.937 | 0.935 | 1.058 | 1.167 | 0.041 | 0.944 | 0.923 | 0.935 | 0.909 | 0.858 | 0.909 | 0.660 | 36.8 | 0.008 | 0.193 |
| *q* = 1000 | 0.995 | 1.004 | 0.003 | 2.17 | 0.980 | 0.993 | 1.042 | 1.146 | 0.030 | 0.944 | 0.924 | 0.995 | 0.927 | 0.896 | 0.927 | 0.789 | 44.3 | 0.000 | 0.241 |
| 0.5% non-null | 0.992 | 0.997 | -0.008 | 5.08 | 0.970 | 0.991 | 1.059 | 1.102 | 0.040 | 0.944 | 0.937 | 0.891 | 0.849 | 0.819 | 0.849 | 0.667 | 11.1 | 0.001 | 0.099 |
| 10% non-null | 0.949 | 1.003 | 0.007 | 3.51 | 0.945 | 0.926 | 1.025 | 1.231 | 0.028 | 0.945 | 0.899 | 0.989 | 0.942 | 0.897 | 0.942 | 0.709 | 132.7 | 0.010 | 0.408 |
| All non-null | 0.810 | 0.999 | 0.009 | 3.28 | 0.813 | 0.728 | 0.992 | 1.239 | 0.028 | 0.953 | 0.889 | 0.943 | 0.904 | 0.645 | 0.904 | 0.623 | 390.9 | 0.028 | 0.544 |
| UK Biobank-like traits | 0.879 | 1.012 | 0.024 | 11.15 | 0.846 | 0.819 | 1.011 | 1.089 | 0.052 | 0.945 | 0.934 | 0.894 | 0.891 | 0.844 | 0.919 | 0.706 | 37.9 | 0.095 | 0.162 |
| Intercepts 1.1, supplied | 0.983 | 0.998 | -0.003 | 4.12 | 0.972 | 0.974 | 1.044 | 1.152 | 0.035 | 0.944 | 0.922 | 0.978 | 0.926 | 0.880 | 0.926 | 0.732 | 42.8 | 0.001 | 0.225 |
| Intercepts 1.1, assumed 1 | 0.980 | 0.999 | 0.679 | 4.13 | 0.968 | 0.971 | 1.055 | 1.243 | 0.076 | 0.929 | 0.883 | 0.973 | 0.922 | 0.847 | 0.922 | 0.738 | 49.0 | 0.008 | 0.261 |
| Local ρ = 0.5 | 0.932 | 0.988 | -0.037 | 8.19 | 0.911 | 0.924 | 1.561 | 1.741 | 0.059 | 0.863 | 0.837 | 0.919 | 0.915 | 0.873 | 0.915 | 0.720 | 35.9 | 0.028 | 0.190 |
| Cross-block ρ | 0.983 | 0.996 | -0.013 | 3.86 | 0.973 | 0.976 | 1.039 | 1.130 | 0.036 | 0.944 | 0.924 | 0.976 | 0.922 | 0.882 | 0.922 | 0.719 | 45.7 | 0.004 | 0.247 |
| Heterogeneous **w** | 0.977 | 0.994 | -0.023 | 3.86 | 0.969 | 0.968 | 1.043 | 1.146 | 0.036 | 0.945 | 0.925 | 0.980 | 0.933 | 0.895 | 0.933 | 0.769 | 54.2 | 0.004 | 0.289 |
| LD from 500 people | 0.963 | 0.995 | 0.028 | 5.70 | 0.944 | 0.957 | 1.345 | 1.524 | 0.070 | 0.869 | 0.833 | 0.947 | 0.924 | 0.902 | 0.924 | 0.723 | 40.3 | 0.015 | 0.204 |
| Tags *r*<sup>2</sup> < 0.8 | 0.966 | 1.003 | 0.009 | 4.46 | 0.959 | 0.945 | 1.019 | 1.086 | 0.046 | 0.948 | 0.933 | 0.840 | 0.869 | 0.816 | 0.869 | 0.698 | 37.9 | 0.012 | 0.239 |
| Tags *r*<sup>2</sup> < 0.2 | 0.988 | 0.996 | -0.016 | 4.19 | 0.967 | 0.986 | 1.057 | 1.164 | 0.024 | 0.943 | 0.924 | 0.998 | 0.974 | 0.944 | 0.974 | 0.773 | 60.3 | 0.003 | 0.401 |

OLS: tag-level OLS estimator (primary). Bias and RMSE in enrichment units. SD std. error: SD of $(\hat{W}_t - W_t)/\widehat{\mathrm{SE}}$ with the background-filled or method-A plug-in of **w**. Type-I error of the one-sided test at null tags with the background-filled plug-in. AUC10: area under the ROC curve for identifying the 10% most enriched tags. Method D: rectified discoveries at FDR 5%. 50 replicates per scenario.

#### Accuracy

The tag-level OLS estimator was unbiased for its clump estimand in every scenario where the model was correctly specified. Its slope on the truth ranged from 0.96 to 1.01, and its Pearson correlation with the truth was 0.98 in the baseline (Fig. 2a; Table 1). Accuracy grew with the number of traits: correlation 0.85 at *q* = 30 , 0.95 at *q* = 100 and 0.995 at *q* = 1000. It fell when traits were UK Biobank-like (five clusters with *r*_*g*_ = 0.5 and full sample overlap; *r* = 0.88), because correlated traits carry less independent information than their number suggests (Supplementary Note §S9.1). It also fell under an infinitesimal architecture (*r* = 0.81), where enrichment varies little between tags. Contrary to the usual expectation, the one-step WLS estimator was slightly attenuated (slope 0.93-0.97) and had a higher RMSE than OLS in all but three scenarios. Its weights depend on the same data as its responses. Heavy-tailed effects also make Gaussian weights inaccurate near causal variants (Supplementary Note §S5). OLS is therefore our primary estimator. The equal-weight estimator was unbiased for the simple trait average (slope 0.97-1.02), at a modest cost in precision (baseline *r* = 0.98).

**Figure 2.**
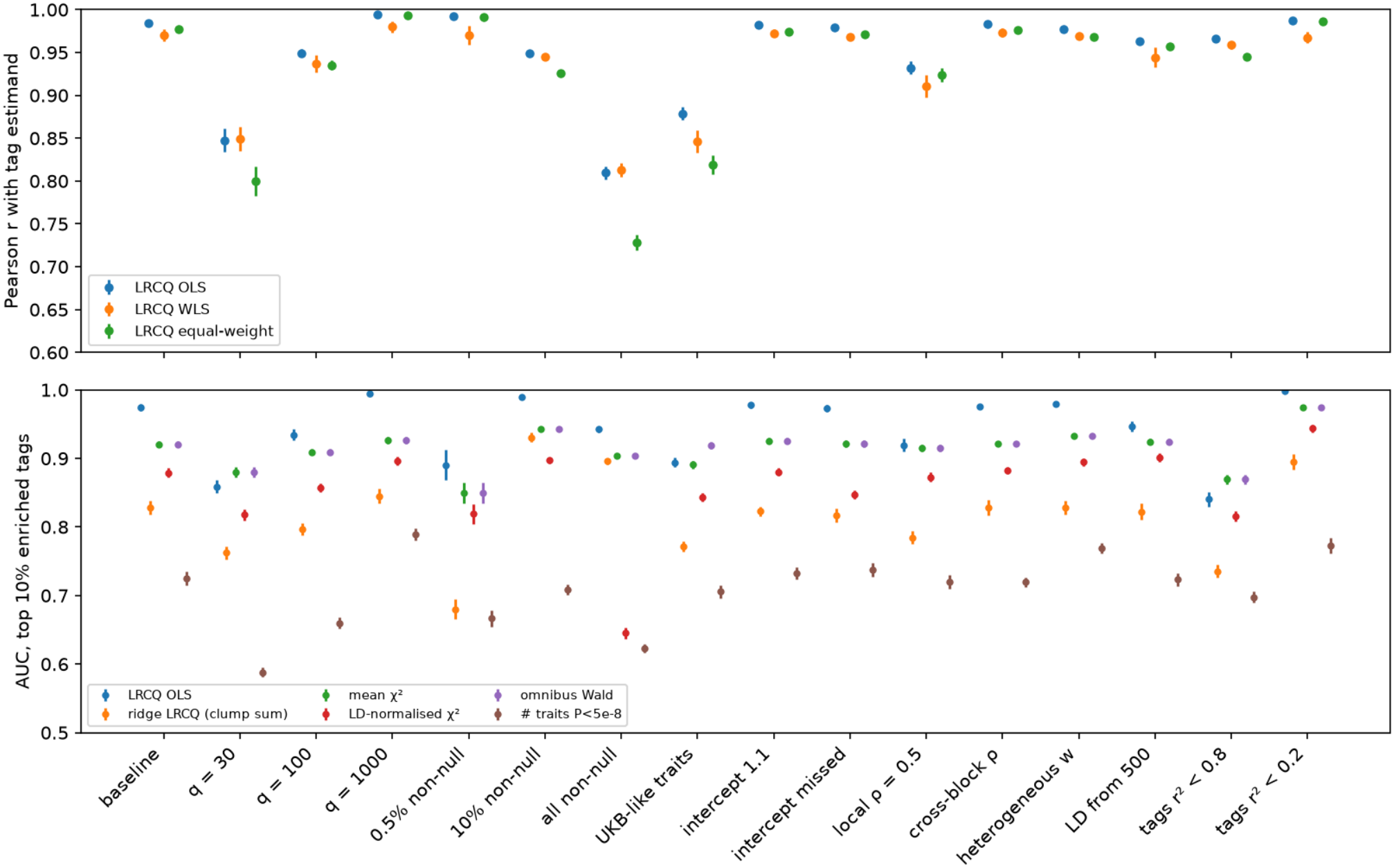
Accuracy and ranking in simulation. (a) Pearson correlation of tag-level LRCQ estimates (OLS, one-step WLS, equal-weight) with the tag-set estimand, across 16 scenarios (mean and 95% CI over 50 replicates). (b) AUC for identifying the 10% most enriched tags: LRCQ, per-SNP ridge LRCQ summed within clumps, and four per-SNP comparators.

#### Calibration

At null tags, one-sided normal-reference tests of *W*_*t*_ > 0 had type-I error 0.024-0.059 at 5% in all scenarios except a misspecified intercept (0.076) and LD from a 500-person reference (0.070). The 5% level hides a distortion in the tail, however (Fig. 3d-g). The estimator is a quadratic form in Z-scores, and when the effective number of traits is small its distribution is right-skewed (median skewness 1.7 for UK Biobank-like traits and 1.3 with the Pan-UKB trait structure, against 0.15 for independent traits). A normal reference is then conservative for most null tags and anti-conservative in the tail, so the QQ plot first falls below the diagonal and then crosses above it. At null tags in the UK Biobank-like scenario, normal-reference *P* < 0.001 occurred 10 times more often than nominal (0.0104), and with the real Pan-UKB intercept and genetic-covariance matrices 9 times (0.0093). Skewness-corrected p-values from the exact third cumulant (Methods) restored calibration: 0.0006 and 0.0000 at *P* < 0.001, and 0.039 and 0.031 at *P* < 0.05, slightly conservative (Supplementary Table S4). With the Pan-UKB trait structure, the tag estimates correlated 0.72 with the truth (slope 1.03), and adding realistic Stage 1-2 errors, multiplicative *h*^2^ errors with SD 20% or intercept errors drawn from Pan-UKB’s own intercept SEs, left the corrected tests calibrated (*P* < 0.001: 0.0007 and 0.0000). Errors in *h*^2^ attenuated the slope to 0.94, a common-scale effect that renormalisation to mean 1 removes. Model-based standard errors need an estimate of **w** themselves. With the background-filled plug-in (Methods) they were calibrated: in every scenario without misspecified intercepts, correlated effects or a small LD reference, the SD of standardised errors was 0.99-1.06 and 95% coverage 0.94-0.95 (Fig. 3a,b). Plugging in only the estimates that survive rectification method A, which drops the polygenic background, made SEs 10-25% too small (SD 1.09-1.24; coverage 0.89-0.94), most under dense enrichment. With the Pan-UKB trait structure, null tags had standardised-error SD 0.91-0.97 with the background-filled plug-in and 1.07-1.19 with method A, across enrichment from 2% non-null SNPs to **w** ≡ **1** (Supplementary Table S6). A delete-one-cluster jackknife over traits did not help: under UK Biobank-like sample overlap its SEs were less than half the true SD, because overlapping trait clusters are not independent (Supplementary Table S3), so we use model-based SEs.

**Figure 3.**
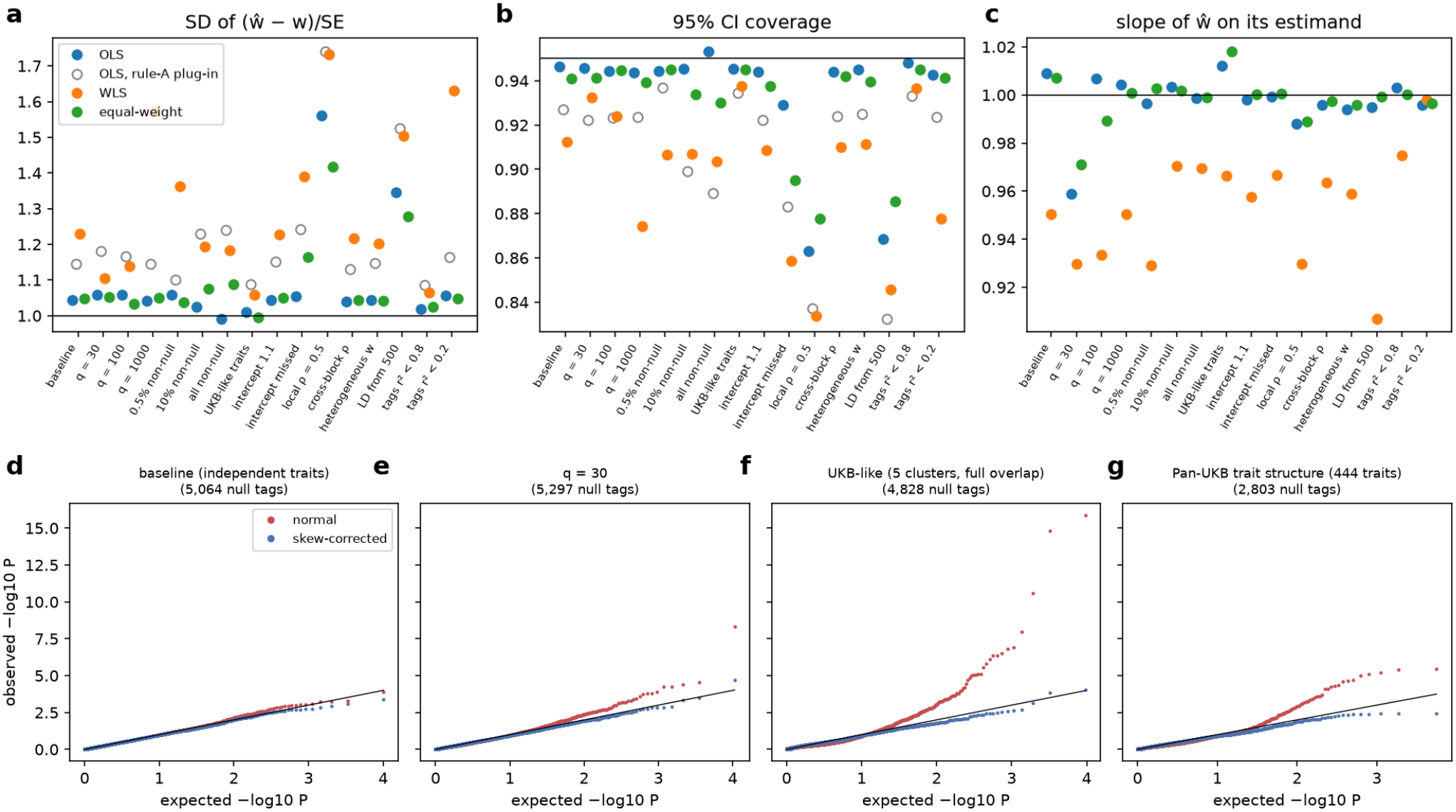
Calibration in simulation. (a) SD of standardised errors 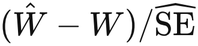, using model-based Ses (Theorem 3.10) with the background-filled plug-in of **w**; open grey circles: OLS with the method-A plug-in. (b) Coverage of 95% confidence intervals. (c) Slope of *Ŵ* on its estimand (for the equal-weight estimator, on the equal-weight estimand). (d-g) QQ plots of one-sided p-values for *H*_0_ : *W*_*t*_ ≤ 0 at null tags (tag-set estimand *∣W*_*t*_*∣* < 0.01), pooled over replicates, with a normal reference (red) and the skewness-corrected three-moment reference (blue): (d) baseline, independent traits; (e) *q* = 30; (f) UK Biobank-like traits (five clusters, full sample overlap); (g) the Pan-UKB 444-trait intercept and genetic-covariance matrices, sample sizes and *h*^2^.

#### Ranking omni-loci

For identifying the 10% most enriched tags, LRCQ outperformed every per-SNP pleiotropy measure we compared: mean *χ*^2^ across traits, LD-score-normalised *χ*^2^, the multi-trait omnibus Wald statistic and the number of genome-wide significant traits. Baseline AUCs were 0.974 for LRCQ, against 0.921, 0.879, 0.921 and 0.725 (Fig. 2b). Among truly enriched tags, LRCQ’s rank correlation with the truth was 0.83, against 0.70 for mean *χ*^2^. LRCQ’s advantage grew with the number of traits (AUC 0.995 at *q* = 1000) and with polygenicity. The comparators were as good or better in three settings: with 30 traits (0.86 vs 0.88), with coarser tags at *r*^2^ < 0.8 (0.84 vs 0.87), and for UK Biobank-like correlated traits, where the omnibus statistic, which models trait correlation, ranked slightly better (0.92 vs 0.89). Counts of significant associations were the worst measure in every scenario, because they reward large single-trait effects and LD with them.

### Robustness

- *Population stratification*. When the inflated intercepts (1.1) were supplied, the estimates were unbiased. When the intercepts were wrongly assumed to be 1, every tag was biased upward by 0.68 on average, and the bias was largest for low-LD tags (type-I error 0.076 with the background-filled plug-in, 0.11 with method A). Intercept estimation is therefore the critical input.
- *Genetic-effect correlation*. Correlation between SNPs in different windows had no effect, as theory predicts. Local correlation (*ρ* = 0.5 among clusters of five non-null SNPs in LD) reduced accuracy (*r* = 0.93) and inflated the spread of standardised errors (SD 1.74). This is consistent with the known bias **D**^−1^ diag(**RCR**) (Theorem 2), which affects only those clusters.
- *Heterogeneous enrichment*. When half of the traits had an independent enrichment profile, OLS estimated the power-weighted mixture without loss of accuracy (*r* = 0.98).
- *Out-of-sample LD*. LD estimated from 500 people reduced accuracy only modestly (*r* = 0.96). It inflated the standardised error spread to 1.52, because LD noise is not in the variance formula. We use 1000 Genomes LD for the genome-wide analysis and in-sample UK Biobank LD as a sensitivity analysis (see below).
- *Tag threshold*. A stricter threshold (*r*^2^ < 0.2) gave coarser but better-ranked clumps (AUC 0.998). A looser threshold (*r*^2^ < 0.8) gave finer but noisier ones (AUC 0.84).

### Negative estimates and discovery

- Rectification method C kept ∑ *Ŵ*at its unrectified value (ratio 1.00 in the baseline, 1.01 for UK
- Biobank-like traits), as expected from Supplementary Note §S3.8. Methods A and B lost 34% and 6% of the total in the baseline, and method B removed all of it for UK Biobank-like traits (Supplementary Table S5).
- The knockoff-style threshold (Method D) kept the realised false discovery proportion below the 10% target in every scenario (0-3.4%). It detected 6-50% of tags with *W*_*t*_ > 1, in line with the per-clump power calculations (Supplementary Note §S9.2).

#### Annotation enrichment

The clump-level enrichment 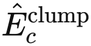 recovered its clump-level target without bias in every scenario. Because the synthetic annotation splits LD clumps, that target is attenuated toward 1 relative to the SNP-level enrichment: 3.9 against a true 5.2 for the top category in the baseline (Fig. 4). The annotation regression recovered the SNP-level enrichment (5.2) but with two to three times the standard deviation. The two estimators trade bias for variance in the way Supplementary Note §S7.2a predicts. We report both, interpreting *Ê*^clump^ as a conservative estimate and the regression as the unbiased one.

**Figure 4.**
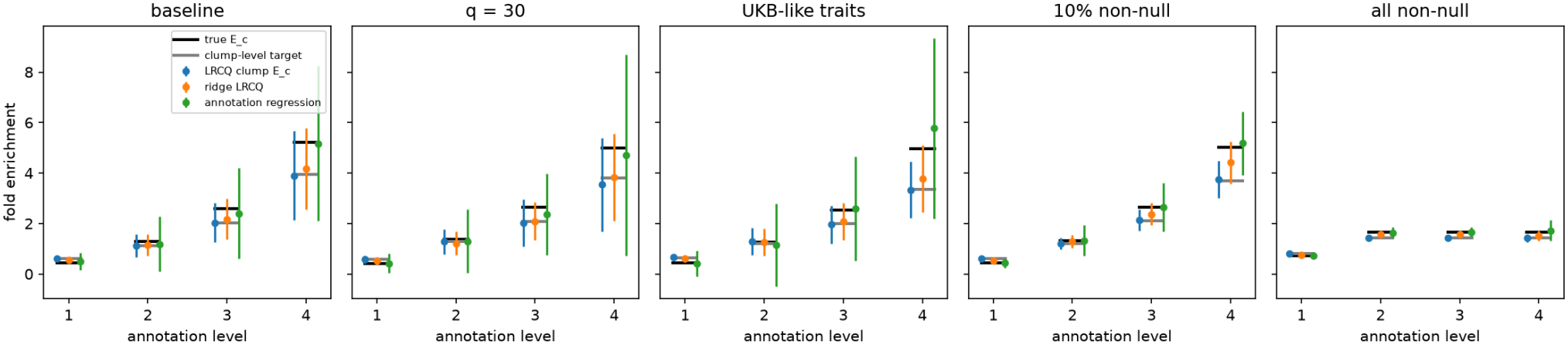
Annotation enrichment in simulation. Fold enrichment of four synthetic annotation levels: SNP-level truth (black), clump-level target (grey), clump-level LRCQ *Ê* ^clump^, per-SNP ridge LRCQ, and annotation regression (oracle categories). Error bars: SD across replicates (three 3-Mb windows each).

### Phenome-wide enrichment across 444 Pan-UK Biobank traits

We applied LRCQ to Pan-UKB European-ancestry GWAS of 444 phenotypes (median *n* = 413,174; range 35,367-420,531; Supplementary Table S2). The traits passed Pan-UKB’s EUR quality control; eight prescription-medication traits whose files repeatedly failed to download were excluded. The analysis covered 1,084,528 HapMap3 SNPs outside the MHC in 301,693 tags (median clump size 2; 41% singletons), with LD from 1000 Genomes Europeans. Per-trait intercepts and heritabilities came from Pan-UKB’s LDSC. Our reestimation reproduced them (*r* = 0.992 for *h*^2^, 0.985 for intercepts), and cross-trait LDSC on all 98,346 trait pairs supplied the off-diagonal intercepts and genetic covariances. Because almost all traits share the same UK Biobank participants, the median SE of a singleton tag’s enrichment was 14.5. This is far above the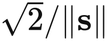 expected for independent GWAS, and in line with the effective-trait calculations of Supplementary Note §S9.1.

**Enrichment is far from uniform**. Squared standardised deviations of *Ŵ*_*t*_ from its expectation under uniform enrichment summed to 336,696 over 301,693 tags, against an expectation of about 1.0 per tag. After subtracting sampling variance, the SD of true clump-mean enrichment was 8.3 (method of moments). With score-type tests (Methods), 881 tags passed a 5% Benjamini-Hochberg threshold for enrichment above the uniform expectation (222 tags at Bonferroni *P* < 0.05), and 1,799 tags had *W*_*t*_ significantly greater than 0 (Table 2; Fig. 5). The FDR applies to tags; merging tags within 1 Mb places the 881 tags in 638 distinct loci and the 1,799 tags in 1,145 loci, and these locus counts do not carry the same error rate. Under the null hypothesis the tag statistics were strongly skewed and heavy-tailed (median skewness 1.16, 10th-90th percentile 0.43-1.67; median excess kurtosis 4.2 among tags with score *z* > 2), as the Pan-UKB-structure simulations predicted. Normal-reference p-values would have declared 5,178 tags in 1,961 loci. In a random 5% of tags, *P* < 0.001 occurred 5.7 times more often than nominal, against 17.1 times with a normal reference (Supplementary Fig. S1). This excess is expected from truly enriched clumps. Below the tail, the QQ curves lie on or under the diagonal, because the null hypotheses are composite: most clumps are less enriched than average, and their p-values are larger than uniform. In simulation under uniform enrichment, the kurtosis-matched p-values were calibrated at *P* = 0.01 but still exceeded nominal rates 1.3-fold at 10^−3^ and 1.6-fold at 3 × 10^−4^ (Supplementary Table S7). Doubling every p-value, as an allowance for this residual excess, leaves 669 tags in 514 loci enriched above the uniform expectation at FDR 5%. This is a sensitivity analysis rather than a bound: with a known covariance, the kurtosis-matched reference exceeded nominal rates by up to about twofold at the Bonferroni level for some combinations of LD and trait correlation (Supplementary Fig. S2). The 881st BH threshold, *P* ≈ 1.5 × 10^−4^, lies within the simulated range; the very small *q* values of the top loci in Table 2 rest on the moment-matched reference beyond it. For 5.0% of tags the projected uniform null 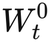 is negative, and the score test then uses the nearest nonnegative null configuration (Methods). Stating the mean of that configuration with the uncorrected **R** *∘* **R** of the 503-sample reference panel instead of the corrected **D** would leave 322-421 tags above the uniform expectation and 568-770 with *W*_*t*_ > 0 (Supplementary Table S8), because it treats reference-panel noise as LD.

**Table 2.**
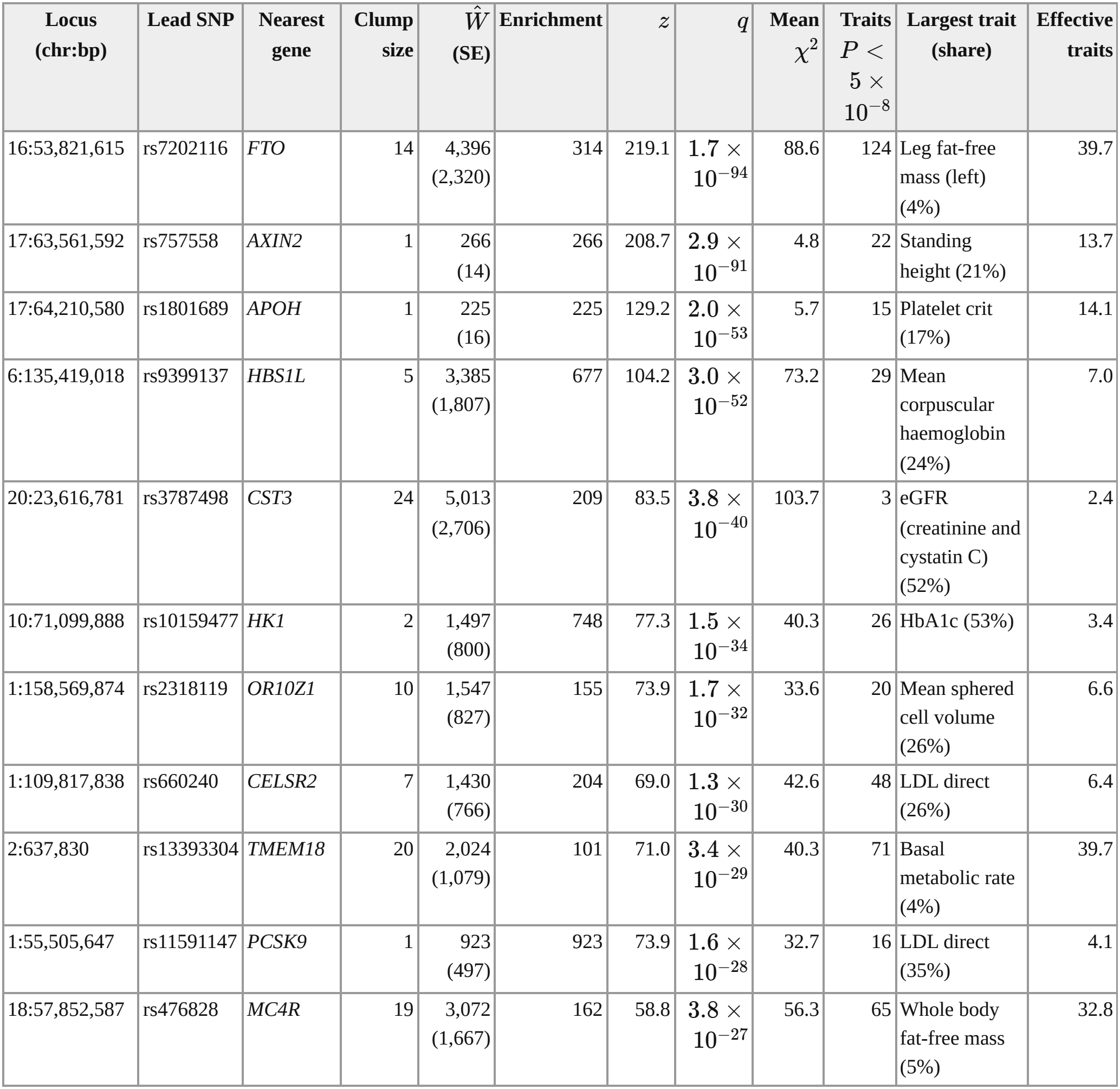
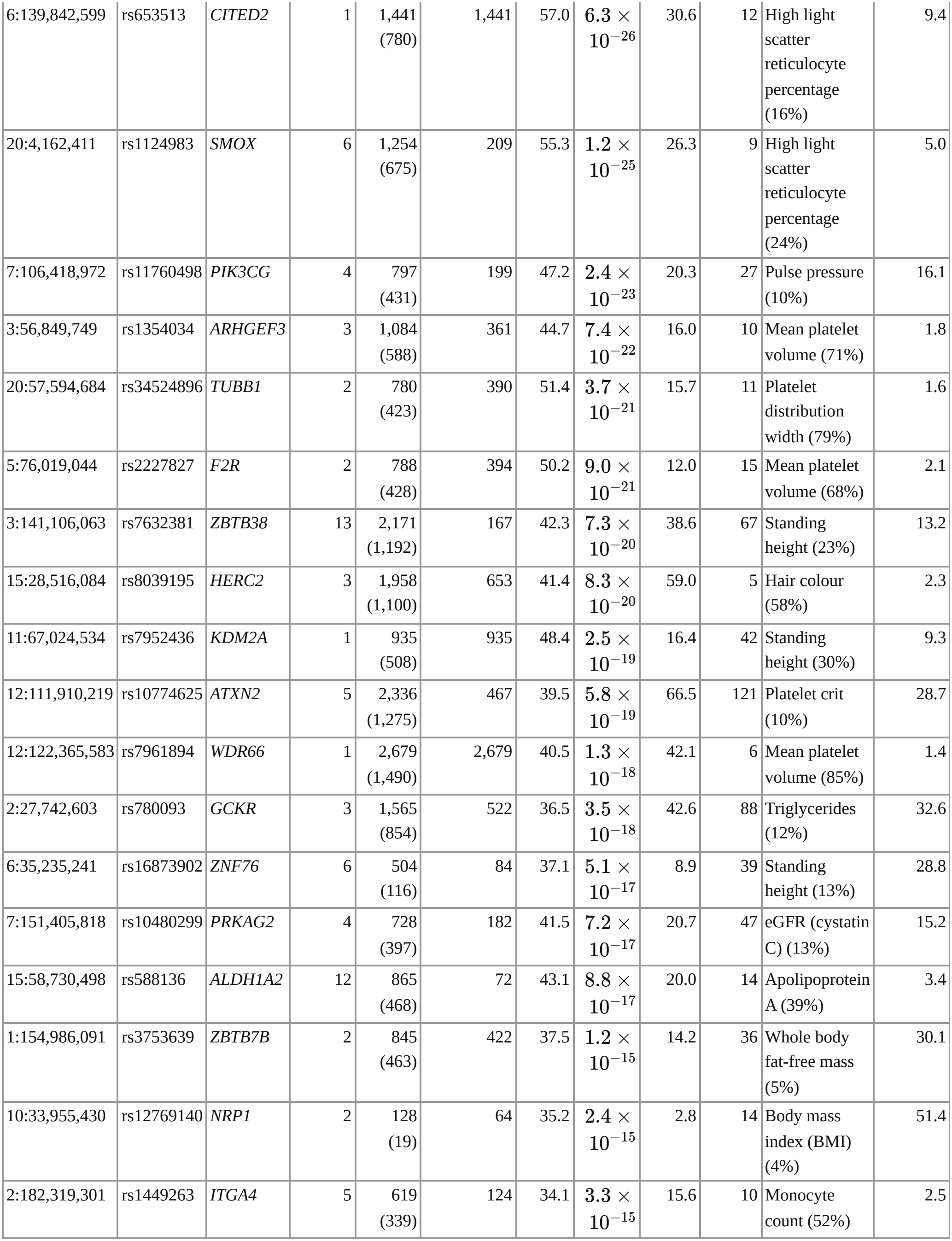
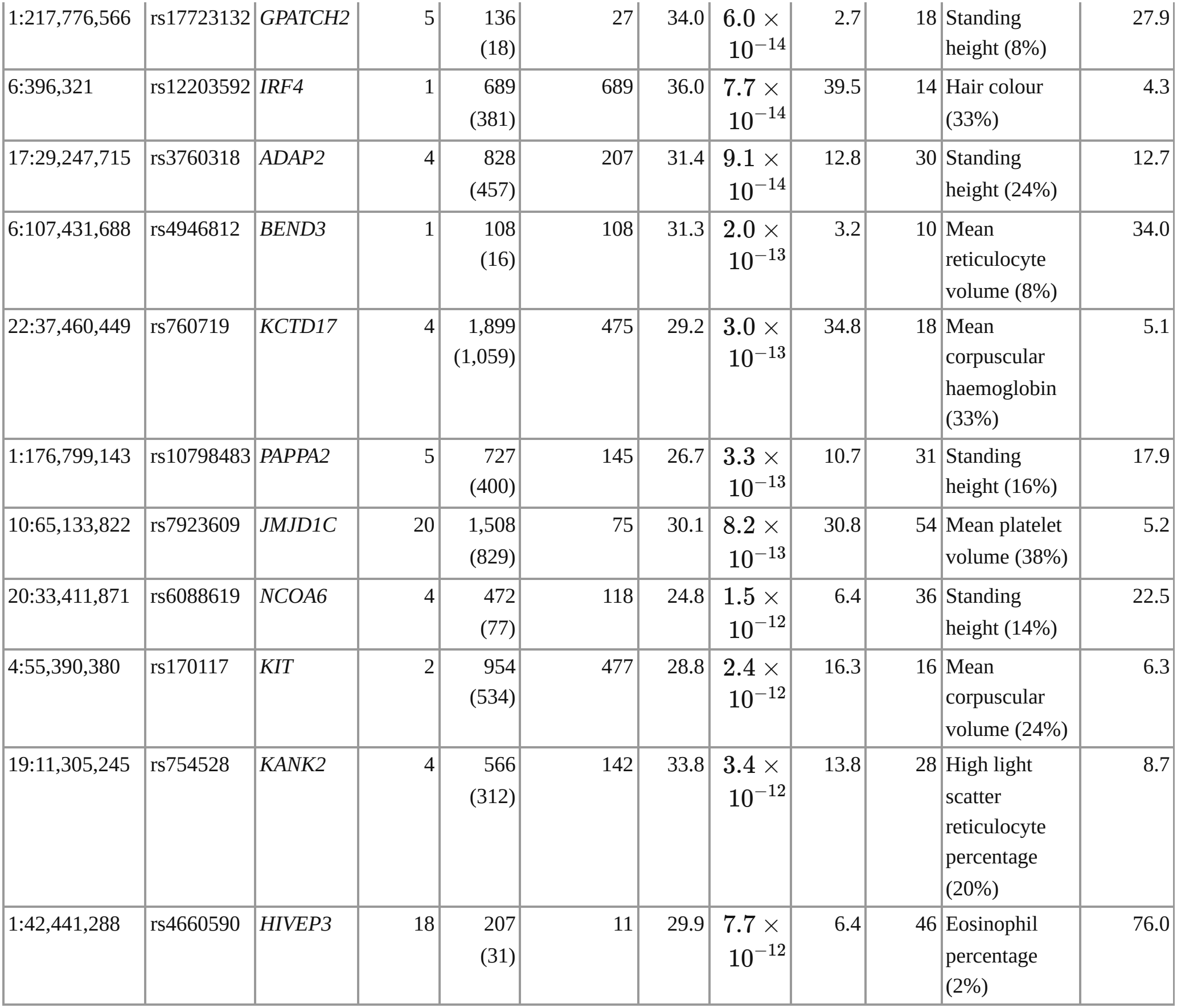
Top omni-loci in Pan-UKB. Best tag per 1-Mb locus, with nearest gene, clump size, *Ŵ* , SE, clumpmean enrichment, *z, q*, mean *χ*^2^ and the number of traits with *P* < 5 × 10^−8^. SE is the model-based standard error of the estimate (background-filled plug-in); *z* and *q* are from the score-type test of 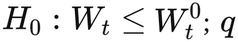 is a tag-level BH value, and values far below the simulated range (about 10^−4^) rest on the moment-matched reference rather than on simulation. Largest trait: the trait contributing the largest share of the lead tag’s pooled response; effective traits: 1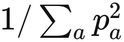 over the positive shares *p*_*a*_. Loci are ordered by significance; GRCh37 positions.

**Figure 5.**
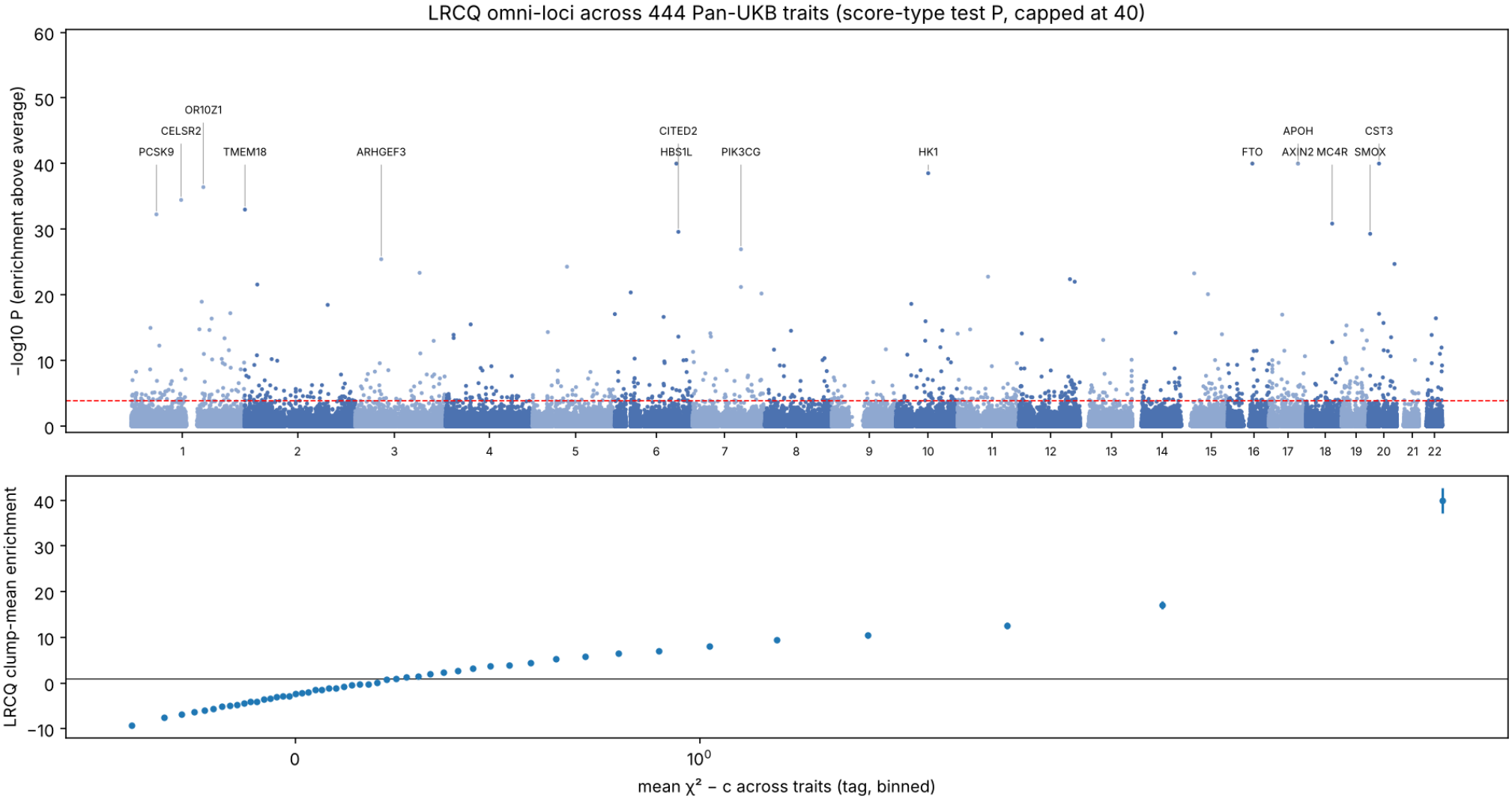
Omni-loci in Pan-UKB. (a) Score-type test − log_10_ *P* for *H*_0_: clump enrichment no greater than its expectation under uniform enrichment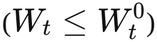 , along the genome (capped at 40); the dashed line is FDR 5%, and the top loci are labelled by nearest protein-coding gene. (b) Mean LRCQ clump enrichment in 50 bins of mean *χ*^2^ across traits.

#### Omni-loci

The most significant loci were well-known pleiotropic regions (Table 2). Grouped by the trait that contributed most to each lead tag, they included:

- FTO, TMEM18 and MC4R (body size and composition); AXIN2, ZBTB38, KDM2A and PAPPA2 (height);
- HBS1L-MYB, CITED2, SMOX, KCTD17 and KIT (red-cell traits), and ARHGEF3, TUBB1, F2R, WDR66 and JMJD1C (platelet traits);
- CELSR2-SORT1, PCSK9, GCKR and ALDH1A2 (lipids);
- CST3 and PRKAG2 (kidney function), HK1 (HbA1c), and HERC2 and IRF4 (hair colour).

Clump-mean enrichment at the lead tags of the 40 top loci had median 209 and reached 2,679, meaning these clumps carry two to three orders of magnitude more phenome-wide heritability per SNP than an average HapMap3 SNP. LRCQ’s ranking agreed only partly with the per-SNP comparators: Spearman 0.54 with mean *χ*^2^, 0.57 with LD-normalised *χ*^2^, 0.22 with counts of significant traits and 0.12 with the omnibus statistic. Several top LRCQ loci had few genome-wide significant traits, CST3 (three) and HERC2 (five) among them. Conversely, of the 2,536 tags with 20 or more genome-wide significant traits, 85% were not significantly enriched above the uniform expectation once LD was accounted for. Their significance counts largely reflect LD with a nearby lead variant. A related question is whether a top locus is enriched across many traits or reflects one very strong association. For the lead tags of the 40 top loci, the largest single trait contributed a median 24% of the pooled response, and the effective number of contributing traits 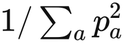, with *p*_*a*_ the positive shares) had median 9.4. FTO, TMEM18 and MC4R were broadly pleiotropic (largest share below 5%, 33 to 40 effective traits). Eight lead tags were dominated by one trait: WDR66, TUBB1, ARHGEF3 and F2R (platelet volume or distribution width, 68-85%), HERC2 (hair colour, 58%), HK1 (HbA1c, 53%), CST3 (glomerular filtration rate from creatinine and cystatin C, 52%) and ITGA4 (monocyte count, 52%). The dominance index is therefore a useful companion to every omni-locus estimate, because the power-weighted estimand cannot by itself tell broad pleiotropy from one very strong association.

#### Significance and the size of the estimate

With the score-type test, significance followed the size of the estimate: 26 of the 30 tags with the largest *Ŵ*_*t*_ (1,254 to 5,013), including FTO, the APOE region, HBS1L-MYB, MC4R, ATXN2-SH2B3, GDF5-UQCC1, HERC2, GCKR and CELSR2-SORT1, were enriched above the uniform expectation at FDR 5%. The four exceptions were tags of the ABO locus, whose enrichment is split between neighbouring tags in strong LD (*z* 2.0-4.8). Their standard errors remain large (for the FTO lead tag, *Ŵ*_*t*_ = 4,396 with SE 2,320): the data show clearly that these clumps are enriched, but with few effective traits they determine by how much only to within a factor of about two. Which traits carry an omni-locus can be checked by deleting one trait at a time. In the TP53 region (Supplementary Fig. S3) the ranking of the 298 core tags was almost unchanged by any single deletion (Spearman correlation with the full ranking at least 0.998 for 440 of the 444 deletions), but the estimate of rs2955617, the strongest tag next to TP53 (*Ŵ*_*t*_ = 639, *q* = 3.8 × 10^−4^), fell to −319 without SHBG: a single trait can carry an omni-locus, and its estimate depends on the trait panel.

#### Gene-level enrichment and constraint

Averaging per-SNP enrichment over HapMap3 SNPs within 10 kb of 18,280 protein-coding genes, phenome-wide enrichment fell steadily with tolerance to loss-of-function variation. Genes in the most constrained gnomAD LOEUF decile had mean enrichment 3.6 (median 1.7), against 1.9 (median 0.4) in the least constrained decile (Spearman *ρ* = −0.16, *P* = 3 × 10^−98^; Fig. 7). This extends to the whole phenome the known concentration of single-trait heritability near constrained genes. It fits the view that variants in dosage-sensitive genes have consequences across many traits.

#### Genes with shared association profiles

We compiled from the GWAS Catalog (Sollis et al. 2023) gene pairs whose association profiles across traits were strikingly similar. For the 46 genes of this compilation found in our gene set, the median LRCQ enrichment ranked at the 80th percentile of all genes. Gene enrichment correlated with each gene’s number of associated traits in the compilation (Spearman 0.56, *P* = 6 × 10^−5^). TP53 had gene-level enrichment 50 (99.6th percentile) and MDM2 was unremarkable (1.1, SE 1.3). One such pair is the TP53 variant rs78378222 and the MDM2 variant rs3730556, whose Z-scores correlate 0.40 across 147 traits of the Common Metabolic Diseases Knowledge Portal PheWAS although the variants are in linkage equilibrium (Chen et al. 2026). The LRCP analysis finds that, in Pan-UKB, their shared signal lies on a narrow axis of body-size and red-cell traits and is significant under the conditional test (bootstrap *P* = 0.015) but not under a class null (*P* = 0.18), so it reports the pair descriptively. The TP53 signal, however, comes mostly from one trait. The TP53 gene window lies about 40 kb from *SHBG*, and at its two most enriched tags sex hormone-binding globulin alone (*z* ≈ −60) contributed 40-60% of the pooled *χ*^2^. The model SE of the TP53 estimate (7.3) assumes signal spread over many traits. The trait-cluster jackknife SE (35), although not valid under overlap, shows how strongly the estimate depends on a single trait cluster. TP53’s place in the phenome-wide map is therefore better described as a SHBG cis-locus with modest further pleiotropy than as evidence for TP53 itself.

#### Functional annotations

We regressed the trait-pooled response on window-wise annotation-stratified LD scores for the 107 baseline-LF v2.2 annotations, fitted jointly. Phenome-wide heritability was concentrated in the following annotations (fold enrichment, block-jackknife SE; Fig. 6):

**Figure 6.**
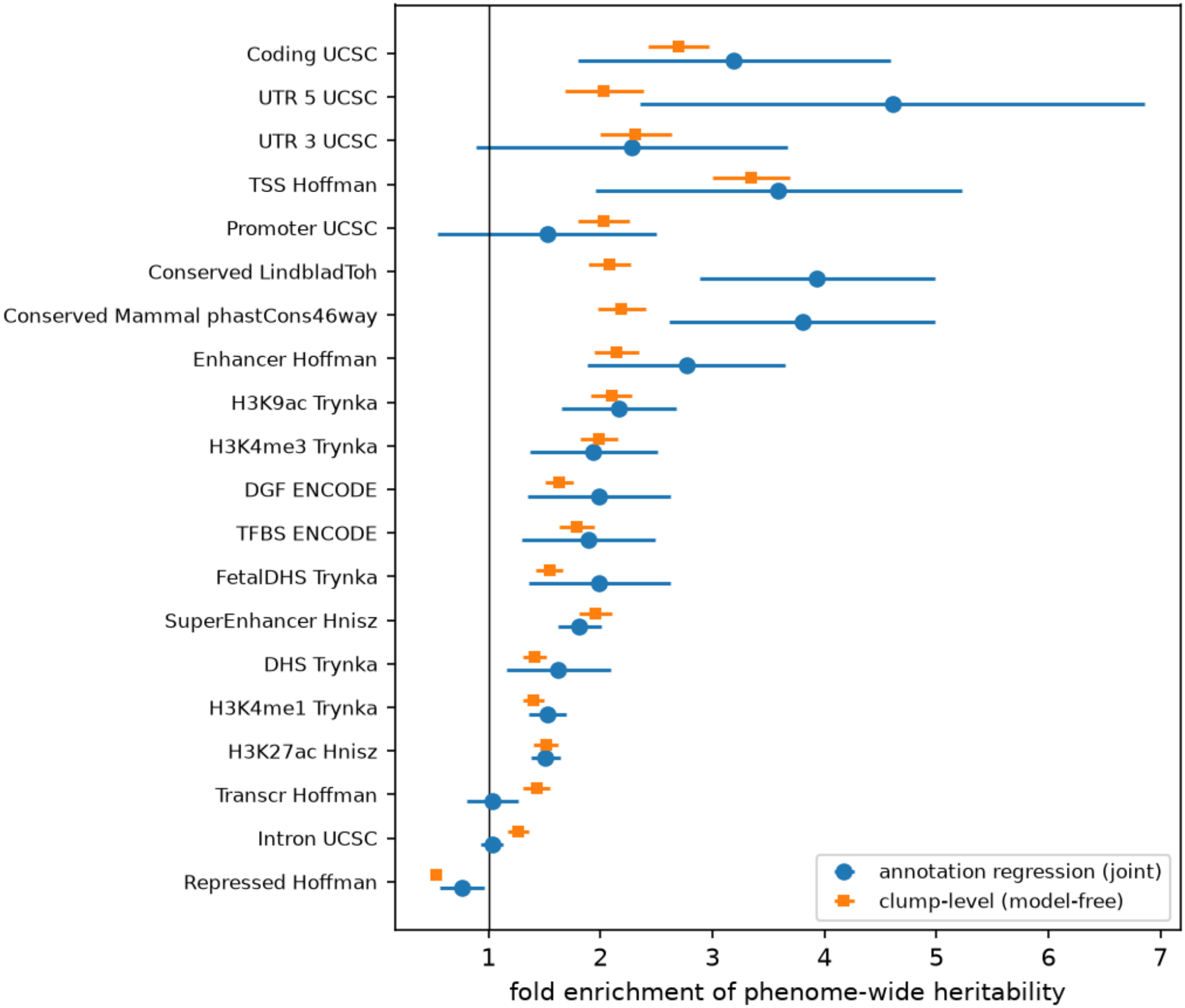
Functional architecture of phenome-wide enrichment. Fold enrichment of phenome-wide heritability in 20 baseline-LF annotations, from the joint annotation regression and the model-free clump-level estimator, with 95% CI (block jackknife).

**Figure 7.**
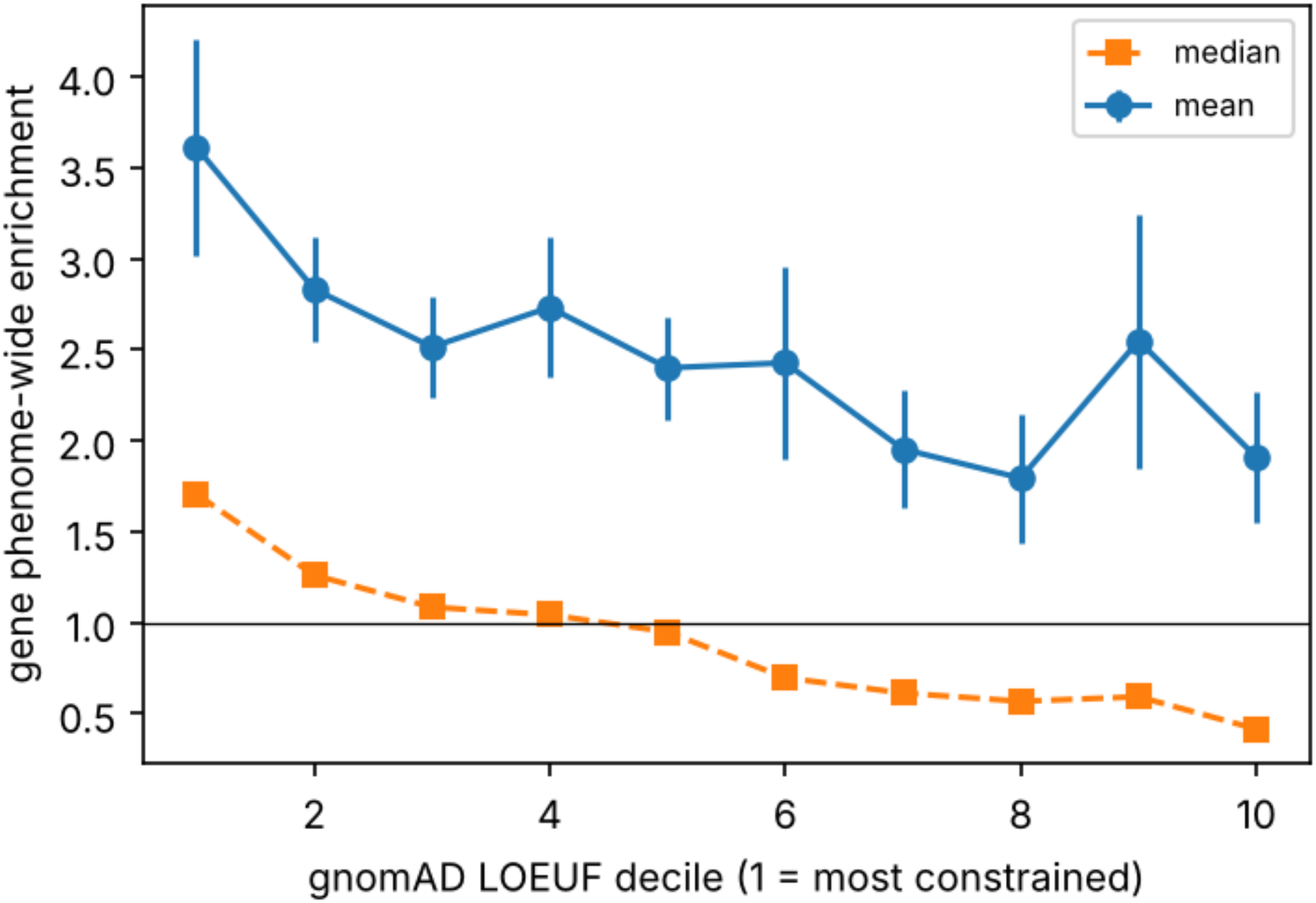
Phenome-wide enrichment and loss-of-function constraint. Mean (95% CI) and median gene-level enrichment by gnomAD v2.1.1 LOEUF decile.

- coding sequence (3.2 ± 0.7);
- 5′ UTRs (4.6 ± 1.2);
- transcription start sites (3.6 ± 0.8);
- conserved sequence (3.9 ± 0.5 for the Lindblad-Toh set); GERP-constrained bases (4.9 ± 1.1);
- active-promoter and enhancer marks (H3K9ac 2.2, H3K4me3 1.9, enhancers 2.8);
- fine-mapped eQTLs (GTEx MaxCPP; 3.7 ± 1.0).

It was depleted in repressed chromatin (0.77 ± 0.10). The model-free clump-level estimates agreed in every case. As simulation predicted, they were attenuated toward 1 because the annotations split LD clumps: coding 2.7, conserved 2.1, TSS 3.4, repressed 0.53, each with SE ≤ 0.2. The pattern resembles meta-analyses of single-trait s-LDSC (Finucane et al. 2015; Gazal et al. 2017). The magnitudes are similar to those in the average single trait, which suggests that the functional architecture of phenome-wide pleiotropy follows that of single-trait heritability rather than being dominated by a distinct class of regulatory hubs.

#### Trait collections and weighting

The estimand depends on which traits are analysed and how they are weighted, so we repeated the analysis in two ways.

- *Pan-UKB’s 150-trait maximally independent set*. Tag estimates correlated 0.65 with the full-set estimates (Spearman 0.55), and 42% of the top 1% of tags were shared.
- *All 444 traits with equal weights*. Correlation was 0.68 (Spearman 0.51), with 51% of the top 1% shared.

Agreement was moderate in both comparisons. The full-set OLS estimate is dominated by the most powerful GWAS, which in Pan-UKB are the many anthropometric and blood-biochemistry traits (Proposition 3.2). The independent set gave a larger spread of true enrichment (method-of-moments SD about 2.5 times that of the full set, computed with the same type of standard error for both), because fewer, more distinct traits make enrichment concentrated in one trait domain look larger. Annotation enrichments, by contrast, replicated closely. In the independent set the estimates were coding 2.7 ± 0.8, conserved 3.4 ± 0.6, TSS 4.5 ± 1.2, 5′ UTR 5.3 ± 1.3 and repressed 0.66 ± 0.12. Equal weighting gave clump-level coding 2.1, TSS 2.7 and repressed 0.72. Aggregate conclusions are robust to the trait collection. Which individual loci rank highest is not, and should be reported together with the trait set.

#### Biomarkers against assessment-centre traits

To ask whether the functional architecture of phenome-wide enrichment differs between trait domains, we contrasted the 51 blood and urine biomarkers with the 252 assessment-centre traits (anthropometry, body composition, blood pressure and lifestyle measures) over 40 randomly chosen non-overlapping windows (47,751 SNPs). Fold enrichments relative to each group’s genome average were:

| **annotation** | **biomarkers** | **assessment centre** |
|---|---|---|
| transcription start sites | 4.4 | 1.7 |
| coding | 3.8 | 2.9 |
| enhancers | 2.7 | 2.0 |
| super-enhancers | 2.5 | 1.7 |
| conserved | 2.2 | 2.9 |
| repressed | 0.22 | 0.64 |

The two groups had similar overall enrichment in these windows (contrast of the base term, bootstrap *P* = 0.39). Biomarker heritability was more concentrated at transcription start sites, enhancers, H3K27ac and super-enhancers, and more depleted in repressed chromatin. With 200 parametric-bootstrap draws, the TSS, H3K27ac, super-enhancer and repressed contrasts reached the bootstrap floor (*P* < 0.005) and the enhancer contrast had *P* = 0.010 (BH *q* < 0.02 over the 10 annotations). Coding, conserved, 5′ UTR and promoter contrasts were not significant (*P* > 0.12). Blood biomarkers are closer to gene expression than the organism-level traits measured at the assessment centre, so heritability concentrated in promoter-proximal regulatory elements fits that difference. The pooled analysis averages these two architectures, weighted by GWAS power.

#### Stage 1-2 inputs

The intercepts are LRCQ’s critical input in simulation, so we recomputed the genome-wide estimates with alternative intercepts, using the exact linear shift of each tag estimate in the intercepts (Methods). Replacing Pan-UKB’s intercepts by our re-estimates, shifting every intercept by ±1 SE, or perturbing each by an independent error with its own SE (20 draws) left the estimates essentially unchanged (Spearman > 0.9998 with the main estimates; 98-100 tags at FDR 5%). Errors of a few hundredths in individual intercepts average out over 444 traits. Only a bias shared by most traits, like the 0.1 in every trait of the misspecified-intercept simulation, would matter.

#### LD reference

Over 20 windows (16,077 SNPs) where we also had UK Biobank in-sample LD, per-SNP clump-mean enrichment from 1000 Genomes and in-sample LD correlated 0.81 (Spearman 0.78), and 250-kb block means correlated 0.95. Mean enrichment differed little (1.58 vs 1.42) and SEs were similar. The choice of reference therefore changes fine-scale assignment within LD clumps, as expected from differences in tagging, but not regional estimates.

#### Why naive replication fails in a single biobank

We tested the motivating observation directly. Traits were split at random into halves, and for each half we computed the LD-score residuals of the pooled per-SNP signal. Across 20 splits the residuals correlated 0.86 at the SNP level and 0.93 for 1-Mb blocks. Under uniform enrichment, however, the expected SNP-level correlation was already 0.91 (Methods), because both halves are measured in largely the same UK Biobank participants and their sampling noise is correlated (Fig. 1). In a single biobank, the visual impression that the same SNPs are “consistently inflated” across traits is therefore dominated by shared samples. Separating true phenome-wide enrichment from that noise needs a model of sample overlap like LRCQ’s. That LRCQ still finds strongly non-uniform enrichment after modelling overlap is the substantive answer to the motivating observation.

#### Replication in independent cohorts

Modelling overlap within one biobank still leaves the question whether the enriched clumps recur in other samples. We therefore reran LRCQ in two cohorts without UK Biobank participants: FinnGen release 12 (Kurki et al. 2023; 57 traits matched to Pan-UKB traits, mostly clinical laboratory measurements, with height, body-mass index, weight and 19 disease endpoints) and BioBank Japan (Sakaue et al. 2021; 39 traits). Each cohort had its own Stage 1-2 estimates and an ancestry-matched 1000 Genomes LD reference, and each discovery tag was looked up through the tag of its clump in the replication run (Methods). One-sided score-type p-values for the discovery tags were:

|  | **FinnGen R12** | **BioBank Japan** |
|---|---|---|
| Matched traits (median $n$) | 57 (376,471) | 39 (145,502) |
| Tags with $W_t > 0$ in discovery (1,799): testable | 1,765 | 1,526 |
| $\hat{W}_t > 0$ | 1,494 (85%) | 1,093 (72%) |
| $P < 0.05$ for $W_t > 0$ | 954 (54%) | 535 (35%) |
| BH 5% within the testable tags | 826 | 356 |
| Other tags of the run with $P < 0.05$ | 7.6% | 6.6% |
| Spearman correlation with discovery $\hat{W}_t$ | 0.53 | 0.37 |
| Tags above the uniform expectation in discovery (881): testable | 860 | 728 |
| $P < 0.05$ above the uniform expectation | 497 (58%) | 282 (39%) |
| BH 5% within the testable tags | 458 | 195 |

Discovery tags were seven times as likely as the other tags of the FinnGen run to have *P* < 0.05, and five times in BioBank Japan (Supplementary Fig. S4; Supplementary Table S9). The trait panels are small, so a complete replication is not expected. As a reference, the same analysis of Pan-UKB restricted to the matched traits, which reuses the discovery participants, gives *P* < 0.05 for 1,523 (FinnGen traits) and 1,548 (BioBank Japan traits) of the 1,799 tags; FinnGen thus recovers about 60% of what its trait panel can show in UK Biobank. Replication across ancestries was weaker but clearly above chance, as expected from the smaller sample, fewer traits, the 14% of HapMap3 SNPs missing in BioBank Japan and differences in LD and allele frequency. Rank agreement with the discovery estimates was moderate, in line with the dependence of the estimand on the trait panel (above). Phenome-wide enrichment found in Pan-UKB is therefore largely reproducible in independent samples, including a non-European population.

## Discussion

We have introduced LRCQ, which estimates how much each locus contributes to genetic variance across a collection of traits, relative to an average locus. It does this from GWAS summary statistics and an LD reference, and it separates true pleiotropic effects from LD tagging and from GWAS-wide confounding. LRCQ extends the LD score regression family along a new axis. LDSC assumes every SNP has the same expected effect variance. Stratified LDSC lets that variance depend on annotations. LRCQ lets it vary freely across the genome and identifies it by pooling traits instead of imposing annotations. The resulting estimates can be aggregated over any set of SNPs chosen after the analysis (functional annotations, genes, pathways, trait categories) without refitting a model, and with model-based standard errors that account for sample overlap and genetic correlation between traits.

Three properties of the problem limit what LRCQ can deliver.

First, resolution. Squared Z-scores identify phenome-wide enrichment only at the resolution of LD clumps. Among SNPs in tight LD, the split of enrichment is not identified: the LD matrix that must be inverted is numerically singular at HapMap3 density (Supplementary Note §S3.11). LRCQ therefore reports clump-level estimates, and an “omni-SNP” is more precisely an omni-locus. Which variant in a clump carries the enrichment is a fine-mapping question that needs functional priors or individual-level data. LRCQ’s tag estimates could serve as a cross-trait prior for that step, for example in PolyFun-style fine-mapping (Weissbrod et al. 2020).

Second, precision. Even with hundreds of biobank traits, the effective number of independent traits is far smaller than their count. Sample overlap and genetic correlation reduce it, and the two effects enter the variance separately (Supplementary Note §S3.10, §S9). With UK Biobank-sized GWAS and about 100 effective traits, detecting a single locus at genome-wide significance needs enrichment of about 20-30 or more (Supplementary Note §S9.2). Near the null, doubling the GWAS sample size is worth quadrupling the effective number of traits. Simulations in which the per-SNP signal was large instead found that the number of GWAS mattered more than their size. The two results agree: once sampling noise is small relative to the spread of a SNP’s effects across traits, only more traits reduce the remaining variance. Aggregated quantities, such as annotation enrichments over thousands of clumps, are estimated precisely. This mirrors the history of heritability partitioning: per-SNP heritability is noisy, but its sums over annotations transformed the interpretation of GWAS. Adding genetically distinct trait domains increases power more than adding more traits from domains already represented.

Third, the estimand depends on the trait collection and its weighting. *w* is defined relative to the traits analysed. The efficient (OLS/WLS) estimator weights each trait by 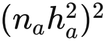, so large, highly heritable GWAS dominate (Proposition 3.2). We report the equal-weight estimator and category-specific estimates alongside. In Pan-UKB, where anthropometric and blood-biochemistry traits are numerous and well powered, the power-weighted and equal-weight maps correlated only 0.68, and the 150-trait independent set agreed with the full set at 0.65. Annotation-level enrichments agreed closely across all three. A locus that is an omni-locus for one trait collection need not be one for another. Phenome-wide enrichment is a property of a variant together with a phenome, much as heritability is a property of a trait together with a population.

### Relation to genetic-effect correlation

LRCQ’s working assumption is that genetic effects of SNPs that share an LD partner are uncorrelated across traits. When this is violated, the *χ*^2^-only estimator absorbs the correlation into apparent enrichment near the correlated pair, with an exactly known bias (Theorem 2). Correlations between SNPs in different LD blocks, which LRCP targets, cause no bias. Squares and cross-products together identify the full SNP covariance within a block (Supplementary Note §S3.9). A joint analysis is a natural extension. In a 100-SNP simulation, adding within-GWAS cross-products of SNP pairs in LD (*r*^2^ > 0.05) raised the adjusted *R*^2^ with the truth from 0.86 to 0.93. The exact joint estimator, however, needs **R**^−1^, which is ill-conditioned at HapMap3 density (Supplementary Note §S3.9), so a regularised joint estimator is the practical next step.

#### Confounding

Stratification and cryptic relatedness enter through the per-trait LDSC intercepts, which we fix at genome-wide estimates. They cannot be re-estimated per window without losing identifiability (Proposition 3.3). If the true intercepts exceed the assumed ones by *δ*_*a*_, the estimates shift by (∑_*a*_ *s*_*a*_*δ*_*a*_/∥**s**∥^2^) **D**^−1^**1**, which is largest for low-LD SNPs. In simulation, an intercept of 1.1 wrongly set to 1 biased every tag upward by 0.7 and doubled the type-I error. LRCQ inherits LDSC’s sensitivity to intercept estimation, including the attenuation-ratio issue in very large biobank GWAS (Loh et al. 2018). LD reference mismatch is a second source of error. In-sample UK Biobank LD removes most of it for Pan-UKB, and in simulation, references of about 500 individuals lowered the correlation with the truth only from 0.98 to 0.96, but inflated the spread of standardised errors to 1.5. Our Pan-UKB analysis uses 1000 Genomes LD. Over 20 windows where in-sample LD was available, per-SNP estimates correlated 0.81 between the two references and regional (250-kb) means 0.95, but the SEs from 1000 Genomes LD are probably optimistic in the way simulation showed.

#### Estimator choice and inference

Three simulation results differ from what one might expect. First, one-step WLS, which is efficient in theory, was slightly attenuated and less accurate than OLS. Its weights are estimated from the same Z-scores as its responses, and Gaussian variance weights are wrong near causal variants with sparse effects (Supplementary Note §S5). Second, model-based standard errors depend on which estimate of **w** is plugged into them. Keeping only the estimates that survive rectification drops the polygenic background and made SEs 10-25% too small, whereas filling the rectified SNPs with the background mean calibrated them in every correctly specified scenario. A jackknife over trait clusters, the usual distribution-free remedy, failed when traits shared samples: overlapping GWAS are never independent clusters, and its SEs were less than half the true SD. Third, with sample overlap the estimates are strongly right-skewed, so normal-reference p-values bend a QQ plot below and then above the diagonal and overstate the significance of the strongest loci. In Pan-UKB they would have declared 5,178 tags in 1,961 loci instead of 881 tags in 638 loci. The exact third cumulant is cheap because the covariance of the Z-scores is a sum of two Kronecker products, and a skewness-matched reference restored tail calibration down to *P* = 10^−3^ in every simulated scenario with correlated or few traits. Further into the tail the excess kurtosis matters too, and a reference matched to the exact fourth cumulant was needed (Supplementary Table S7). Tests use score-type variances, with the tested tag’s plug-in set to its null value, because a variance evaluated at a large estimate grows with that estimate and leaves almost no power for the most enriched loci. We therefore recommend OLS with model-based SEs, score-type tests with moment-matched p-values, and the background-filled plug-in of **w**, with each estimate reported together with the share of the signal from the single largest trait. That index exposed eight of the 40 top loci, among them WDR66, TUBB1 and ARHGEF3 (platelet volume), whose estimates rested mostly on one trait. A parametric bootstrap that draws Z-scores with the estimated cross-trait covariance and a non-Gaussian effect distribution is the natural remedy. Category contrasts need a parametric bootstrap, because plug-in variances from the same data are conservative (Supplementary Note §S7.4a).

### Limitations

- The model assumes a common SNP covariance pattern across traits up to scale (assumption A2). When enrichment differs between trait groups, LRCQ estimates a weighted average, which we partly address with category-specific runs.
- Inference uses Gaussian moments up to the fourth, and the p-values match three or four cumulants rather than the full distribution. Under the Pan-UKB trait structure the kurtosis-matched p-values were calibrated at *P* = 0.01 but 1.3-to 1.6-fold anti-conservative between 10^−3^ and 10^−4^ (Supplementary Table S7), so the FDR at the 5% threshold may be somewhat above nominal; doubling every p-value still left 514 omni-loci. Model SEs need a plug-in estimate of **w** from the same data; the background-filled plug-in was calibrated in simulation but slightly conservative with the Pan-UKB trait structure, and plug-in variances from the same data are conservative for contrasts, which we therefore test by parametric bootstrap. A trait-cluster jackknife is not a valid alternative when GWAS share samples.
- Significance is tested with score-type variances, but each estimate is reported with its model-based standard error at the plug-in estimate, which is the right measure of its precision. A locus can therefore be highly significant and still have a wide confidence interval: with few effective traits, the data show clearly that a clump is enriched but determine its enrichment only to within a factor of about two (Table 2). Ranking omni-loci by *z* and by *Ŵ*_*t*_ answers different questions.
- The analysis is restricted to HapMap3 SNPs with MAF > 1% in European-ancestry UK Biobank participants. Rare variants and other ancestries are not addressed. Enrichment is defined relative to HapMap3-tagged heritability.
- Binary traits enter on the observed scale. Only 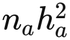 matters, so switching to effective sample sizes or the liability scale changes nothing if 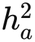 is re-estimated on the same footing. What matters is that *n*_*a*_ and 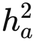 come from the same LDSC fit.

#### Outlook

LRCQ provides a genome-wide, LD-corrected map of where phenome-wide genetic variance is concentrated. With its companion LRCP, it gives a summary-statistic description of the SNP-side covariance of the genetic effect matrix: how much each locus matters across the phenome, and which loci act together. Natural next steps are annotation-informed priors for within-clump resolution (for example, a classifier trained on functional annotations, with strongly negative and strongly positive *Ŵ* as labels), multi-ancestry LD for finer resolution, and using phenome-wide enrichment as a prior for cross-trait polygenic prediction.

## Methods

### Model and estimand

We consider *q* traits measured in individuals with standardised genotypes at *m* SNPs, **Y** = **XB** + **E**. The *m* × *q* matrix of joint (causal) effects **B** is random with mean zero and separable second moments,

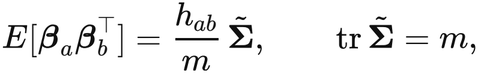

where ***β***_*a*_ is column *a* of **B**, *h*_*ab*_ is the SNP co-heritability of traits *a* and 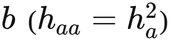, and 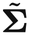 is a normalised SNP-by-SNP covariance shared by all traits up to scale (Supplementary Note §S1.3). This is the second-moment structure of a matrix-normal model, 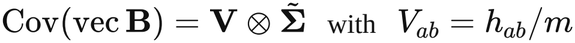 with *V*_*ab*_ = *h*_*ab*_/*m* (Supplementary Note §S2.1). Normality is not needed for the moment identities below.

The **phenome-wide heritability enrichment** of SNP *s* is 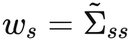 , so avg(**w**) = 1. Equivalently, *w*_*s*_ = diag(**Σ**_*β*_)_*s*_/ avg diag(**Σ**_*β*_) with **Σ**_*β*_ = *E*[**BB**^*⊤*^]/*q*: the variance of the SNP’s causal effect across the traits analysed, relative to an average SNP. A SNP with *w*_*s*_ = 0 affects none of the traits (“null SNP”). A SNP with *w*_*s*_ *≫* 1 carries a disproportionate share of heritability across many traits (“omni-SNP”, or omni-locus at LD-clump resolution; see below). Unlike per-SNP heritability enrichment from stratified LD score regression (s-LDSC; Finucane et al. 2015), *w* is defined across the phenome and needs no functional annotation. The off-diagonal part of 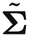 olds the genetic-effect correlations *ρ*_*ij*_, the target of the companion method LRCP.

### Summary-statistic moment equation

GWAS Z-scores follow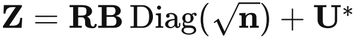, where **R** is the LD matrix and the sampling errors satisfy 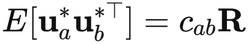. Here *c*_*aa*_ is trait *a*’s LD score regression intercept, which absorbs population stratification and cryptic relatedness, and 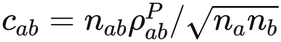 reflects sample overlap and phenotypic correlation (Lemmas 1.1-1.2). Then for SNP *k* and trait *a* (Theorem 1, Corollary 1),

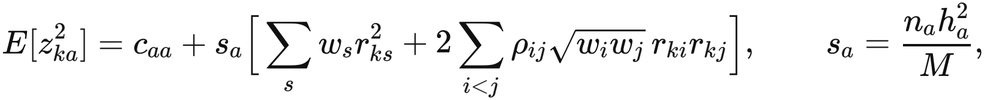

with *M* the genome-wide number of SNPs over which 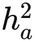 is defined. With **w** ≡ **1** and *ρ* ≡ 0 this is LD score regression (Bulik-Sullivan et al. 2015a). With **w** in the span of annotations it is s-LDSC. LRCQ is the case of one free parameter per SNP, made estimable by pooling many traits.

From squared Z-scores alone only **Dw** + diag(**RCR**) is identified, where **D** = **R** *∘* **R** and **C** is the off-diagonal part of 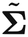 (Proposition 1.6). LRCQ therefore estimates **w** under the working assumption that genetic-effect correlations between SNPs sharing an LD partner are zero. When this fails, the bias of any linear unbiased LRCQ estimator is exactly **D**^−1^ diag(**RCR**) (Theorem 2). The bias is zero for correlations between SNPs in different LD blocks, and positive near correlated SNPs in LD. We quantify it by simulation.

### Estimation

LRCQ is a stage-wise plug-in estimator (Supplementary Note §S0). It proceeds in stages: trait heritabilities and cross-trait terms first, under **w** ≡ **1** and *ρ* ≡ 0; then **w** with *ρ* = 0; then, in LRCP, SNP pairs screened by **Ŵ** before *ρ* is estimated. Table 3 lists the unknowns of each stage, how they are estimated and where the Pan-UKB application stands.

**Table 3.** Unknown parameters of the LRCQ-LRCP model, how each is estimated, and status in Pan-UKB. Inputs are the Z-score matrix **Z** (*m* × *q*; *m* = 1,094,844 HapMap3 SNPs, *q* = 444 Pan-UKB EUR traits), sample sizes *n*_*a*_ and LD **R** (1000 Genomes EUR in 1,383 3-Mb windows; UK Biobank in-sample LD as a sensitivity analysis). Each stage holds the previous stages’ estimates fixed. Stage 4 is the companion method LRCP.

| Stage | Parameter | Number of unknowns | Estimator (default) | Identification and assumptions | Inference | Pan-UKB |
|---|---|---|---|---|---|---|
| 1 | Heritability $h_a^2$ and genetic covariance $h_{ab}$ | $q(q+1)/2$ (98,790) | Univariate and cross-trait LDSC, genome-wide; genetic covariances rescaled to the Pan-UKB $h^2$ and projected to PSD | Fitted under $\mathbf{w} \equiv \mathbf{1}$ , $\rho \equiv 0$. A common misscaling of $h^2$ only rescales $\hat{\mathbf{w}}$, which is renormalised to mean 1 (Supplementary Note §S1.7) | Plug-in | $h^2$ and intercepts agree with Pan-UKB LDSC ($r$ = 0.992, 0.985) |
| 2 | Intercepts $c_{aa}$ (stratification) and $c_{ab} = n_{ab}\rho_{ab}^P/\sqrt{n_a n_b}$ (sample overlap × phenotypic correlation) | $q(q+1)/2$ | LDSC intercepts, genome-wide | Held fixed. Free per-window intercepts leave $\mathbf{w}$ unidentified (Proposition 3.3) | Plug-in; enter every later variance through $\boldsymbol{\Gamma} = (c_{ab})$ | Median SE of a singleton tag 14.5 under the 444 traits’ overlap and genetic correlation |
| 3 | Phenome-wide heritability enrichment $\mathbf{w}$ | $m$ in principle; identified only as LD-projected tag enrichments $W_t$, one per tag (clump totals for tight clumps) | Tag-set LRCQ: closed-form OLS in 3-Mb sliding windows (2-Mb cores), unrectified; WLS, equal-weight and per-SNP ridge as sensitivity analyses | $\rho = 0$ between SNPs sharing LD partners (otherwise bias $\mathbf{D}^{-1}\,\mathrm{diag}(\mathbf{RCR})$ , Theorem 2); tags pruned at $r^2 < 0.5$; second moments only | Exact cross-trait variance (Theorem 3.10); background-filled plug-in of $\hat{\mathbf{w}}$ (Supplementary Note §S3.10a); one-sided score-type tests with moment-matched (three- or four-cumulant) p-values, BH FDR | 301,693 tags; 881 tags at tag-level FDR 5% (638 loci); 1,799 tags with $W_t > 0$ |
| 3′ | Aggregates of $\mathbf{w}$ : annotation enrichment $E_c$ and $\boldsymbol{\tau}$, gene enrichment $E_G$, phenotype-category contrasts | One per annotation, gene or contrast | $\mathbf{D}^2$-weighted annotation regression (primary); clump-level $E_c$ (check); LRCQ per trait category | $\mathbf{w}$ flat within clumps under the null | Block jackknife (200 blocks); parametric bootstrap for contrasts | Enrichment at coding, conserved, TSS and 5′ UTR SNPs; biomarker vs assessment-centre contrast |
| 4 | Genetic-effect correlation $\rho_{ij}$ (off-diagonal of $\mathbf{R}_\beta$; LRCP) | $m(m-1)/2$ in principle; $K(K-1)/2$ among prescreened tags | Closed-form GLS LD-deconvolution of Z cross-products for distal pairs (Gaussian MLE had lower RMSE at $q$ = 30 but under-covering SEs, so GLS is the default); windowed regression or Gaussian MLE for local pairs | Prescreen on LRCQ (no variance, no covariance); each unordered pair counted once | Conditional test, exact under any dependence among GWAS | LRCP primary $K$ = 1,799 (score-type $W_t > 0$, $q < 0.05$; 868 windows, 1.61 M distal pairs; 437 pairs at FDR 5%, 75 Bonferroni, 14 modules); TP53 × MDM2 descriptive only (class-null $P$ = 0.18; Chen et al. 2026) |

#### Stage 1 (heritability)

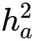 for each trait from LD score regression on the same SNPs. In the Pan-UKB application we use the Pan-UKB LDSC estimates; our own re-estimation agreed closely (below). A common misscaling of *h*^2^ rescales **ŵ** by a constant that is removed by renormalising **ŵ** to mean 1 (Supplementary Note §S1.7). We report renormalised estimates.

#### Stage 2 (intercepts)

*c*_*aa*_ from univariate LDSC and *c*_*ab*_ from cross-trait LDSC intercepts, both genome-wide. They are held fixed in Stage 3: with free per-window intercepts the design has a rank deficit of one and **w** is not identified (Proposition 3.3).

#### Stage 3 (enrichment)

Let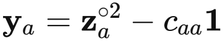, so that *E***y**_*a*_ = *s*_*a*_**Dw**. Stacking traits gives a regression with design **s** *⊗* **D**. Its OLS solution collapses to a single *m* × *m* solve, **ŵ** = ∥**s**∥^−2^**D**^−1^ ∑_*a*_ *s*_*a*_**y**_*a*_ (Theorem 3.1). The closed form replaces the stacked *qm*-row regression, so that only **D** has to be inverted, once per LD reference. The WLS solution with weights 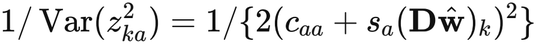 needs only 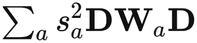 (Supplementary Note §S3.5). In smaller simulations with Gaussian effects in blocks of 20-500 SNPs, OLS, WLS, iteratively reweighted least squares and GLS gave nearly identical estimates; WLS gained slightly only when GWAS sample sizes varied, and maximum likelihood failed when some *w* were zero.

In our simulations with sparse, heavy-tailed effects, one-step WLS was slightly attenuated (slope 0.93-0.97) and less accurate than OLS: its weights are estimated from the same data, and Gaussian weights are wrong near causal variants with heavy-tailed effects (Supplementary Note §S5). We therefore use OLS as the primary estimator, with WLS as a sensitivity analysis. Estimation runs in sliding windows of LD blocks, keeping estimates only for SNPs in the window core (Supplementary Note §S3.7). In the application, windows are 3 Mb long and spaced 2 Mb apart (the layout of the UK Biobank LD release), with LD from 1000 Genomes (below), and cores are their central 2 Mb.

#### Trait weighting

OLS estimates the 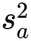 -weighted average of trait-specific enrichments, 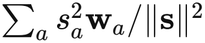 , so large, highly heritable GWAS dominate (Proposition 3.2). WLS weights traits differently again, with weights that also depend on each SNP’s LD score and the current estimate (Supplementary Note §S3.2). We also report the equal-weight estimator 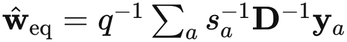, which is unbiased for the simple average across traits but noisier, and category-specific estimates from subsets of traits.

#### Resolution under LD (tag-set LRCQ)

At HapMap3 density in UK Biobank LD, **D** is numerically singular. In a representative 3 Mb window (1,106 SNPs) its condition number was about 5 × 10^13^, and the median (**D**^−1^)_*ii*_, which scales the variance of *ŵ* _*i*_, was 13 (90th percentile 364). Only totals of *w* within groups of tightly linked SNPs are identified (Supplementary Note §S3.11). Our primary estimator is therefore defined on a tag set *T* , obtained by greedy LD pruning at *r*^2^ < 0.5 in order of decreasing LD score. Responses are taken from all SNPs:

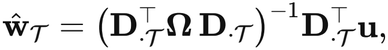

with **Ω** and **u** the trait-pooled weights and responses (**Ω** = ∥**s**∥^2^**I** and **u** = ∑_*a*_ *s*_*a*_**y**_*a*_ for OLS). Its estimand, 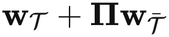 , assigns each untagged SNP’s enrichment to the tags that best reproduce its *r*^2^ profile, and equals the clump total when an untagged SNP is in near-perfect LD with one tag (Supplementary Note §S3.11, eq. 3.6). In general it is a signed LD projection, the LD-projected tag enrichment: allocation weights need not be non-negative or sum to 1, and the null value 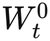 under uniform enrichment can be negative for a tag whose profile is partly explained by its neighbours. After pruning at *r*^2^ < 0.5 the median 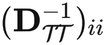 was 1.4. Per-SNP ridge LRCQ, 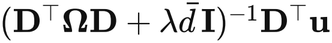 with *λ* = 10^−3^ and 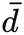 the mean diagonal, is a sensitivity analysis. It spreads each clump’s enrichment evenly over its proxies. Before any solve, *r*^2^ is bias-corrected for the reference size, 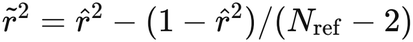, and the LD matrix is projected to the positive-definite cone (eigenvalue floor 10^−4^, rescaled to unit diagonal). We refer to tags with large *ŵ* as omniloci.

#### Negative estimates

Unrectified estimates are used for all inference and aggregation. For reporting a non-negative map we apply the rectifications of Supplementary Note §S3.8. Method C, which zeroes all estimates below the threshold at which the positive mass below it equals the total negative mass, is the unique threshold rule that preserves ∑_*i*_ *ŵ* _*i*_ and hence the mean-1 normalisation (Supplementary Note §S3.8). Methods A and B are compared in simulation, together with the knockoff-style FDR threshold (Method D) proposed in Supplementary Note §S3.8.

### Inference

Traits are not independent replicates. Shared samples and genetic correlation make Z-scores correlated across GWAS, and the effective number of traits can be an order of magnitude below *q* (Supplementary Note §S3.10). We use the exact variance of LRCQ’s quadratic-form estimators under Gaussian **Z** (Theorem 3.10):

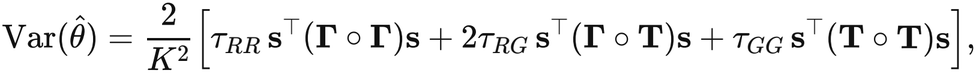

where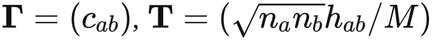, and the *τ* terms are traces that depend on the SNP (or set) being estimated and the LD. The same formula with general trait weights gives variances for equal-weight estimates, annotation enrichments and phenotype-category contrasts (Supplementary Note §S7.1). Gaussian fourth moments understate the variance of *z*^2^ near causal variants when effects are sparse (Supplementary Note §S5).

The traces *τ*_*RG*_ and *τ*_*GG*_ involve **R** diag(**w**)**R**, so **w** must be plugged in from the same data. We use a background-filled estimate (Supplementary Note §S3.10a). Within each window, estimates larger than | min_*i*_ *ŵ*_*i*_| are kept, as in rectification method A (Supplementary Note §S3.8), and every SNP that method A would set to zero receives the mean of *ŵ* over that set, floored at 0. Method A alone removes the polygenic background and made SEs 10-25% too small in simulation, most under dense enrichment, whereas per-SNP clipping max(*ŵ* _*i*_, 0) overstates them because sampling noise enters the variance quadratically. The same plug-in enters the higher cumulants below. For standard errors this plug-in is used as it stands; for tests, the tested tag’s own entry is replaced by its null value (below), because a large estimate kept in the plug-in enters its own variance. We also evaluated a delete-one-cluster jackknife over traits (clusters from average-linkage clustering of 1 − |*r*_*g*_|). With independent traits it was usable at 50 clusters (SD of standardised errors 1.2), but with UK Biobank-like sample overlap and genetic correlation its SEs were less than half the true SD (standardised-error SD 2.3-2.7), because trait clusters are not independent. We use model-based SEs throughout and report the jackknife only as a diagnostic. Independence-based standard errors are never used.

#### Score-type tests and p-values

The OLS tag estimate is a quadratic form in the Z-scores, 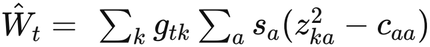, where **g**_*t*_ is row *t* of 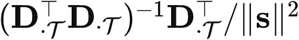. Under Gaussian **Z** with covariance **Σ** = **Γ** ⊗ **R** + **T** ⊗ **A, A** = **R** diag(**w**)**R**, its cumulants are *κ*_*r*_ = 2^*r*−1^(*r* − 1)! tr{(**M**_*t*_**Σ**)^*r*^} with **M**_*t*_ = diag(**s**) ⊗ diag(**g**_*t*_). *κ*_2_ is the variance above. Because both terms of **Σ** are Kronecker products, *κ*_3_ factorises into four trait-side traces, such as tr{(**SΓ**)^3^} with **S** = diag(**s**), computed once, and four SNP-side traces per tag, such as tr{(diag(**g**_*t*_)**R**)^3^}. For the tests of 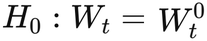 and *H*_0_ : *W*_*t*_ = 0 (below) the cumulants are evaluated at the plug-in with the entries of tag *t* set to their null value: *ŵ* of the tag’s SNPs is replaced by the value that gives *W*_*t*_ its null value, and every other SNP keeps the background-filled plug-in. The null configuration must itself be a valid model: its implied mean for tag 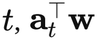 with **a**_*t*_ row *t* of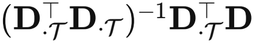 , must equal the null value, and **w** must be nonnegative so that **A** is positive semidefinite. Setting the tag’s entries to a nonnegative null value meets both conditions. Because the projection can assign negative shares, 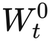 (below) is negative for 5.0% of Pan-UKB tags; for these the null configuration is the nonnegative **w** closest to the background-filled plug-in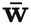 with 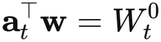, which is 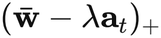 with *λ* found by root-finding. The mean constraint uses **D**, the *r*^2^ corrected for the finite reference panel, because that is the mean of the estimator. The uncorrected **R** ∘ **R** adds about 1/(*N*_ref_ − 2) for every unlinked pair (1/501 here) and would treat reference-panel noise as LD; a constraint stated with it is reported as a sensitivity analysis (Supplementary Table S8). This is a score-type test, since the null distribution is computed under the null. A variance evaluated at the tag’s own estimate would grow almost in proportion to *Ŵ*_*t*_ when the estimate is large, leaving a test with almost no power for the most enriched clumps (Supplementary Note §S3.10a). That variance is still the right measure of how precisely a large *W*_*t*_ is estimated, because with few effective traits *Ŵ*_*t*_ behaves like a variance estimated on few degrees of freedom; we report it with every estimate and use the score-type variance only for testing. When the effective number of traits is small, as under extensive sample overlap, the null distribution of *Ŵ*_*t*_ is strongly skewed and heavy-tailed. A normal reference is then conservative for most tags but anti-conservative in the upper tail. We therefore match moments. With *κ*_4_ = 48 tr{(**M**_*t*_**Σ**)^4^} expanded in the same way into six trait-side and six SNP-side traces, the p-value is the upper tail of a central 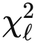 matched to the variance and excess kurtosis, 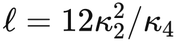 and 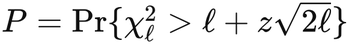 with *z* the standardised distance of *Ŵ*_*t*_ from its null value, the small-sample approximation used for quadratic-form tests in sequencing studies (Lee et al. 2012). The shifted gamma distribution with the first three cumulants (a Pearson type III approximation) matches skewness instead; in simulation it was anti-conservative in the far tail because the excess kurtosis of *Ŵ*_*t*_ (median 4.6) exceeds that of a gamma with the same skewness (median skewness 1.4, gamma kurtosis 1.5*γ*^2^ ≈ 2.9). In the genome-wide analysis, *κ*_3_ is computed for every tag with score *z* > 1.5 and for a random 5% of tags used to check calibration, with SNP-side traces summed over the entries of **g**_*t*_ larger than 3 × 10^−3^ of its maximum (10^−3^ for *z* > 3); these tags get the gamma p-value. *κ*_4_, at tolerance 10^−3^, is computed for every tag with score *z* > 2 under either null (18,514 tags), and these tags get the kurtosis-matched p-value. The remaining tags, all with score *P* > 0.07, keep the normal p-value. Because each tag estimate is linear in the intercepts, ∂*Ŵ*_*t*_/∂*c*_*aa*_ = −*s*_*a*_ ∑_*k*_ *g*_*tk*_, its sensitivity to the Stage 2 intercepts can be evaluated exactly without refitting.

#### Omni-locus tests

For tag *t* we test *H*_0_ : *W*_*t*_ ≤ *W* ^0^ (clump no more enriched per SNP than average) and *H*_0_ : *W*_*t*_ ≤ 0. Here 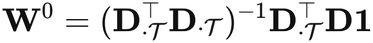 , renormalised like **Ŵ** , is the tag estimand under uniform enrichment **w** ≡ **1** (Supplementary Note §S3.10a). It equals the clump size *g*_*t*_ only when the tags span their clumps’ LD exactly; in Pan-UKB its median ratio to *g*_*t*_ was 1.07 (10th-90th percentile 0.45-2.15). Both tests are one-sided score-type tests with the p-values above and Benjamini-Hochberg FDR (Benjamini & Hochberg 1995) across tags. Both nulls are composite: tags whose true enrichment lies below the boundary give p-values stochastically larger than uniform, so a QQ plot over all tags is expected to fall below the diagonal in its bulk even when the test is calibrated at the boundary. The knockoff-style threshold (Method D) gives a second FDR-controlling discovery set. Under *w*_*i*_ = 0 and without LD, SE 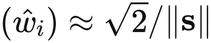 (Supplementary Note §S3.6). For the 444 Pan-UKB traits, sample overlap and genetic correlation raise the median SE of a singleton tag to 14.5, so only strongly enriched loci are individually detectable. Most of the biological inference therefore comes from aggregation.

#### Annotation enrichment

Functional annotations usually split LD clumps, and how *w* divides within a clump is not identified from Z-scores and LD alone (Supplementary Note §S7.2a). Annotation enrichment is estimated in two ways.

The primary analysis is the **D**^2^-weighted annotation regression 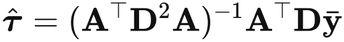 (Supplementary Note §S7.3). It needs no **D**^−1^, borrows strength across clumps, and recovers the fold-enrichment 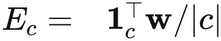 when **w** lies in the span of the annotations. This is the identification used by s-LDSC, applied here to the trait-pooled response.

The model-free check is the clump-level enrichment 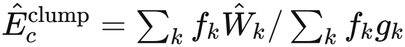 . Here *Ŵ* _*k*_ is the tag-set estimate for clump *k, g*_*k*_ is its size and *f*_*k*_ the fraction of it in *c*. It estimates the enrichment of the clumps that *c* touches, attenuated toward the clump mean when *c* splits clumps.

Both estimators are linear in 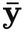 , so their exact variances follow from the weighted Theorem 3.10. In the application we use a block jackknife over 200 contiguous genomic blocks, which is valid for annotation-level quantities. Here 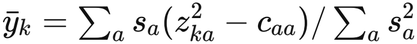 is the trait-pooled response (over traits with non-missing Z at SNP *k*), and **DA** are window-wise annotation-stratified LD scores computed from the same reference as Stage 3. We fit each annotation jointly with the full baseline model and marginally (base plus one annotation). Under the null of flat **w** within clumps, both are unbiased and the tests are valid. Gene-level enrichment is the mean clump-mean enrichment *Ŵ*_*t*_/*g*_*t*_ over HapMap3 SNPs within 10 kb of a protein-coding gene (gnomAD v2.1.1 coordinates; Karczewski et al. 2020). Its SE combines the tag model SEs, ignoring covariance between tags (which are pruned at *r*^2^ < 0.5). For the top loci we also report single-trait dominance: the share of the pooled response 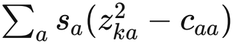 from the largest trait, and the effective number of contributing traits 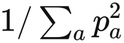.

#### Phenotype-category contrasts

Running LRCQ on a trait category *g* gives **ŵ** ^(*g*)^. For a locus or annotation, the contrast between categories *g*_1_ and *g*_2_ has an exact null variance, including cross-category genetic correlation and sample overlap, from the weighted Theorem 3.10 (Supplementary Note §S7.4). With a plug-in **Ĝ** estimated from the same data, that variance is biased upward and the test conservative (type-I error 0.013 at 5%). We therefore compute p-values by parametric bootstrap under *H*_0_, drawing **Z**^*\**^ from the pooled fit with **ŵ** clipped at 0 and recomputing the whole studentised statistic, with the debiased plug-in variance as its denominator (Supplementary Note §S7.4a). We report only these bootstrap p-values: normal-reference p-values from the debiased plug-in remained anti-conservative (type-I error 0.073 at 5%) and serve only as a diagnostic. Contrasts at a single dominant locus remain conservative when few genetically distinct trait clusters are analysed, and we report them as such. A locus enriched in every category is an omni-locus. A locus enriched in one category only is category-specific.

### Comparator per-SNP pleiotropy measures

On the same data we compute: (i) mean *χ*^2^ across traits, 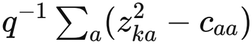 (ii) the LD-score-normalised mean *χ*^2^; (iii) the number of traits with *P* < 5 × 10^−8^, as in the GWAS Atlas pleiotropy counts (Watanabe et al. 2019); and (iv) the multi-trait omnibus Wald statistic 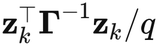, which tests whether a SNP is associated with any trait, accounting for trait correlation (cf. CPASSOC, Zhu et al. 2015). For annotation enrichment we compare with a pooled multi-trait s-LDSC-type regression on category LD scores using the true categories (an oracle in simulation).

### Simulations

We simulated GWAS summary statistics from the model above using real LD. Each replicate used three 3 Mb windows of UK Biobank in-sample LD (Weissbrod et al. 2020) restricted to HapMap3 SNPs (chr1:0-3 Mb, chr1:2-5 Mb, chr22:28-31 Mb; 3,178 SNPs), each treated as an independent block. Per-SNP signal used a genome-scale *M* = 1.1 × 10^6^, so that 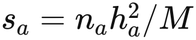 matched UK Biobank GWAS. Effects were drawn as **B** *∼ ℳ N* (**0, W**^1/2^**R**_*β*_**W**^1/2^, **H**/*M*), and Z-scores as 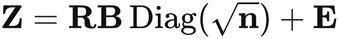 Cov(vec **E**) = **Γ** ⊗ **R** (implemented in the *lrcpq* R package).

In the baseline scenario: *q* = 300 traits; 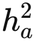 drawn from a scaled Beta(2,5) on [0.02, 0.6]; 2% of SNPs with non-zero *w* drawn from a log-normal (sdlog 1), rescaled to mean 1; independent traits without sample overlap; intercepts 1. Non-zero SNPs were placed preferentially in a synthetic four-level annotation: runs of 20 consecutive SNPs, with category frequencies 70/15/10/5% and relative probabilities of a non-zero effect of 1, 3, 6 and 12.

- Variations, one factor at a time:
- number of traits *q ∈* {30, 100, 1000};
- polygenicity: 0.5% or 10% non-zero, or all SNPs non-zero with sdlog 0.5;
- UK Biobank-like trait dependence: five trait clusters with *r*_*g*_ = 0.5, full sample overlap, phenotypic correlation 0.2;
- stratification: intercepts 1.1, either correctly supplied or misspecified as 1;
- genetic-effect correlation: *ρ* = 0.5 within clusters of five non-null SNPs in the same window (“local”), or between pairs in different windows (“cross”);
- heterogeneous enrichment: half of the traits with an independent **w**;
- noisy reference LD: estimated from 500 individuals, mimicking 1000 Genomes; tag threshold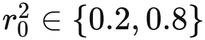.

Each scenario had 50 replicates. Truth for tag-level estimates is the tag-set estimand (eq. 3.6). Metrics: Pearson and Spearman correlation with the truth, regression slope (calibration), bias, RMSE, AUC for the top 10% of tags by true enrichment, calibration of standard errors, type-I error at null tags, and recovery of annotation enrichment.

#### Calibration of the tests

A second set of simulations used the same windows and generator to compare normal and skewness-corrected p-values (60 replicates each for the baseline, *q* = 30, infinitesimal and UK Biobank-like scenarios). Three further scenarios (30 replicates each) used the Pan-UKB trait structure: the 444 traits’ LDSC intercept matrix (sample overlap and phenotypic correlation), their genetic-covariance matrix rescaled to Pan-UKB *h*^2^, and their sample sizes, with enrichment as in the baseline. In two of them the estimator was given imperfect Stage 1-2 inputs: each trait’s *h*^2^ multiplied by an independent log-normal error with SD 0.2, or each intercept perturbed by a normal error with that trait’s Pan-UKB intercept SE. Null tags were those with tag-set estimand |*W*_*t*_| < 0.01. A third set (30 replicates each, Pan-UKB trait structure) compared plug-ins of **w** in the model SE and third cumulant, with 2%, 20% or 100% of SNPs non-null, and with **w** ≡ **1**; calibration was assessed at the true tag estimand (Supplementary Table S6). A fourth set used the Pan-UKB trait structure with 1000 Genomes LD and uniform enrichment (**w** ≡ **1**), so that every tag is null for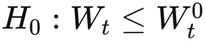. It compared normal, shifted-gamma, kurtosis-matched and Pearson type VI p-values for the score-type test (224 replicates; Supplementary Table S7).

#### Simulations of per-SNP estimation

We also simulated the same model with *w* drawn from scaled F distributions (with or without a point mass at zero) or from a matrix-normal generator, in blocks of 20-500 SNPs and 100-1,000 traits. The main findings were these. Accuracy rose with the number of traits (mean *R*^2^ 0.39, 0.53, 0.77 and 0.85 for *q* = 100, 200, 500 and 1,000). It depended little on GWAS sample size once the per-SNP signal was large. With a matrix-normal generator, per-SNP WLS was nearly unbiased for non-null SNPs (slope 0.95, adjusted *R*^2^ 0.96 at *N* = 10^5^). Per-SNP estimates degraded as the genome-wide SNP count *M* grew, because the per-SNP signal 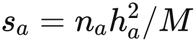 shrinks. At null SNPs, the SD of *ŵ* was 0.74, 0.86, 1.71, 5.13 and 19.3 for *M* = 10^4^ to 10^8^, and *R*^2^ with the truth fell from 0.97 to 0.07. This is the behaviour predicted by 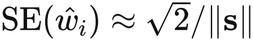 (Supplementary Note §S3.6). It motivated the use of genome-scale *M* in our simulations, the clump-level estimand and the emphasis on aggregation.

### Real data

#### GWAS summary statistics

Pan-UK Biobank (Karczewski et al. 2025) European-ancestry GWAS for all both-sexes phenotypes passing Pan-UKB’s EUR quality control (452 traits: 197 continuous, 145 categorical, 38 phecodes, 27 prescriptions, 24 ICD-10 codes and 21 biomarkers; 150 in Pan-UKB’s maximally independent set). Z-scores were 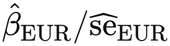 for the 1,094,844 HapMap3 SNPs of the Pan-UKB EUR LD-score release. A Z-score was set to missing if flagged low-confidence in EUR or if the EUR allele frequency (the control frequency for binary traits) was outside [0.01, 0.99]. *n*_*a*_ was the EUR sample size (cases plus controls), the same *N* that Pan-UKB used for its observed-scale LDSC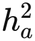. The model needs only the product 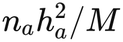, the expected 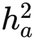 signal per unit LD score. Hence *n*_*a*_ and *h*^2^ must come from the same fit, and *M* must match the LD that enters **D**. With HapMap3-only windows, *M* is the number of HapMap3 SNPs (1,094,844), not Pan-UKB’s *M*_5−50_, which goes with full-panel LD scores. The genome-wide ratio of observed to expected mean *χ*^2^ signal under **w** ≡ **1** was 1.13 with this *M* and 7.0 with *M*_5−50_. The choice of *M* only rescales ***ŵ*** and its SEs by a common factor, which renormalisation to mean 1 removes exactly. Our total-*N* re-estimates matched Pan-UKB’s *h*^2^ equally well for binary traits at every prevalence (median ratio 0.96) and for continuous traits (0.98), which confirms the convention. Pairing effective sample sizes with these *h*^2^ values would understate *s*_*a*_ for binary traits by the factor *N*_eff_ /*N* (median 0.34). Eight prescription-medication traits whose files repeatedly failed to download were dropped, leaving 444. Missing Z-scores (median 0.05% of SNPs per trait) were imputed in Stage 3 at their expectation under. *w* = 1, *z*^2^ = *c*_*aa*_ + *s*_*a*_ℓ_*k*_, which contributes no information about enrichment.

#### Stages 1-2 in Pan-UKB

Per-trait 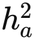 and intercepts *c*_*aa*_ were Pan-UKB’s own univariate LDSC estimates (EUR, HapMap3 LD scores from UK Biobank, *M*_5−50_ = 6,805,960). Cross-trait intercepts *c*_*ab*_ and genetic covariances were estimated by a vectorised cross-trait LDSC on all trait pairs with common weights; genetic covariances were rescaled to the Pan-UKB diagonal and projected to the positive semi-definite cone. Our re-estimated 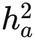 and intercepts correlated 0.992 and 0.985 with Pan-UKB’s across the 444 traits.

#### Motivating analysis

For a trait set 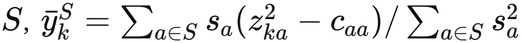 has expectation ℓ_*k*_ under **w** ≡ **1**. We split traits at random into halves *S*_1_, *S*_2_, regressed each 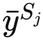 on LD score, and correlated the residuals across SNPs. Under **w** ≡ **1** and Gaussian Z, 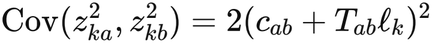 with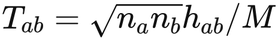 , which gives the null correlation exactly. The correlation of 1-Mb block means is reported descriptively.

#### LD reference

The primary reference is the 503 European-ancestry individuals of 1000 Genomes phase 3 (1000 Genomes Project Consortium 2015). We computed LD within 3-Mb windows spaced 2 Mb apart (1,383 windows, the layout of the UK Biobank LD release), with SNPs matched to Pan-UKB by position and alleles and signs aligned to the Pan-UKB alternative allele; 99.3% of the universe SNPs matched. *r*^2^ was bias-corrected with *N*_ref_ = 503. Estimates are kept for SNPs in the central 2 Mb of each window, and the MHC region (chr6:25-34 Mb) is excluded. As a sensitivity analysis we reran Stage 3 with UK Biobank in-sample LD (Weissbrod et al. 2020; 337k British-ancestry individuals), subset and allele-aligned to the same SNPs, over 20 windows (16,077 core SNPs; chr1:0-39 Mb and chr22:28-31 Mb). Over the window chr22:28-31 Mb, 1000 Genomes and UK Biobank *r* values correlated at 0.95. In-sample LD was not used genome-wide because of its computational cost (1-2 GB per window).

#### Trait subsets

All 444 traits (primary); Pan-UKB’s 150-trait maximally independent set; all 444 traits with equal weights; and, for the category contrast, the 51 blood and urine biomarkers (Pan-UKB category “Biological samples”) against the 252 assessment-centre traits (anthropometry, body composition, blood pressure, lifestyle and cognitive measures), using the Pan-UKB category field.

#### Functional follow-up

Enrichment of ***ŵ*** in the baseline-LF v2.2 annotations (Gazal et al. 2017, 2018; UK Biobank version distributed with Weissbrod et al. 2020), with each annotation’s low-frequency and common-variant versions merged (they are disjoint by MAF) via *Ê* _*c*_ and 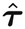 Gene-level enrichment over gene windows (±10 kb). Comparison of omni-loci with GWAS Atlas pleiotropy counts.

#### Replication cohorts

We matched Pan-UKB traits by content to GWAS from two cohorts without UK Biobank participants. FinnGen release 12 (Kurki et al. 2023) contributed 57 traits: 35 clinical laboratory measurements (sample-wise median values from the national laboratory database), height, body-mass index and weight, and 19 disease endpoints (median *n* 376,471; range 26,975-500,348). BioBank Japan (Sakaue et al. 2021) contributed 39 traits from its BBJ-only analyses, 33 quantitative traits and 6 diseases (median *n* 145,502; range 71,221-178,726; GRCh37 files from the GWAS Catalog). Variants were matched to the HapMap3 SNPs by position and both alleles (FinnGen endpoints by rsID), kept if their allele frequency was in [0.01, 0.99], and oriented to the Pan-UKB alternative allele; *n*_*a*_ was the total sample size. Stages 1-2 were re-estimated from the matched traits by univariate and cross-trait LDSC with Pan-UKB’s EUR (FinnGen) or EAS (BioBank Japan) LD scores and their *M*_5−50_. Stage 3 used the same windows, *M* and *r*^2^ < 0.5 tag selection as the discovery analysis, with LD from 1000 Genomes EUR (503 individuals) or EAS (504 individuals), and missing Z-scores imputed at their expectation under *w* = 1. A discovery tag was matched to the tag of the replication clump that contains its SNP. Its score-type p-values used the kurtosis-matched, shifted-gamma and normal references as in the main analysis. For a tag whose projected uniform null 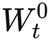 was negative in the replication data (40 of the 860 FinnGen and 23 of the 728 BioBank Japan tags tested above the uniform expectation), we used the test of *W*_*t*_ > 0 instead, which is conservative because 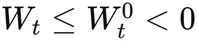 implies *W*_*t*_ < 0. The other tags of each run used normal-reference p-values only, so genome-wide discovery counts are not reported for the replication runs.

For reference, the same procedure was applied to Pan-UKB restricted to the matched traits; this reuses the discovery participants and is not a replication.

### Software

The *lrcpq* R package (version 0.3.1) implements Stages 1-4 of the method and the simulations. The analyses reported here used its default settings: OLS estimation, background-filled standard errors, and score-type tag tests with kurtosis-matched p-values. The package reads raw GWAS summary statistics (`munge_sumstats()`) and can be run from R or from a command-line tool modelled on `ldsc.py`. Two tutorials accompany it: a getting-started example on chromosome 22 bundled with the package, and a real-data walkthrough of the 444 Pan-UKB traits with a runnable 20-trait extract around TP53. LD can be computed from any PLINK reference panel; the 1000 Genomes phase 3 European reference used here (503 individuals, GRCh37, restricted to the 1,094,844 HapMap3 SNPs) is provided with the package release (see Data and code availability).

## Supporting information

Supplementary Information: Supplementary Figures S1-S4 and Supplementary Note (theory of LRCQ and LRCP)

## Data and code availability

Pan-UK Biobank GWAS summary statistics, LDSC estimates and the maximally independent trait set are publicly available from the Pan-UK Biobank project (Karczewski et al. 2025). FinnGen release 12 summary statistics are available from the FinnGen project (https://www.finngen.fi/en/access_results), and BioBank Japan summary statistics from the GWAS Catalog (the study accessions are listed in Supplementary Table S9). The 1000 Genomes phase 3 genotypes, the UK Biobank in-sample LD matrices and the baseline-LF v2.2 annotations (Weissbrod et al. 2020) and gnomAD v2.1.1 constraint metrics are public. The R package *lrcpq* (version 0.3.1) is available at https://github.com/unfated/lrcpq; its releases include the 1000 Genomes EUR HapMap3 LD reference. All analysis code will be made publicly available upon journal publication and is available from the corresponding author on request.

## Acknowledgements

We thank Justin D. Tubbs for contributions to an earlier version of this work.

## Funding

This work was supported by the State Key Laboratory of Brain and Cognitive Sciences, The University of Hong Kong.

## Competing interests

L.G.C. is the founder and chief executive officer of Gene to Health Limited and of 2HEALTH AI LIMITED. The business activities of these companies are unrelated to the work reported in this article. The authors declare no other competing interests.

## Author contributions

L.G.C. conceived the study, developed the method and its theory, implemented the software, performed the simulations and the real-data analyses, and wrote the manuscript. T.Q.T. contributed to the revision and audit of the manuscript and to supplementary real-data analyses. P.C.S. contributed to the theoretical conception of the method, reviewed the data analyses and guided the revision of the manuscript.

## Figures and tables

**Table 3. Unknown parameters by stage** (in Methods, “Estimation”): estimator, identification, inference and Pan-UKB status for stages 1-4.

**Supplementary Figures S1-S4**, with legends, are in the Supplementary Information.

**Supplementary Table S1**. Per-SNP solves (exact; ridge with *λ* = 10^−4^, 10^−3^ and 10^−2^) and the tag-set solve in one 3-Mb window of UK Biobank in-sample LD: SD of the estimates and correlation with the truth. **Supplementary Table S2**. The 444 Pan-UKB traits, with category, sample size and LDSC *h*^2^ and intercept. **Supplementary Table S3**. Trait-cluster jackknife versus model SEs in simulation. **Supplementary Table S4**. Calibration of normal and skewness-corrected tests at null tags, including the Pan-UKB-structure scenarios with Stage 1-2 plug-in errors. **Supplementary Table S5**. Rectification and Method D applied to the OLS estimates. **Supplementary Table S6**. Plug-ins of **w** in the model SE and third cumulant, under the Pan-UKB trait structure with 2%, 20% and 100% non-null SNPs and **w** ≡ **1**: SD of standardised errors at the true tag estimand, upper-tail rates and median SE ratio to method A. **Supplementary Table S7**. Calibration of the score-type tag test under uniform enrichment with the Pan-UKB trait structure: normal, shifted-gamma, kurtosis-matched and Pearson type VI p-values. **Supplementary Table S8**. Null configuration of the score-type test in Pan-UKB: tags and loci at FDR 5% under the primary nonnegative null with the mean stated in **D**, the configuration without the nonnegative fix, and the mean stated with uncorrected **R** ∘ **R** (minimum distance, or window total also fixed), with the number of LRCP FDR-5% pairs that involve tags leaving the *W*_*t*_ > 0 set. **Supplementary Table S9**. Replication in FinnGen R12 and BioBank Japan: matched traits with study identifiers and sample sizes, summary by cohort (with the Pan-UKB matched-trait reference), and for each of the 1,799 tags with *W*_*t*_ > 0 the discovery estimate and q-value and the replication estimate, SE, score-type p-values and within-set q-values.

## Notes

https://github.com/unfated/lrcpq

https://pan.ukbb.broadinstitute.org

### Summary of Updates

Version 3. (1) Score-type test: for the roughly 5% of tags whose null value under uniform enrichment is negative after LD projection, the null configuration is now the nonnegative per-SNP enrichment closest to the background-filled plug-in with the required projected mean (new Supplementary Table S8). At a tag-level FDR of 5%, 881 LD clumps in 638 distinct loci are enriched above the uniform expectation; the abstract and genome-wide results are updated. (2) Replication in two cohorts without UK Biobank participants, FinnGen release 12 and BioBank Japan (new Supplementary Table S9 and Supplementary Figure S4). (3) Supplementary figures and the shared theory note are now provided together as one Supplementary Information PDF. (4) The software description refers to lrcpq v0.3.1.

