## Supplementary Information: Supplementary Figures S1-S4 and Supplementary Note (theory of LRCQ and LRCP) for "LRCQ: phenome-wide heritability enrichment of genomic loci from GWAS summary statistics"

**PDF version: 2026-11-20**

**Contents**

1. Supplementary Figures S1-S4
2. Supplementary Note: Theory of LRCQ and LRCP

Supplementary Tables S1-S9 are provided as a separate Excel file (LRCQ-Supplementary-Tables).

### Supplementary Figures

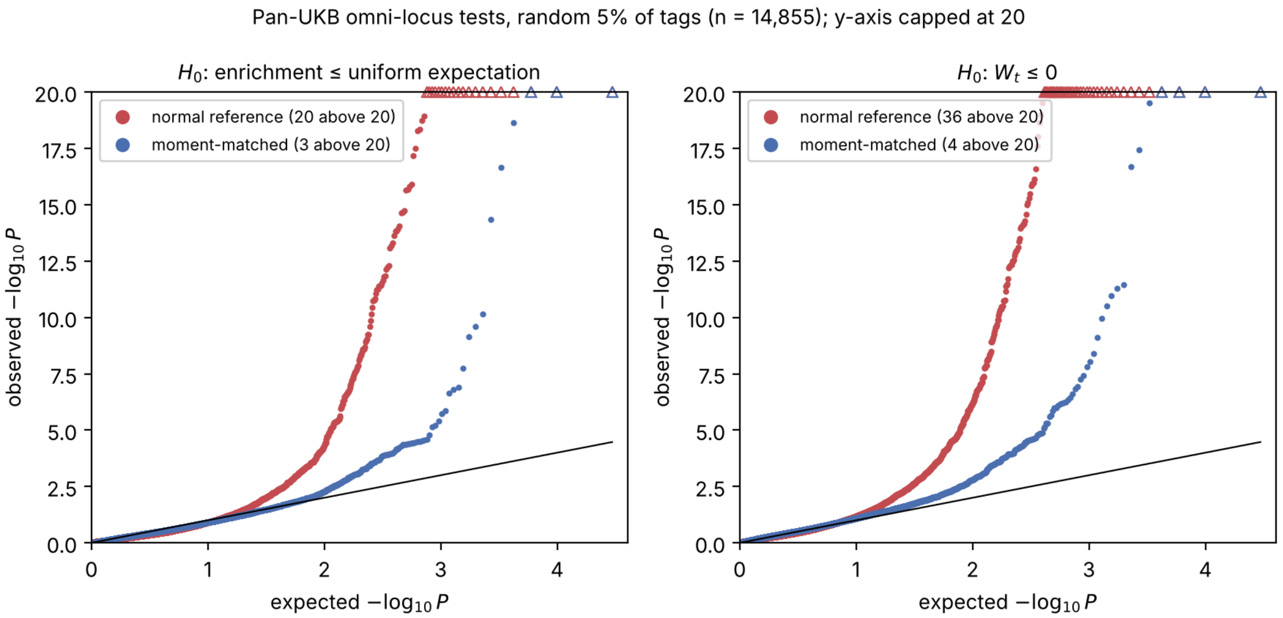


**Supplementary Figure S1.** QQ plots of the Pan-UKB omni-locus tests ($H_0 : W_t \leq W_t^0$, the expectation under uniform enrichment, and $H_0 : W_t \leq 0$) for a random 5% of tags (14,855), for the score-type tests with moment-matched and normal-reference p-values. The black line is $y = x$. The y-axis is capped at 20; open triangles on the top edge mark tags with larger $-\log_{10} P$, and the legend gives their number. Because these are real data with many associated loci, the upper tail is expected to rise above the line; the lower part of each curve shows the calibration.

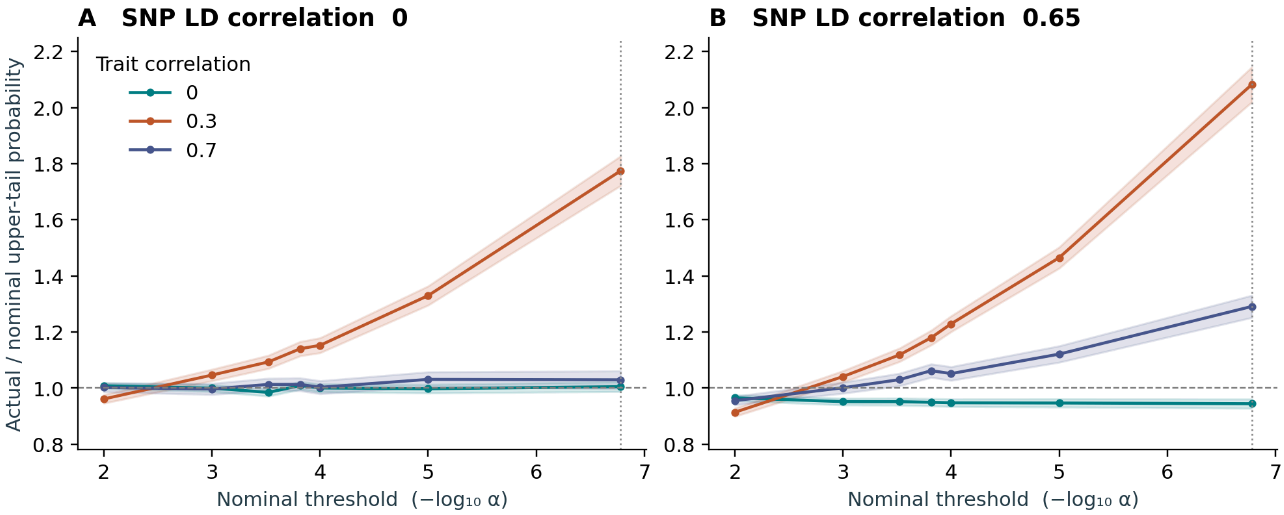


**Supplementary Figure S2.** Upper-tail calibration of the kurtosis-matched reference when the covariance is known. Two SNPs with uniform enrichment and 24 equally weighted traits (scale 0.3 each); the null statistic is then exactly a weighted sum of independent $\chi^2$ variables. Each point is the actual-to-nominal ratio of the upper-tail probability at a kurtosis-matched threshold, estimated from 120,000 exponentially tilted draws; ribbons are $\pm 1.96$ Monte Carlo standard errors. (a) SNP LD correlation 0; (b) 0.65. Colours give the equicorrelation between traits, used for both the genetic and the intercept covariance. The dotted vertical line marks $0.05/301{,}693$. With nuisance parameters fixed, this isolates the error of the tail approximation.

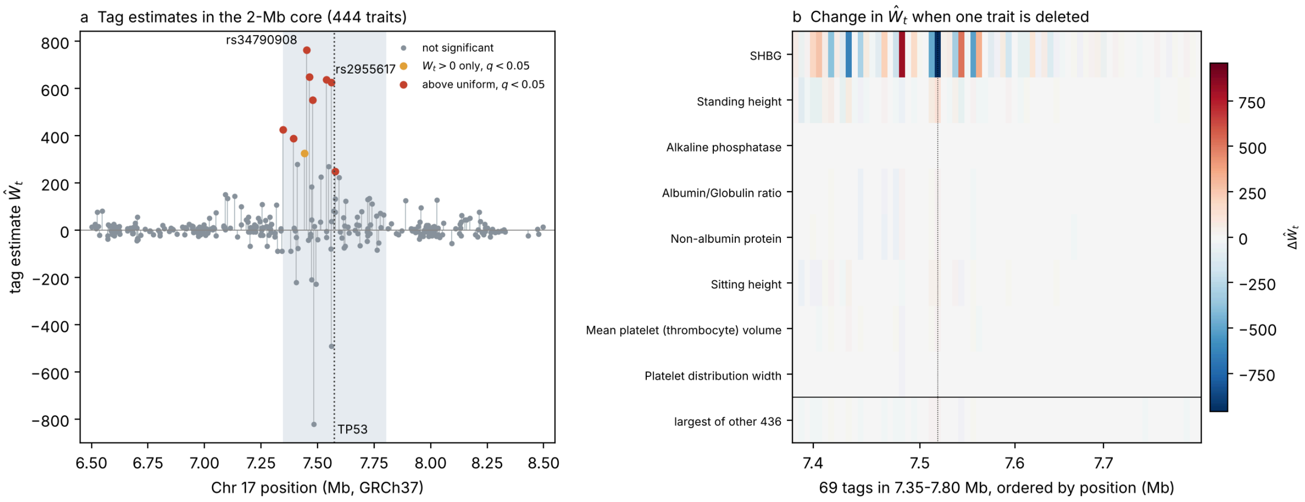


**Supplementary Figure S3.** Tag estimates and single-trait influence in the TP53 region (chr17, 3-Mb window 6-9 Mb, 2-Mb core 6.5-8.5 Mb; 444 Pan-UKB traits; GRCh37). (a) Estimates $\hat{W}_t$ of the 298 core tags, coloured by the score-type test: enriched above the uniform expectation at FDR 5% (red), $W_t > 0$ only (orange), neither (grey). The dotted line marks TP53 (rs78378222). (b) Change in $\hat{W}_t$ when one trait is deleted (LD, tags and the renormalisation held fixed), for the tags between 7.35 and 7.80 Mb (shaded in a). Rows are the eight traits with the largest absolute change at any core tag; the last row shows, for each tag, the largest change among the other 436 deletions. Deleting SHBG moves rs2955617 (dotted line) from 639 to $-319$, and deleting standing height moves it to 753; the Spearman correlation between the full and the deleted tag rankings is at least 0.998 for 440 of the 444 deletions (minimum 0.74, SHBG). Deletion shows how much an estimate depends on the observed trait panel; it is not a confidence interval or a jackknife.

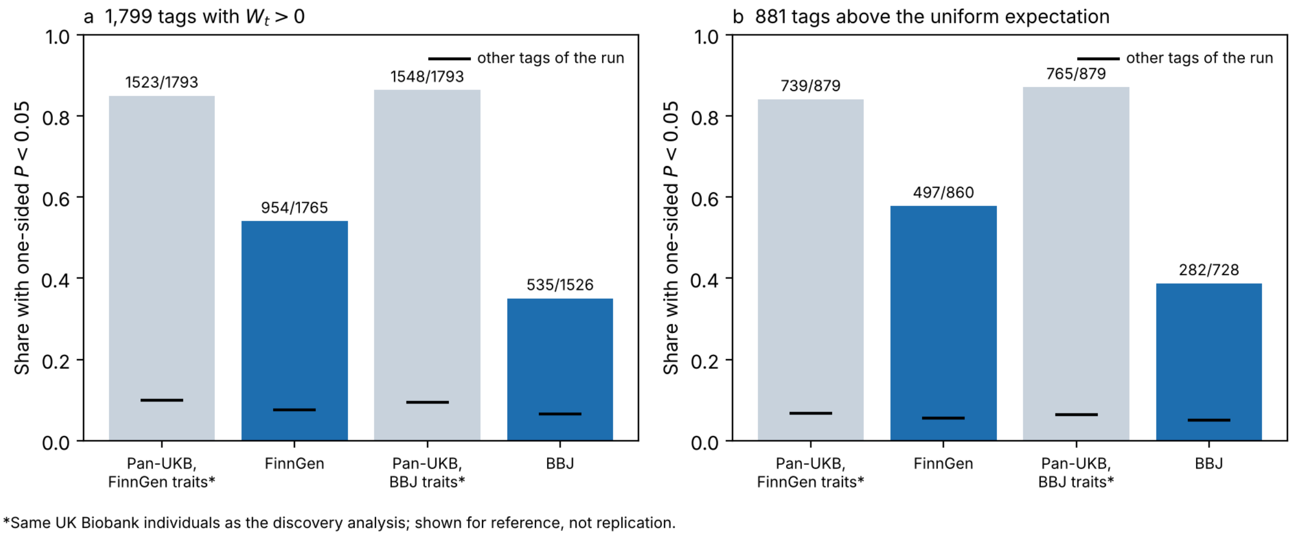


**Supplementary Figure S4.** Replication of the Pan-UKB enriched tags in FinnGen R12 (57 matched traits) and BioBank Japan (39 matched traits), each analysed with its own Stage 1-2 estimates and an ancestry-matched 1000 Genomes LD reference. Bars: share of testable discovery tags with one-sided score-type $P < 0.05$ for (a) $W_t > 0$, among the 1,799 tags with $W_t > 0$ in Pan-UKB, and (b) $W_t$ above the uniform expectation, among the 881 tags enriched above it in Pan-UKB; numbers give tags with $P < 0.05$ over testable tags. Black marks: share of the other tags of the same run with $P < 0.05$ (normal reference). Light bars: Pan-UKB restricted to the matched traits, which reuses the discovery participants and is shown only as a ceiling.

### Supplementary Note: Theory of LRCQ and LRCP

*A unified framework for phenome-wide genetic-effect variance (enrichment) and genetic-effect correlation from GWAS summary statistics.*

Lane Guolan Chen, Thuan Quoc Thach, Pak C Sham. Supplementary Note shared by the LRCQ and LRCP papers. Version of 2026-11-19.

Every numbered identity here was checked numerically, and Appendix A lists the results. All estimators and tests are implemented in the R package lrcpq (version 0.3.1, https://github.com/unfated/lrcpq), whose releases also distribute the 1000 Genomes EUR HapMap3 LD reference. The numerical-check scripts will be made publicly available upon journal publication and are available from the corresponding author (Lane Guolan Chen,) on request.

---

#### S0. Notation and overview

| Symbol | Meaning |
|---|---|
| $m, q$ | number of SNPs (in the reference panel / window) and of traits (GWAS) |
| $n_a, n_{ab}$ | GWAS sample size of trait $a$; number of individuals shared by GWAS $a$ and $b$ |
| $\mathbf{X}$ $(n \times m)$, $\mathbf{Y}$ $(n \times q)$ | standardised genotypes and phenotypes |
| $\mathbf{B}$ $(m \times q)$ | joint (causal, "direct") genetic effects; column $\boldsymbol{\beta}_a$ = effects on trait $a$; row $\boldsymbol{\beta}_{s\cdot}$ = effects of SNP $s$ across the phenome |
| $\mathbf{R}$, $r_{kl}$ | LD (genotype correlation) matrix of the population |
| $\mathbf{D} = \mathbf{R} \circ \mathbf{R}$ | element-wise squared LD, $D_{kl} = r_{kl}^2$; $\ell_k = \sum_s r_{ks}^2$ is the LD score |
| $\tilde{\boldsymbol{\Sigma}}$ | normalised SNP-wise effect covariance, $\mathrm{tr}\,\tilde{\boldsymbol{\Sigma}} = m$ |
| $\mathbf{w} = \mathrm{diag}(\tilde{\boldsymbol{\Sigma}})$ | **genetic-effect enrichment** $w_\beta$ (mean 1 over the $m$ SNPs) |
| $\mathbf{R}_\beta = \mathbf{W}^{-1/2}\tilde{\boldsymbol{\Sigma}}\mathbf{W}^{-1/2}$, $\rho_{ij}$ | **genetic-effect correlation**; $\mathbf{W} = \mathrm{Diag}(\mathbf{w})$ |
| $\mathbf{C} = \tilde{\boldsymbol{\Sigma}} - \mathbf{W}$ | off-diagonal part of $\tilde{\boldsymbol{\Sigma}}$, $C_{ij} = \rho_{ij}\sqrt{w_i w_j}$ |
| $h_a^2$, $h_{ab}$ | SNP heritability and co-heritability (genetic covariance), $h_{ab} = r_{g,ab} h_a h_b$ |
| $s_a = n_a h_a^2/m$, $t_{ab} = \sqrt{n_a n_b}\, h_{ab}/m$ | per-trait and per-pair signal scales ($t_{aa} = s_a$) |
| $c_{ab}$ | intercepts: $c_{aa} = 1+$(stratification/relatedness inflation), $c_{ab} = n_{ab}\rho_{ab}^P/\sqrt{n_a n_b}$ |
| $\mathbf{Z}$ $(m \times q)$ | GWAS Z-scores, $z_{ka}$ = SNP $k$, trait $a$ |
| $\mathbf{S} = \mathrm{Diag}(\mathbf{s})$, $\mathbf{T} = (t_{ab})$ | |

Equivalently, with the unnormalised SNP-wise covariance $\boldsymbol{\Sigma}_\beta = E[\mathbf{B}\mathbf{B}^\top]/q = (\bar{h}^2/m)\,\tilde{\boldsymbol{\Sigma}}$ and $\bar{h}^2 = q^{-1}\sum_a h_a^2$, the enrichment is $\mathbf{w} = \mathrm{diag}(\boldsymbol{\Sigma}_\beta)/\operatorname{avg}\mathrm{diag}(\boldsymbol{\Sigma}_\beta)$, and $\mathbf{R}_\beta$ is the correlation matrix of $\boldsymbol{\Sigma}_\beta$.

**What the two methods estimate.**

- **LRCQ** (LD score regression on $\chi^2$ across many GWAS): $\mathbf{w}$. It uses the $mq$ squared Z-scores $z_{ka}^2$ and LD components $\mathbf{D}$.
- **LRCP** (likelihood and regression on cross-products of Z scores): $\rho_{ij}$ (equivalently $C_{ij}$). It uses Z-score cross-products $z_{ka}z_{lb}$ and LD cross components $r_{ki}r_{lj}$.

Both rest on one moment identity (Theorem 1). Their estimands, and the conditions for identifying them, follow from it (§S1.6, §S4.1).

**Analysis stages.** Both methods run in four stages. Stage 1 estimates SNP heritabilities $h_a^2$ and genetic covariances $h_{ab}$, and Stage 2 the intercepts $c_{aa}$ and $c_{ab}$, both by genome-wide (cross-trait) LD score regression. Stage 3 is LRCQ, which estimates $\mathbf{w}$ with the Stage 1–2 quantities held fixed. Stage 4 is LRCP, which estimates $\rho_{ij}$ (or $C_{ij}$) for candidate pairs selected using Stage 3.

---

#### S1. Model, assumptions and the master moment identity

##### S1.1 From joint effects to GWAS marginal estimates

For each trait $a$ the individual-level model is $\mathbf{y}_a = \mathbf{X}\boldsymbol{\beta}_a + \mathbf{e}_a$, with columns of $\mathbf{X}$ and $\mathbf{y}_a$ standardised in the GWAS sample of size $n_a$ and $\mathrm{Var}(e_{ia}) = 1 - h_a^2$. The GWAS marginal estimate of SNP $k$ is the per-SNP OLS slope $\hat{\alpha}_{ka} = \mathbf{x}_k^\top \mathbf{y}_a / n_a$. Stacking over SNPs:

$$\hat{\boldsymbol{\alpha}}_a = \frac{\mathbf{X}^\top\mathbf{X}}{n_a}\boldsymbol{\beta}_a + \frac{\mathbf{X}^\top\mathbf{e}_a}{n_a} = \mathbf{R}\boldsymbol{\beta}_a + \mathbf{u}_a, \qquad \mathbf{u}_a = \Big(\frac{\mathbf{X}^\top\mathbf{X}}{n_a} - \mathbf{R}\Big)\boldsymbol{\beta}_a + \frac{\mathbf{X}^\top\mathbf{e}_a}{n_a}.$$

**Lemma 1.1.** Assume (i) genotypes of different individuals are independent with correlation $\mathbf{R}$, (ii) $\mathbf{e}_a$ is independent of $\mathbf{X}$ and $\mathbf{B}$, and (iii) the total variance explained by the SNPs in the model is small, $\boldsymbol{\beta}_a^\top \mathbf{R}\boldsymbol{\beta}_a = h_a^2 \ll 1$, with genotypes standardised in the population and with bounded fourth moments. Then $E[\mathbf{u}_a \mid \mathbf{B}] = \mathbf{0}$ and $\mathrm{Var}(\mathbf{u}_a \mid \mathbf{B}) = \mathbf{R}/n_a\,(1 + O(h_a^2))$.

*Proof.* $E[\mathbf{X}^\top\mathbf{X}/n_a] = \mathbf{R}$ and $E[\mathbf{X}^\top\mathbf{e}_a] = \mathbf{0}$ give the mean. For the variance, $\mathrm{Var}(\mathbf{X}^\top\mathbf{e}_a/n_a) = (1 - h_a^2)\mathbf{R}/n_a$. For Gaussian genotypes the first term of $\mathbf{u}_a$ has covariance $\{(\boldsymbol{\beta}_a^\top\mathbf{R}\boldsymbol{\beta}_a)\mathbf{R} + (\mathbf{R}\boldsymbol{\beta}_a)(\mathbf{R}\boldsymbol{\beta}_a)^\top\}/n_a$. For other genotype distributions a fourth-cumulant term is added, which is of the same order under bounded fourth moments. The two terms are uncorrelated. □

The GWAS standard error is $\mathrm{se}(\hat{\alpha}_{ka}) = \sqrt{(1 - \hat{\alpha}_{ka}^2)/(n_a - 2)} \approx n_a^{-1/2}$. Hence

$$\mathbf{z}_a = \sqrt{n_a}\,\hat{\boldsymbol{\alpha}}_a = \sqrt{n_a}\,\mathbf{R}\boldsymbol{\beta}_a + \sqrt{n_a}\,\mathbf{u}_a, \qquad \text{i.e.} \qquad \mathbf{Z} = \mathbf{R}\mathbf{B}\ \mathrm{Diag}(\sqrt{\mathbf{n}}) + \mathbf{U}^*, \quad (1.1)$$

where $\mathbf{U}^*$ has columns $\sqrt{n_a}\mathbf{u}_a$. The approximation errors are $O(h^2)$, as in LDSC (Bulik-Sullivan et al. 2015a, Supplementary Note). Appendix A, check M1, shows a relative error of 3.5% at a deliberately extreme per-SNP $h^2$ of 0.7%. The bound is in terms of the trait's total $h_a^2$, not the largest single-SNP effect; the term $(\mathbf{R}\boldsymbol{\beta}_a)(\mathbf{R}\boldsymbol{\beta}_a)^\top$ can be locally larger near strong associations. We therefore treat $\mathrm{Var}(\mathbf{U}^*)$ in (1.1) as the working noise model of summary-statistic methods rather than an exact result.

#### S1.2 Covariance of sampling errors across GWAS

**Lemma 1.2.** Let GWAS $a$ and $b$ share $n_{ab}$ individuals, and let $\rho^P_{ab}$ be the phenotypic correlation of traits $a, b$ among them. Then, to the order of Lemma 1.1,

$$E[\sqrt{n_a}\mathbf{u}_a\,\sqrt{n_b}\mathbf{u}_b^\top] = c_{ab}\,\mathbf{R}, \qquad c_{ab} = \frac{n_{ab}\,\rho^P_{ab}}{\sqrt{n_a n_b}}, \quad c_{aa} = 1.$$

*Proof.* $\sqrt{n_a n_b}\,E[\mathbf{u}_a\mathbf{u}_b^\top] = \frac{1}{\sqrt{n_a n_b}}\sum_{i\in a}\sum_{i'\in b} E[\mathbf{x}_i\mathbf{x}_{i'}^\top \epsilon_{ia}\epsilon_{i'b}]$, where $\epsilon$ is the part of $y$ not captured by the marginal mean. Terms with $i \neq i'$ vanish by independence. The $n_{ab}$ shared individuals contribute $\mathbf{R}\cdot \mathrm{Cov}(y_a, y_b) = \mathbf{R}\rho^P_{ab}$ to leading order. □

This is the cross-trait LDSC intercept (Bulik-Sullivan et al. 2015b). Population stratification and cryptic relatedness add a constant to $c_{aa}$ (Bulik-Sullivan et al. 2015a). For UK Biobank GWAS on the same individuals, $n_{ab} = n_a = n_b$ and $c_{ab} = \rho^P_{ab}$. Note that $\rho^P_{ab}$ is the total phenotypic correlation in the overlapping sample, not a residual correlation; check M1 uses the total correlation and matches.

#### S1.3 Assumptions on the random effects

Throughout, $\mathbf{B}$ is random, $\mathbf{U}^*$ has mean zero given $\mathbf{B}$ (Lemma 1.1; its conditional variance depends on $\mathbf{B}$ only at relative order $h_a^2$), and:

- **(A1) Mean zero.** $E[\mathbf{B}] = \mathbf{0}$.
- **(A2) Separable second moments.** For all traits $a, b$: $E[\boldsymbol{\beta}_a\boldsymbol{\beta}_b^\top] = \frac{h_{ab}}{m}\,\tilde{\mathbf{\Sigma}}$, with $\mathrm{tr}\,\tilde{\mathbf{\Sigma}} = m$.

(A2) says the SNP-wise covariance pattern is shared by all traits up to a trait-pair scale. That scale is the genetic covariance, because $\sum_k E[\beta_{ka}\beta_{kb}] = h_{ab}$. For a single trait, (A2) says the SNP-wise covariance of trait $a$ is $(h_a^2/\bar{h}^2)\mathbf{\Sigma}_\beta$. It is equivalent to $\mathrm{Cov}(\mathrm{vec}\,\mathbf{B}) = \mathbf{V}\otimes\tilde{\mathbf{\Sigma}}$ with $V_{ab} = h_{ab}/m$, the second-moment structure of the matrix-normal model (§S2). **No distributional assumption beyond second moments is needed for the identity below.** Normality enters only through variances and likelihoods (§S3.4, §S4.5, §S5).

Here $h_a^2$ means $\sum_k E[\beta_{ka}^2]$, the $a$-th diagonal entry of $E[\mathbf{B}^\top\mathbf{B}]$. With LD and non-zero $\mathbf{C}$, the genetic variance of trait $a$ is $(h_a^2/m)\,\mathrm{tr}(\mathbf{R}\tilde{\mathbf{\Sigma}})$. This differs from $h_a^2$ by $(h_a^2/m)\sum_{i\neq j} r_{ij}C_{ij}$, which is negligible unless correlated SNPs are also in strong LD.

When (A2) fails, §S3.2 and §S2.3 give what the estimators target instead.

#### S1.4 The master identity

**Theorem 1.** Under (1.1), Lemma 1.2 and (A1)–(A2), for any SNPs $k, l$ and traits $a, b$:

$$\boxed{E[z_{ka}z_{lb}] = c_{ab}\,r_{kl} + t_{ab}\,(\mathbf{R}\tilde{\mathbf{\Sigma}}\mathbf{R})_{kl}} \tag{1.2}$$

with

$$(\mathbf{R}\tilde{\boldsymbol{\Sigma}}\mathbf{R})_{kl} = \sum_{s=1}^{m} w_s\, r_{ks} r_{ls} + \sum_{i<j} \rho_{ij} \sqrt{w_i w_j}\, \big(r_{ki} r_{lj} + r_{kj} r_{li}\big). \tag{1.3}$$

In matrix form, $E[\mathbf{z}_a \mathbf{z}_b^\top] = c_{ab}\mathbf{R} + t_{ab}\mathbf{R}\tilde{\boldsymbol{\Sigma}}\mathbf{R}$.

*Proof.* From (1.1), $E[\mathbf{z}_a \mathbf{z}_b^\top] = \sqrt{n_a n_b}\, \mathbf{R}\, E[\boldsymbol{\beta}_a \boldsymbol{\beta}_b^\top]\, \mathbf{R} + E[\sqrt{n_a n_b}\, \mathbf{u}_a \mathbf{u}_b^\top]$. The cross terms vanish because $E[\mathbf{U}^* \mid \mathbf{B}] = \mathbf{0}$. Substituting (A2) and Lemma 1.2 gives (1.2). For (1.3), write $\tilde{\boldsymbol{\Sigma}} = \mathbf{W} + \mathbf{C}$. Then $(\mathbf{RCR})_{kl} = \sum_{i \neq j} r_{ki} C_{ij} r_{jl}$. Grouping the ordered pairs $(i, j)$ and $(j, i)$ and using $C_{ij} = C_{ji}$ gives the unordered sum. □

**Corollary 1 (LRCQ equation; squares, $k = l, a = b$).**

$$E[z_{ka}^2] = c_{aa} + s_a \Big[ \sum_s w_s r_{ks}^2 + 2 \sum_{i<j} \rho_{ij} \sqrt{w_i w_j}\, r_{ki} r_{kj} \Big] = c_{aa} + s_a (\mathbf{D}\mathbf{w})_k + s_a (\mathbf{RCR})_{kk} \tag{1.4}$$

**Corollary 2 (LRCP equation; cross-products).** Equation (1.2) with (1.3). Two special cases matter:

- *Cross-block* ($k$ in block 1, $l$ in block 2, no LD between blocks): $r_{kl} = 0$ and $r_{kj} = r_{li} = 0$ for $i$ in block 1 and $j$ in block 2. So

$$E[z_{ka} z_{lb}] = t_{ab} \sum_{i \in 1, j \in 2} \rho_{ij} \sqrt{w_i w_j}\, r_{ki} r_{lj} = t_{ab} (\mathbf{R}_1 \mathbf{C}_{12} \mathbf{R}_2)_{kl}. \tag{1.5}$$

  The cross-block expectation does not depend on $\mathbf{w}$, on the intercepts, or on any within-block correlation.
- *LDSC as a special case*: with $\mathbf{w} = \mathbf{1}$ and $\mathbf{C} = \mathbf{0}$, (1.4) is $E z_{ka}^2 = c_{aa} + s_a \ell_k$ (Bulik-Sullivan et al. 2015a). (1.2) gives cross-trait LDSC, $E z_{ka} z_{kb} = c_{ab} + t_{ab} \ell_k$ (Bulik-Sullivan et al. 2015b). With $\mathbf{w}$ given by annotations, $w_s = \sum_c \tau_c a_{cs}$, (1.4) is stratified LDSC (Finucane et al. 2015). LRCQ is therefore s-LDSC with one "annotation" per SNP, pooled over $q$ traits.

**Remark (pair counting).** In (1.3) each unordered SNP pair $\{i, j\}$ enters once. In the squared-Z equation (1.4) the factor 2 is the sum of the two equal terms $r_{ki} r_{kj} + r_{kj} r_{ki}$. In the cross-product equation for $k \neq l$ the two terms $r_{ki} r_{lj}$ and $r_{kj} r_{li}$ differ and no factor 2 appears. The scale of the genetic term is $t_{ab} = \sqrt{n_a n_b}\, h_{ab}/m$.

##### S1.5 The cross-block regressor carries no factor 2

By (1.5), a regression of cross-block products $z_{ka} z_{la}$ on $g_{kl} = \frac{nh^2}{m} r_{ki} r_{lj}$ has slope $\rho_{ij} \sqrt{w_i w_j}$, which is $\rho_{ij}$ when $w_i = w_j = 1$. A regressor that counts the pair twice, $\frac{nh^2}{m}\, 2 r_{ki} r_{lj}$, returns exactly half of that.

We checked this with a simple design (check F1): $n = 10{,}000$, $h^2 = 0.8$, two LD blocks of 50 SNPs with no LD between them, $q = 200$ traits, all SNPs causal with $w = 1$, one non-zero cross-block $\rho$, 400 replicates.

| true $\rho$ | slope, regressor (1.5) (MC s.e.) | slope, double-counted regressor |
|---|---|---|
| 0.8 | 0.817 (0.041) | 0.408 |
| 0.5 | 0.485 (0.039) | 0.243 |
| 0.2 | 0.205 (0.038) | 0.102 |

| true $\rho$ | slope, regressor (1.5) (MC s.e.) | slope, double-counted regressor |
|---|---|---|
| −0.5 | −0.518 (0.039) | −0.259 |

Check M1 confirms (1.2) from individual-level data: at the SNP with the largest signal, $Ez^2$ is 52.2 empirically and 52.8 from (1.4); the double-counted form gives 59.1.

##### S1.6 What squared Z-scores identify: the LRCQ estimand

By (1.4), the squared Z-scores of all traits depend on $\tilde{\boldsymbol{\Sigma}}$ only through the length-$m$ vector

$$\boldsymbol{\lambda} = \operatorname{diag}(\mathbf{R}\tilde{\boldsymbol{\Sigma}}\mathbf{R}) = \mathbf{D}\mathbf{w} + \operatorname{diag}(\mathbf{R}\mathbf{C}\mathbf{R}).$$

**Proposition 1.6 (identification).** From $\{z_{ka}^2\}$ alone, with $c_{aa}$ and $s_a$ known, $\boldsymbol{\lambda}$ is identified but $(\mathbf{w}, \mathbf{C})$ is not. If $\mathbf{D}$ is invertible, $\mathbf{w}$ is identified under the working assumption $\operatorname{diag}(\mathbf{R}\mathbf{C}\mathbf{R}) = \mathbf{0}$. This holds whenever no two genetically correlated SNPs share an LD partner, in particular when every non-zero $\rho_{ij}$ links SNPs in different LD blocks.

*Proof.* The squares' means are $c_{aa}\mathbf{1} + s_a\boldsymbol{\lambda}$. Any $(\mathbf{w}', \mathbf{C}')$ with the same $\boldsymbol{\lambda}$ yields identical distributions of the squares' first moments. The pair $(\mathbf{w} + \mathbf{D}^{-1}\operatorname{diag}(\mathbf{R}\mathbf{C}\mathbf{R}), \mathbf{0})$ is one such. □

**Theorem 2 (bias of chi-square-only LRCQ).** Let $\hat{\mathbf{w}}$ be any estimator that is linear and unbiased for $\mathbf{w}$ under $\mathbf{C} = \mathbf{0}$ (OLS, WLS with fixed weights, §S3). In general it satisfies

$$E[\hat{\mathbf{w}}] = \mathbf{w} + \mathbf{D}^{-1}\operatorname{diag}(\mathbf{R}\mathbf{C}\mathbf{R}). \tag{1.6}$$

*Proof.* Write $\hat{\mathbf{w}} = \mathcal{L}(\operatorname{vec}\mathbf{Y}^{(2)})$ with $\mathcal{L}$ linear. Unbiasedness under $\mathbf{C} = \mathbf{0}$ for every $\mathbf{w}$ means $\mathcal{L}(\mathbf{s} \otimes \mathbf{D}\mathbf{w}) = \mathbf{w}$ for all $\mathbf{w}$. In general $E\operatorname{vec}\mathbf{Y}^{(2)} = \mathbf{s} \otimes \boldsymbol{\lambda} = \mathbf{s} \otimes \mathbf{D}(\mathbf{D}^{-1}\boldsymbol{\lambda})$, so $E\hat{\mathbf{w}} = \mathbf{D}^{-1}\boldsymbol{\lambda} = \mathbf{w} + \mathbf{D}^{-1}\operatorname{diag}(\mathbf{R}\mathbf{C}\mathbf{R})$. □

Check L3 compares the observed bias with (1.6): correlation 0.999, slope 1.02.

**Consequences.**

- Chi-square-only LRCQ is the main estimator. Its estimand is $\mathbf{w}^\dagger = \mathbf{D}^{-1}\boldsymbol{\lambda}$, the "LD-resolved" phenome-wide variance. It equals $\mathbf{w}$ when local genetic-effect correlations are absent.
- When local $\rho_{ij} > 0$ between SNPs in LD, the bias is positive near the pair: correlated causal SNPs in LD look more enriched. In summary statistics, an observed association pattern is the sum of a functional link and LD.
- The local $\rho$ terms in (1.4) are not estimable from squares alone. They become estimable once cross-products are used (§S3.9, §S4).

##### S1.7 Plug-in quantities from Stages 1–2

- **Heritability.** If $\hat{s}_a = \kappa_a s_a$, OLS gives $E[\hat{\mathbf{w}}] = \mathbf{w}\sum_a \kappa_a s_a^2 / \sum_a \kappa_a^2 s_a^2$. A common misscaling ( $\kappa_a \equiv \kappa$) rescales $\hat{\mathbf{w}}$ by $1/\kappa$ and is removed by renormalising $\hat{\mathbf{w}}$ to mean 1 over the genome. Rankings and tests of $w_i = 0$ are unaffected.
- **LD-dependent architecture.** Stage-1 LDSC assumes $\mathbf{w} = \mathbf{1}$. If $\mathbf{w}$ correlates with LD score, LDSC's $\hat{h}^2$ is biased (Gazal et al. 2017; Speed et al. 2017). A practical remedy is one alternation: re-estimate $h_a^2$

by regressing $z_{ka}^2$ on the enrichment-weighted LD score $(\mathbf{D}\hat{\mathbf{w}})_k$, then re-run Stage 3.

- **Intercepts.** Intercepts must come from Stage 2 (genome-wide LDSC) and be held fixed. Proposition 3.3 shows they cannot be estimated jointly with $\mathbf{w}$ within a window.

---

#### S2. Matrix-normal and mediator models; fixed versus random effects

##### S2.1 Matrix-normal effects

$\mathbf{B} \sim \mathcal{MN}_{m\times q}(\mathbf{0}, \mathbf{U}, \mathbf{V})$ means $\operatorname{vec}\mathbf{B} \sim N(\mathbf{0}, \mathbf{V}\otimes\mathbf{U})$, with density

$$f(\mathbf{B}) = \frac{\exp\{-\frac{1}{2}\operatorname{tr}(\mathbf{V}^{-1}\mathbf{B}^\top\mathbf{U}^{-1}\mathbf{B})\}}{(2\pi)^{mq/2}|\mathbf{U}|^{q/2}|\mathbf{V}|^{m/2}}$$

(Dawid 1981; Gupta & Nagar 2000). Its moments are $E[\boldsymbol{\beta}_a\boldsymbol{\beta}_b^\top] = V_{ab}\mathbf{U}$, $E[\mathbf{B}\mathbf{B}^\top] = \operatorname{tr}(\mathbf{V})\mathbf{U}$ and $E[\mathbf{B}^\top\mathbf{B}] = \operatorname{tr}(\mathbf{U})\mathbf{V}$.

So (A2) holds with $\tilde{\boldsymbol{\Sigma}} = m\mathbf{U}/\operatorname{tr}\mathbf{U}$ and $h_{ab} = V_{ab}\operatorname{tr}\mathbf{U}$. The pair $(\mathbf{U}, \mathbf{V})$ is identified only up to $(\kappa\mathbf{U}, \mathbf{V}/\kappa)$. Fixing $\operatorname{tr}\tilde{\boldsymbol{\Sigma}} = m$ is the normalisation $\operatorname{avg}\mathbf{w} = 1$.

A point (spike-and-slab) matrix normal adds sparsity: $\mathbf{B} = \boldsymbol{\Delta}\circ\mathbf{G}$ with $\mathbf{G}$ matrix normal and $\boldsymbol{\Delta}$ a 0/1 mask. If the mask is independent of $\mathbf{G}$ and has a separable second moment, $E[\delta_{ia}\delta_{jb}] = \pi_{ij}^U\pi_{ab}^V$, then (A2) still holds with $\mathbf{U}\circ\boldsymbol{\pi}^U$ and $\mathbf{V}\circ\boldsymbol{\pi}^V$. Theorem 1 therefore applies unchanged; only fourth moments differ (§S5).

*Caution.* A per-SNP mask shared across traits, $\delta_{ia} = \delta_i$, multiplies the off-diagonal $\tilde{\Sigma}_{ij}$ by $E[\delta_i\delta_j]/\sqrt{E\delta_i E\delta_j}$. For independent masks that factor is $\pi$, so simulations must draw the masks with the intended co-causality.

##### S2.2 Simulation generator

Given a SNP correlation template $\mathbf{U}_0$ (unit diagonal, encodes $\mathbf{R}_\beta$), a trait correlation template $\mathbf{V}_0$ (encodes $r_g$ ), enrichments $\mathbf{w}$ with mean 1, and heritabilities $h_a^2$:

1. $\mathbf{U} = \mathbf{W}^{1/2}\mathbf{U}_0\mathbf{W}^{1/2}$, so $\operatorname{tr}\mathbf{U} = m$.
2. $\mathbf{V} = \operatorname{Diag}(\mathbf{h}/\sqrt{m})\,\mathbf{V}_0\,\operatorname{Diag}(\mathbf{h}/\sqrt{m})$, so $V_{ab} = h_{ab}/m$.
3. $\mathbf{B} = \mathbf{A}\mathbf{G}\mathbf{C}^\top$ with $\mathbf{A}\mathbf{A}^\top = \mathbf{U}$, $\mathbf{C}\mathbf{C}^\top = \mathbf{V}$, and $\mathbf{G}$ having iid unit-variance entries.

Then $\operatorname{Cov}(\operatorname{vec}\mathbf{B}) = \mathbf{V}\otimes\mathbf{U}$ for *any* iid $\mathbf{G}$. Gaussian $\mathbf{G}$ gives the matrix normal. Point-normal $\mathbf{G}$ gives sparse, heavy-tailed effects with the same second moments; check M1 uses this. The correlation structures of $\mathbf{B}$ equal $\mathbf{U}_0$ and $\mathbf{V}_0$ in expectation.

##### S2.3 Mediators

Let $L$ latent mediators carry the genetic effects: $\mathbf{m} = \mathbf{P}\mathbf{x} + \mathbf{e}$ and $\mathbf{y} = \mathbf{Q}\mathbf{m} + \mathbf{u}$, so $\mathbf{B}^\top = \mathbf{Q}\mathbf{P}$, i.e. $\mathbf{B} = \sum_{r=1}^L \mathbf{p}_r\mathbf{q}_r^\top$. Here $\mathbf{p}_r \in \mathbb{R}^m$ holds the SNP effects on mediator $r$ and $\mathbf{q}_r \in \mathbb{R}^q$ the mediator's effects on traits.

**Proposition 2.3.** Let the $\mathbf{p}_r$ and $\mathbf{q}_r$ be mutually independent with mean zero, $\mathrm{Cov}(\mathbf{p}_r) = \mathbf{U}_r$ and $\mathrm{Cov}(\mathbf{q}_r) = \mathbf{V}_r$.

- a. One mediator: $\mathrm{Cov}(\mathrm{vec}\,\mathbf{B}) = \mathbf{V}_1 \otimes \mathbf{U}_1$, the matrix-normal second-moment structure. $\mathbf{B}$ is **not** matrix normal, however. If $\mathbf{p}, \mathbf{q}$ are Gaussian, $E[B_{ia}^4] = 9U_{ii}^2 V_{aa}^2$, whereas the matrix normal gives $3U_{ii}^2 V_{aa}^2$ (check M3: 35.8 vs theory 36).
- b. $L$ mediators: $\mathrm{Cov}(\mathrm{vec}\,\mathbf{B}) = \sum_r \mathbf{V}_r \otimes \mathbf{U}_r$, a sum of Kronecker products. It is separable, so (A2) holds, iff all $\mathbf{U}_r$ are proportional or all $\mathbf{V}_r$ are proportional.
- c. As $L \to \infty$ with no mediator dominating, $\mathrm{vec}\,\mathbf{B}$ is asymptotically Gaussian by the Lindeberg CLT.

*Proof.* $\mathrm{Cov}(B_{ia}, B_{jb}) = \sum_{r,r'} E[p_{ir} q_{ar} p_{jr'} q_{br'}]$. Independence across $r \neq r'$ and zero means remove the cross terms, leaving $\sum_r U_{r,ij} V_{r,ab}$. The fourth moment: $E[p^4 q^4] = 3U^2 \cdot 3V^2$. (c) is the CLT for sums of independent non-identical terms. □

**Remark.** One mediator reproduces the matrix-normal second moments, which is all Theorem 1 uses, but not its higher moments. Under (b) without separability, (1.2) becomes $E[\mathbf{z}_a \mathbf{z}_b^\top] = c_{ab}\mathbf{R} + \sqrt{n_a n_b} \sum_r V_{r,ab} \mathbf{R} \mathbf{U}_r \mathbf{R}$. LRCQ then targets a trait-weighted mixture of the mediator-specific patterns (§S3.2).

#### S2.4 Fixed-effect view and the limiting genetic-effect correlation

Treat the $q$ traits as a sample from a population of phenotypes. The empirical SNP-wise covariance is $\hat{\boldsymbol{\Sigma}}_\beta^{(q)} = \mathbf{B}\mathbf{B}^\top / q$ (a centred version differs by $O(1/q)$ under (A1)). The empirical genetic-effect correlation is $\hat{\rho}_{ij}^{(q)} = \frac{\boldsymbol{\beta}_{i\cdot}^\top \boldsymbol{\beta}_{j\cdot}}{\|\boldsymbol{\beta}_{i\cdot}\|\,\|\boldsymbol{\beta}_{j\cdot}\|}$.

- *Limiting (trait-free) correlation.* If the columns $\boldsymbol{\beta}_a / \sqrt{h_a^2}$ are exchangeable with finite fourth moments and weakly dependent across traits, then $\hat{\boldsymbol{\Sigma}}_\beta^{(q)} - \boldsymbol{\Sigma}_\beta \to \mathbf{0}$ and $\hat{\rho}_{ij}^{(q)} \to \rho_{ij}$ almost surely as $q \to \infty$ (SLLN and continuous mapping). This separates a trait-set-specific correlation $\hat{\rho}^{(q)}$ from a trait-free one $\rho$.
- *Finite-$q$ variability.* Under matrix normality, $\mathrm{Var}(\hat{\rho}_{ij}^{(q)}) \approx (1 - \rho_{ij}^2)^2 / q_{\text{eff}}$ with $q_{\text{eff}} = (\mathrm{tr}\,\mathbf{V})^2 / \mathrm{tr}(\mathbf{V}^2)$. This follows from $\mathrm{Var}(\sum_a \beta_{ia}\beta_{ja}) = \mathrm{tr}(\mathbf{V}^2)(U_{ii}U_{jj} + U_{ij}^2)$ and the delta method. Genetically correlated traits shrink $q_{\text{eff}}$ below $q$. With unrelated traits $q_{\text{eff}} = q^2 \bar{h}^4 / \sum_a h_a^4 \leq q$.
- The fixed-effect view's empirical $\hat{\rho}^{(q)}$ is a parameter of the realised $\mathbf{B}$. The random-effect view's $\rho$ is its limit. Both LRCP estimators below estimate the genetic-effect *covariance* without bias under (A2). The correlation estimate divides by estimated variances, so it is a ratio estimator: it is consistent when the variance estimates are positive and stable, but it carries a finite-sample ratio bias, and $|\hat{\rho}|$ can exceed 1 when they are imprecise. For the realised trait set they estimate an $s$-weighted empirical version (cf. §S3.2).

---

#### S3. LRCQ: estimating $\mathbf{w}$

Throughout §S3, $c_{aa}$ and $s_a$ come from Stages 1–2 and $\mathbf{C} = \mathbf{0}$ is the working model (Proposition 1.6). The data are the $m \times q$ matrix $\mathbf{Y}^{(2)}$ with columns $\mathbf{z}_a^{\circ 2} - c_{aa}\mathbf{1}$.

##### S3.1 Fast closed-form OLS

The stacked regression is $\operatorname{vec}\mathbf{Y}^{(2)} = \mathbf{X}\mathbf{w} + \boldsymbol{\varepsilon}$ with $\mathbf{X} = \mathbf{s} \otimes \mathbf{D}$ (the $q$ blocks $s_a\mathbf{D}$ stacked).

**Theorem 3.1.** If $\mathbf{D}$ is invertible,

$$\hat{\mathbf{w}}_{\text{OLS}} = (\mathbf{X}^\top\mathbf{X})^{-1}\mathbf{X}^\top \operatorname{vec}\mathbf{Y}^{(2)} = \frac{1}{\|\mathbf{s}\|^2}\,\mathbf{D}^{-1}\mathbf{Y}^{(2)}\mathbf{s} = \frac{1}{\|\mathbf{s}\|^2}\,\mathbf{D}^{-1}\sum_{a=1}^{q} s_a(\mathbf{z}_a^{\circ 2} - c_{aa}\mathbf{1}) \quad (3.1)$$

*Proof.* $\mathbf{X}^\top\mathbf{X} = (\mathbf{s}^\top\mathbf{s}) \otimes \mathbf{D}^\top\mathbf{D} = \|\mathbf{s}\|^2\mathbf{D}^2$, since $\mathbf{D}$ is symmetric. $\mathbf{X}^\top \operatorname{vec}\mathbf{Y}^{(2)} = \sum_a s_a\mathbf{D}(\mathbf{z}_a^{\circ 2} - c_{aa}\mathbf{1}) = \mathbf{D}\mathbf{Y}^{(2)}\mathbf{s}$. Hence $\hat{\mathbf{w}} = \|\mathbf{s}\|^{-2}\mathbf{D}^{-2}\mathbf{D}\mathbf{Y}^{(2)}\mathbf{s}$. □

This holds for any $m$ and $q$. The cost is one $m \times m$ solve plus an $O(mq)$ product, independent of $q$ apart from that product. $\mathbf{D}^{-1}$ (or its factorisation) can be precomputed per LD block and population. Unbiasedness, $E\hat{\mathbf{w}} = \mathbf{w}$, follows from (1.4) with $\mathbf{C} = \mathbf{0}$. In check L1, fast and brute-force OLS agree to $10^{-14}$.

*Note.* $\mathbf{D}$ is positive semi-definite by the Schur product theorem, and positive definite whenever $\mathbf{R}$ is. Even when a reference-panel $\hat{\mathbf{R}}$ is singular ($m > n_{\text{ref}}$), $\operatorname{rank}(\hat{\mathbf{R}} \circ \hat{\mathbf{R}}) \le r(r+1)/2$ for $r = \operatorname{rank}\hat{\mathbf{R}}$, so $\hat{\mathbf{D}}$ is often full rank but ill-conditioned (§S3.6).

##### S3.2 Heterogeneous enrichment across traits: what OLS targets

Drop (A2) and let trait $a$ have its own enrichment $\mathbf{w}_a$, still with $\mathbf{C} = \mathbf{0}$.

**Proposition 3.2.**

$$E[\hat{\mathbf{w}}_{\text{OLS}}] = \sum_a \frac{s_a^2}{\|\mathbf{s}\|^2}\,\mathbf{w}_a.$$

OLS LRCQ therefore estimates a power-weighted average of trait-specific enrichments, with weights $\propto (n_a h_a^2)^2$. The equal-weight alternative

$$\hat{\mathbf{w}}_{\text{eq}} = \frac{1}{q}\sum_a \frac{1}{s_a}\mathbf{D}^{-1}(\mathbf{z}_a^{\circ 2} - c_{aa}\mathbf{1})$$

is unbiased for $\bar{\mathbf{w}} = q^{-1}\sum_a \mathbf{w}_a$ but has larger variance.

*Proof.* $E[\mathbf{z}_a^{\circ 2} - c_{aa}\mathbf{1}] = s_a\mathbf{D}\mathbf{w}_a$; substitute into (3.1). □

In check L4, OLS matches the $s^2$-weighted target (correlation 1.000). It differs from the simple mean by up to 0.80 in enrichment units. **In practice,** a large-$n$, high-$h^2$ GWAS dominates $\hat{\mathbf{w}}$. Report which weighting is used. When the trait collection is unbalanced, for example many anthropometric traits in Pan-UKB, also report $\hat{\mathbf{w}}_{\text{eq}}$

or phenotype-category-specific estimates (§S7.4). The WLS estimator weights traits differently again, roughly $\propto s_a^2/(c+s_a\lambda_k)^2$.

Read as "the average over traits of the per-SNP enrichment", $\mathbf{w}$ is the estimand of $\hat{\mathbf{w}}_{\text{eq}}$. OLS targets the $s_a^2$-weighted average, and the two agree only when enrichment is the same in every trait.

##### S3.3 Intercepts cannot be fitted per window

**Proposition 3.3.** Consider $E[\mathbf{z}_a^{\circ 2}] = c_a\mathbf{1} + s_a\mathbf{D}\mathbf{w}$ with free intercepts $c_1,\ldots,c_q$. The stacked design $[\mathbf{s}\otimes\mathbf{D},\ \mathbf{I}_q\otimes\mathbf{1}_m]$ has rank $m+q-1$. Its null space is spanned by $(\mathbf{D}^{-1}\mathbf{1},\ -\mathbf{s})$. Hence $(\mathbf{w}+\tau\mathbf{D}^{-1}\mathbf{1},\ c_a-\tau s_a)$ fit equally well for every $\tau$, whatever the $s_a$. With a single common intercept $c$ the direction is different: two parameter sets fit equally well only if $s_a\mathbf{D}\Delta\mathbf{w} + \Delta c\,\mathbf{1} = \mathbf{0}$ for every $a$, and subtracting two traits with $s_a \neq s_b$ gives $\Delta\mathbf{w} = \mathbf{0}$, hence $\Delta c = 0$. A common intercept is therefore formally identified when the $s_a$ differ, but the fit is ill-conditioned when they are similar, and a common intercept is not a realistic model for traits with different stratification or sample overlap.

*Proof.* $(\mathbf{s}\otimes\mathbf{D})\mathbf{D}^{-1}\mathbf{1} = \mathbf{s}\otimes\mathbf{1} = (\mathbf{I}_q\otimes\mathbf{1})\mathbf{s}$. □

Check L5 confirms the rank deficit of exactly one for free intercepts. Genome-wide LDSC identifies intercepts only because it constrains $\mathbf{w}\equiv\mathbf{1}$. **In practice,** take $c_{aa}$ (and $c_{ab}$) from Stage 2 and fix them in Stages 3–4. A free intercept per window cannot absorb population stratification. It does not need to: under the usual model, stratification adds terms that are constant across SNPs (Yengo et al. 2018), and the genome-wide Stage-2 intercept already contains them.

##### S3.4 Variances of squared and cross-product Z-scores

If $\mathbf{z}$ is jointly Gaussian with $\sigma_{ka,lb} = E[z_{ka}z_{lb}]$ given by (1.2), Isserlis' theorem (Isserlis 1918) gives:

| quantity | value |
|---|---|
| $\mathrm{Var}(z_{ka}^2)$ | $2\sigma_{ka}^4$ |
| $\mathrm{Cov}(z_{ka}^2, z_{lb}^2)$ | $2\sigma_{ka,lb}^2$ |
| $\mathrm{Var}(z_{ka}z_{lb})$ | $\sigma_{ka}^2\sigma_{lb}^2 + \sigma_{ka,lb}^2$ |
| $\mathrm{Cov}(z_{ka}z_{lb}, z_{k'c}z_{l'd})$ | $\sigma_{ka,k'c}\sigma_{lb,l'd} + \sigma_{ka,l'd}\sigma_{lb,k'c}$ |
| $\mathrm{Cov}(z_{ka}^2, z_{ka}z_{lb})$ | $2\sigma_{ka}^2\sigma_{ka,lb}$ |

Check M2 confirms these to 0.5%. Effects are not Gaussian in practice; §S5 gives the non-Gaussian correction.

##### S3.5 Fast WLS and IRLS

Let $v_{ka} = \mathrm{Var}(z_{ka}^2)$ and $\mathbf{W}_a = \mathrm{Diag}(1/v_{1a},\ldots,1/v_{ma})$. Then

$$\hat{\mathbf{w}}_{\text{WLS}} = \Big(\sum_a s_a^2\,\mathbf{D}\mathbf{W}_a\mathbf{D}\Big)^{-1}\sum_a s_a\,\mathbf{D}\mathbf{W}_a(\mathbf{z}_a^{\circ 2} - c_{aa}\mathbf{1}). \tag{3.2}$$

*Proof.* $\mathbf{X}^\top\mathbf{W}\mathbf{X} = \sum_a(s_a\mathbf{D})\mathbf{W}_a(s_a\mathbf{D})$ and $\mathbf{X}^\top\mathbf{W}\mathbf{y} = \sum_a s_a\mathbf{D}\mathbf{W}_a\mathbf{y}_a$, since $\mathbf{W}$ is block diagonal over traits. □

The normal matrix $\sum_a s_a^2 \mathbf{D}\mathbf{W}_a\mathbf{D}$ does not collapse to a single $\mathbf{D}^{-1}$ because the weights vary over SNPs and traits. Building it costs $O(qm^2)$.

- **IRLS:** iterate $v_{ka}^{(t)} = 2\big(c_{aa} + s_a(\mathbf{D}\hat{\mathbf{w}}^{(t)})_k\big)^2$, flooring $\hat{\mathbf{w}}^{(t)}$ at 0 inside the weights, as LDSC floors its weights.
- A single reweighting step (OLS followed by one WLS step) is a reasonable compromise. Correctly specified weights give the efficient linear estimator (Aitken 1935). Weights from a poor first-stage estimate can be worse than equal weights in small $q$.
- WLS ignores the correlation of $z_{ka}^2$ and $z_{la}^2$ through LD. Full GLS would need $\mathrm{Cov}(\mathbf{z}_a^{\circ 2}) = 2(\mathbf{\Sigma}_a \circ \mathbf{\Sigma}_a)$ with $\mathbf{\Sigma}_a = c_{aa}\mathbf{R} + s_a\mathbf{RWR}$. Its cost is rarely justified, and we do not use it.

In check L2, fast and brute-force WLS agree to $10^{-13}$.

##### S3.6 Standard errors and conditioning

**Analytic sandwich (OLS).** For traits without sample overlap and $h_{ab} = 0$,

$$\mathrm{Var}(\hat{\mathbf{w}}_{\mathrm{OLS}}) = \frac{1}{\|\mathbf{s}\|^4}\,\mathbf{D}^{-1}\Big[\sum_a s_a^2\,\mathrm{Cov}(\mathbf{z}_a^{\circ 2})\Big]\mathbf{D}^{-1}, \qquad \mathrm{Cov}(\mathbf{z}_a^{\circ 2}) = 2\,\mathbf{\Sigma}_a \circ \mathbf{\Sigma}_a + \mathbf{K}_a \tag{3.3}$$

where $\mathbf{K}_a$ is the non-Gaussian term of §S5. With overlap or genetic correlation, add $\sum_{a\neq b} s_a s_b\, 2(\mathbf{\Sigma}_{ab} \circ \mathbf{\Sigma}_{ab})$. Theorem 3.10 (§S3.10) gives the exact cross-trait variance in a cheap closed form, and that is the formula to use in practice.

**Jackknife over GWAS: not recommended.** A delete-one-trait-cluster jackknife treats clusters as independent replicates. It therefore estimates only the within-cluster part of $\mathbf{s}^\top(\mathbf{\Gamma} \circ \mathbf{\Gamma})\mathbf{s}$, and misses the between-cluster covariance that sample overlap creates. In a biobank every pair of traits shares samples, so no clustering yields independent clusters. In check J1 (`check_jackknife_overlap.py`: one null SNP, $q = 300$ in 20 clusters, overlap correlation 0.4 within and $r_b$ between clusters):

- At $r_b = 0$, the jackknife median SE is 3.10 against an empirical SD of 3.52.
- At $r_b = 0.1$, 2.94 against 4.84.
- At $r_b = 0.2$, 2.55 against 7.31.
- The model SE (3.5) matched the empirical SD in all three settings (3.51, 4.75, 7.30).

Simulations with UK Biobank-like overlap (LRCQ paper) show the same: the SD of standardised errors is 2.3–2.7 for the jackknife, against 1.03–1.12 for (3.5). **Use the model SEs of Theorem 3.10, with the non-Gaussian term of §S5 where needed.** Use the jackknife only as a diagnostic, and only for trait sets from disjoint samples. The block jackknife over SNPs used by LDSC is *not* appropriate here, because each $w_i$ is SNP-specific.

**Back-of-envelope power.** Without LD and under $w_i = 0$, $\mathrm{SE}(\hat{w}_i) \approx \sqrt{2}/\|\mathbf{s}\|$. For example, with $q = 500$ effectively independent GWAS, $n = 4 \times 10^5$, $h^2 = 0.1$ and $m = 1.2 \times 10^6$: $s = 0.033$, $\|\mathbf{s}\| = 0.75$, $\mathrm{SE} \approx 1.9$. LRCQ therefore detects SNPs with enrichment of order 5–10 or more ("omni-SNPs") at nominal levels. At genome-wide thresholds, with realistic $q_{\mathrm{eff}}$ and LD, the requirement is nearer $w \gtrsim 20$–30 (§S9.2). LD inflates the SE by roughly $\sqrt{(\mathbf{D}^{-1})_{ii}}$-type factors. Genetic correlation among traits reduces the effective $q$ (§S2.4).

**Conditioning of $\mathbf{D}$.**

- i. Use the bias-adjusted $\tilde{r}^2 = \hat{r}^2 - (1-\hat{r}^2)/(n_{\text{ref}}-2)$, as in LDSC, because $E\hat{r}^2 \approx r^2 + (1-r^2)^2/n_{\text{ref}}$.
- ii. Replace $\mathbf{D}^{-1}$ by a ridge or truncated-eigen inverse. A ridge $\mathbf{D}_\lambda = \mathbf{D} + \lambda\mathbf{I}$ gives $E\hat{\mathbf{w}} = \mathbf{D}_\lambda^{-1}\mathbf{D}\mathbf{w}$: a known, local smoothing of $\mathbf{w}$ over SNPs in high LD.
- iii. SNPs in near-perfect LD are not separately identifiable. Report $\hat{w}$ for LD clumps (sum over the clump), which is well determined.

#### S3.7 Sliding three-block window

Equation (1.4) for SNP $k$ involves only SNPs in LD with $k$. If LD blocks are approximately independent (Berisa & Pickrell 2016), the equations for SNPs in the middle block of a three-block window are exact. Those for SNPs in the outer blocks are missing LD partners outside the window. Keeping only middle-block estimates, and sliding one block at a time, therefore uses exactly-specified equations for every reported SNP. The outer blocks absorb edge effects through $\mathbf{D}^{-1}$, which decays with LD distance. At chromosome ends a two-block window is exact for the end block.

#### S3.8 Negative estimates: rectification methods A, B, C as symmetric-null thresholds

Under $w_i = 0$, $\hat{w}_i$ is a linear combination of many centred squared Z-scores. It is approximately symmetric about 0 by the CLT over traits. Write $N_-(t) = \#\{i : \hat{w}_i \le -t\}$ and $N_+(t) = \#\{i : \hat{w}_i \ge t\}$. Symmetry gives $E[\#\text{nulls} \ge t] \approx E[\#\text{nulls} \le -t]$, the target–decoy / knockoff principle (Elias & Gygi 2007; Barber & Candès 2015). Each method sets $\hat{w}_i \leftarrow 0$ for all $\hat{w}_i \le t^*$:

- **Method A**, $t^* = |\min_i \hat{w}_i|$. No negative estimate exceeds $t^*$ in magnitude, so the estimated number of false positives above $t^*$ is zero. This is the most conservative rule (estimated FDP $\approx 1/N_+(t^*)$).
- **Method B** zeroes all negatives plus the $N_-(0)$ smallest positives. It matches *counts*: about $2N_-(0)$ null SNPs, half of them positive.
- **Method C** chooses $t^*$ so that $\sum_{0<\hat{w}_i \le t^*} \hat{w}_i = \sum_{\hat{w}_i<0} |\hat{w}_i|$. It matches *mass*. **Property:** C is the unique threshold rule that preserves the total $\sum_i \hat{w}_i$, and hence the unbiasedness of the mean enrichment and the $\operatorname{avg}\mathbf{w} = 1$ normalisation, while making all estimates non-negative.

**Method D (FDR-controlled).** $t^*_\alpha = \min\{t > 0 : (1 + N_-(t))/\max(N_+(t), 1) \le \alpha\}$, the knockoff+ threshold. It gives approximate FDR control at level $\alpha$ for declaring $w_i > 0$. Exact control needs independence across SNPs, which $\mathbf{D}^{-1}$ induces only approximately. Its calibration under LD dependence needs to be checked by simulation before use.

*Rectified zeros are not evidence that $w_i = 0$.* At genome scale the null SD of a single $\hat{w}_i$ is large (§S9.2), so negative raw estimates are common (54% of clump estimates in the Pan-UKB analysis) and include truly enriched clumps. If negative estimates are used as "null" labels, for example to train an annotation model, apply a significance filter first.

#### S3.9 Using all within-block cross-products: the joint $\tilde{\boldsymbol{\Sigma}}$ estimator

By (1.2) with $a = b$, $E[\mathbf{z}_a\mathbf{z}_a^\top - c_{aa}\mathbf{R}] = s_a\mathbf{R}\tilde{\boldsymbol{\Sigma}}\mathbf{R}$. Hence

$$\hat{\tilde{\boldsymbol{\Sigma}}} = \frac{1}{\|\mathbf{s}\|^2}\mathbf{R}^{-1}\Big[\sum_a s_a(\mathbf{z}_a\mathbf{z}_a^\top - c_{aa}\mathbf{R})\Big]\mathbf{R}^{-1} = \frac{1}{\|\mathbf{s}\|^2}\mathbf{R}^{-1}\mathbf{Z}\mathbf{S}\mathbf{Z}^\top\mathbf{R}^{-1} - \frac{\sum_a s_a c_{aa}}{\|\mathbf{s}\|^2}\mathbf{R}^{-1} \tag{3.4}$$

is unbiased for the whole $\tilde{\boldsymbol{\Sigma}}$ within a block: its diagonal estimates $\mathbf{w}$ and its off-diagonal estimates $\mathbf{C}$, with no working assumption on local $\rho$. This answers Proposition 1.6: cross-products identify what squares cannot. The OLS fit of the stacked model $\operatorname{vec}(\mathbf{z}_a\mathbf{z}_a^\top)$ on $s_a(\mathbf{R}\otimes\mathbf{R})$ gives exactly (3.4). It is also the multi-trait analogue of the quadratic form $\mathbf{z}^\top\mathbf{R}^{-1}\mathbf{z}$ behind HESS local heritability (Shi et al. 2016). It needs a well-conditioned $\mathbf{R}$ (regularise as in §S3.6) and has larger variance than (3.1) for $\mathbf{w}$ when local $\rho$ is truly 0. In check P3, $\operatorname{diag}\hat{\tilde{\boldsymbol{\Sigma}}}$ recovers $w = 50$ and 1 without bias.

##### S3.10 Cross-trait dependence: exact variance of the Stage-3 and Stage-4 estimators

Traits are not independent replicates. Shared samples and phenotypic correlation (intercepts $c_{ab}$) and genetic correlation ($t_{ab}$) correlate Z-scores across GWAS. Collect the intercepts in $\boldsymbol{\Gamma} = (c_{ab})$ and let $\mathbf{G} = \mathbf{R}\tilde{\boldsymbol{\Sigma}}\mathbf{R}$. By Theorem 1,

$$\operatorname{Cov}(\operatorname{vec}\mathbf{Z}) = \boldsymbol{\Gamma}\otimes\mathbf{R} + \mathbf{T}\otimes\mathbf{G}.$$

Every linear estimator in §S3–S4 has the form $\hat{\theta} = K^{-1}\sum_a s_a\,\mathbf{z}_a^\top\mathbf{A}\mathbf{z}_a$ for a symmetric $m\times m$ matrix $\mathbf{A}$ :

- LRCQ OLS $\hat{w}_i$ (3.1): $\mathbf{A} = \operatorname{Diag}(\mathbf{D}^{-1}\mathbf{e}_i)$, $K = \|\mathbf{s}\|^2$.
- LRCP REG $\hat{C}_{ij}$ (4.2): $\mathbf{A} = \frac{1}{2}(\mathbf{a}\mathbf{b}^\top + \mathbf{b}\mathbf{a}^\top)$ with $\mathbf{a} = \mathbf{R}\mathbf{e}_i$, $\mathbf{b} = \mathbf{R}\mathbf{e}_j$, and $K = \|\mathbf{s}\|^2\ell_i\ell_j$.
- MOM (4.1) is the same with $\mathbf{R}^{-1}$ in place of $\mathbf{R}$.

The intercept offsets are constants and do not affect the variance.

**Theorem 3.10.** For Gaussian $\mathbf{Z}$,

$$\operatorname{Var}(\hat{\theta}) = \frac{2}{K^2}\Big[\tau_{RR}\,\mathbf{s}^\top(\boldsymbol{\Gamma}\circ\boldsymbol{\Gamma})\mathbf{s} + 2\,\tau_{RG}\,\mathbf{s}^\top(\boldsymbol{\Gamma}\circ\mathbf{T})\mathbf{s} + \tau_{GG}\,\mathbf{s}^\top(\mathbf{T}\circ\mathbf{T})\mathbf{s}\Big], \qquad \tau_{XY} = \operatorname{tr}(\mathbf{A}\,\text{[illegible]}) \tag{3.5}$$

*Proof.* For jointly Gaussian vectors, $\operatorname{Cov}(\mathbf{z}_a^\top\mathbf{A}\mathbf{z}_a, \mathbf{z}_b^\top\mathbf{A}\mathbf{z}_b) = 2\operatorname{tr}(\mathbf{A}\boldsymbol{\Sigma}_{ab}\mathbf{A}\boldsymbol{\Sigma}_{ba})$ (Isserlis). Substitute $\boldsymbol{\Sigma}_{ab} = c_{ab}\mathbf{R} + t_{ab}\mathbf{G}$, expand the trace, and sum over $a, b$ with weights $s_a s_b$. □

The cross-trait dependence enters only through three $q\times q$ quadratic forms. The SNP-level work is three traces, computed once per SNP or pair. For LRCQ $\hat{w}_i$ the traces simplify: with $\mathbf{d} = \mathbf{D}^{-1}\mathbf{e}_i$, $\tau_{RR} = \mathbf{d}^\top(\mathbf{R}\circ\mathbf{R})\mathbf{d} = (\mathbf{D}^{-1})_{ii}$, $\tau_{RG} = \mathbf{d}^\top(\mathbf{R}\circ\mathbf{G})\mathbf{d}$ and $\tau_{GG} = \mathbf{d}^\top(\mathbf{G}\circ\mathbf{G})\mathbf{d}$. Plug-in values come from Stage 1–2 ($\hat{h}_{ab}$, $\hat{c}_{ab}$) and Stage 3 ($\hat{\mathbf{w}}$ in $\mathbf{G}$; which form of $\hat{\mathbf{w}}$ to use is §S3.10a). Non-Gaussian effects add the cumulant term of §S5.

The *independence* variance is (3.5) with $\boldsymbol{\Gamma}$ and $\mathbf{T}$ replaced by their diagonals.

**Effective number of traits.** Define $q_{\text{eff}} = q\cdot\operatorname{Var}_{\text{indep}}/\operatorname{Var}$. When one term dominates, $q_{\text{eff}}$ has a closed form:

- *Null or background SNPs*, where the $\boldsymbol{\Gamma}$ term dominates: $q_{\text{eff}} = q\,\mathbf{s}^\top\operatorname{Diag}(\boldsymbol{\Gamma}\circ\boldsymbol{\Gamma})\mathbf{s}/\mathbf{s}^\top(\boldsymbol{\Gamma}\circ\boldsymbol{\Gamma})\mathbf{s}$. With equal $s_a$ and $c_{aa} = 1$ this is $q^2/\sum_{a,b}c_{ab}^2 = (\operatorname{tr}\boldsymbol{\Gamma})^2/\operatorname{tr}(\boldsymbol{\Gamma}^2)$.

- *Strongly enriched SNPs*, where the $\mathbf{T}$ term dominates: the same formula with $\mathbf{T}$, i.e. driven by genetic correlations, $q^2/\sum_{a,b} r_{g,ab}^2$ when $s_a$ are equal.

The single-matrix form $q_{\text{eff}} = \text{tr}(\mathbf{M})^2/\text{tr}(\mathbf{M}^2)$ is exact only when the per-SNP cross-trait covariance $\mathbf{\Sigma}_{ab}$ is proportional, across trait pairs, to one SNP-space matrix. That holds when one term dominates, or when $c_{ab}/c_{aa} \approx t_{ab}/\sqrt{t_{aa}t_{bb}}$ (phenotypic correlation in the overlap ≈ genetic correlation). In between, use (3.5). It costs the same and is exact. For the LRCP Stage-4 test of a pair $(k, l)$, (3.5) with $\mathbf{A} = \frac{1}{2}(\mathbf{e}_k\mathbf{e}_l^\top + \mathbf{e}_l\mathbf{e}_k^\top)$ gives the cross-trait-correct null variance of $\sum_a s_a z_{ka} z_{la}$. With $\mathbf{M}_k = (\Sigma_{ab,kk})_{ab}$ it reduces to $\text{Cov}(z_{ka}z_{la}, z_{kb}z_{lb}) = M_{k,ab}M_{l,ab} + \Sigma_{ab,kl}\Sigma_{ab,lk}$.

**Check X1** (`check_crosstrait.py`): $q = 120$ traits in 4 clusters ($r_g$ 0.6 within, 0.1 between), full sample overlap with phenotypic correlation $0.5r_g + 0.1$, an enriched correlated pair.

| estimator | empirical SD | (3.5) | independence formula | implied $q_{\text{eff}}$ |
|---|---|---|---|---|
| LRCQ $\hat{w}_i$ | 18.6 | 18.3 | 6.7 | 16 |
| LRCP $\hat{C}_{ij}$ | 14.8 | 14.7 | 5.3 | 16 |

The independence formula understates SDs about 2.8-fold here. In the simulated trait structures of the LRCP paper $q_{\text{eff}}$ ranges from 16 to 120 for 300 traits, and in Pan-UKB it is 58 for the sampling-noise correlation $\mathbf{\Gamma}$ of 452 traits (null and background SNPs) but only about 11 for the genetic covariance $\mathbf{T}$ of 444 traits (strongly enriched SNPs); independence-based SEs then inflate type-I error. **Default:** analytic SEs from (3.5). Never use independence-based SEs. The trait-cluster jackknife is not a valid alternative under sample overlap (§S3.6, check J1).

##### S3.10a Which enrichment estimate to plug into $\mathbf{G}$

The terms $\tau_{RG}$ and $\tau_{GG}$ need $\mathbf{G} = \mathbf{R}\,\text{Diag}(\mathbf{w})\mathbf{R}$, and $\mathbf{w}$ must be estimated from the same GWAS. Three natural plug-ins fail in opposite directions:

- *Per-SNP clipping*, $\max(\hat{w}_i, 0)$, is conservative. Sampling noise in $\hat{w}$ enters $\tau_{GG}$ quadratically, so the plug-in overstates $\mathbf{G} \circ \mathbf{G}$ (§S7.4a); null z-scores had SD 0.4–0.8 in every design below.
- *Rectification method A* (§S3.8), which sets every $\hat{w}_i \leq |\min \hat{w}|$ to 0, is anti-conservative when the background is dense. It removes the polygenic background from $\mathbf{G}$, and the omitted part of $\tau_{RG}$ and $\tau_{GG}$ grows with $s\,\ell$, the per-SNP signal times the LD score.
- *A local second moment* of $\hat{w}$ is very conservative (SD about 0.5) for the same reason as clipping.

**Default.** Keep the estimates that survive method A, and give every SNP that method A sets to zero the mean of $\hat{w}$ over that zeroed set, floored at 0. This background-filled plug-in removes the noise of single estimates but keeps the average background. A plain window mean of $\hat{w}$ (floored at 0) for all SNPs behaves similarly. The background-filled plug-in is the default in lrcpq 0.3.1 (`se_rectify = "bgA"`). For the test of $w_i > 1$ on tag-set clumps (§S3.11), the null value is the tag estimand under $\mathbf{w} \equiv \mathbf{1}$, $(\mathbf{D}_T^\top\mathbf{D}_T)^{-1}\mathbf{D}_T^\top\mathbf{D}\mathbf{1}$, not 1.

**Remark (the reported SE is not the test variance).** With this plug-in, a tag whose own estimate is kept contributes its own $\hat{W}_t^2$ to $\tau_{GG}$. For a large estimate the variance tends to $2\hat{W}_t^2\,\mathbf{s}^\top(\mathbf{T} \circ \mathbf{T})\mathbf{s}/\|\mathbf{s}\|^4$, so the ratio $\hat{W}_t/\widehat{\text{SE}}$ is bounded near $\|\mathbf{s}\|^2/\{2\,\mathbf{s}^\top(\mathbf{T} \circ \mathbf{T})\mathbf{s}\}^{1/2}$ whatever the size of $\hat{W}_t$ (about 1.7 for the 444 Pan-UKB traits). This SE is the right measure of how precisely $W_t$ is estimated: with few effective traits a large

$\hat{W}_t$ behaves like a variance estimated on few degrees of freedom, so it is reported with every estimate as $\hat{W}_t \pm \widehat{\mathrm{SE}}$. It is not used for testing, because a test built on it would have almost no power for the most enriched tags. Significance is assessed only with the score-type test of §S3.10b, whose variance is evaluated under the null hypothesis.

**Check P4** (`check_se_plugin.py`): 4 LD blocks of 50 SNPs with AR(1) LD, $q = 50$ traits in 5 genetic clusters ( $r_g$ 0.5 within, 0.1 between), full sample overlap with phenotypic correlation 0.15, equal $s_a = s$, 400 replicates. Entries are the SD of $(\hat{w}_i - w_i)/\widehat{\mathrm{SE}}$ over SNPs with background $w_i$ (1 is calibrated). Sparse: 20 SNPs at $w = 10$, rest 0; very sparse: 4 at $w = 50$; mixed: background 0.5 plus 4 at $w = 25.5$.

| mean LD score | design | $s$ | true $\mathbf{w}$ | method A | background-filled A | window mean | clipped |
|---|---|---|---|---|---|---|---|
| 2.8 | dense ($w \equiv 1$) | 0.076 | 1.00 | 0.93 | — | 1.01 | <0.8 |
| 8.6 | dense | 0.076 | 1.00 | 1.11 | 0.95 | 1.00 | <0.8 |
| 8.6 | dense | 0.2 | 1.00 | 1.38 | 0.93 | — | <0.8 |
| 15.7 | dense | 0.076 | 1.00 | 1.25 | 0.99 | 1.00 | <0.8 |
| 15.7 | dense | 0.2 | 1.00 | 1.68 | 0.97 | — | <0.8 |
| 15.7 | sparse | 0.2 | 1.00 | 1.60 | 0.97 | 0.97 | <0.8 |
| 15.7 | very sparse | 0.2 | 1.00 | 1.09 | 0.97 | 0.99 | <0.8 |
| 15.7 | mixed | 0.2 | 1.00 | 1.55 | 1.02 | 0.99 | <0.8 |
| 2.8 | very sparse | 0.2 | 1.00 | 0.92 | 0.92 | 0.89 | <0.8 |
| 2.8 | mixed | 0.2 | 1.00 | 0.99 | 0.90 | 0.95 | <0.8 |

With true $\mathbf{w}$ the SD was 1.00–1.02 throughout. Method A is adequate only when $s\,\ell$ is small or enrichment is very sparse; at $s\ell \approx 3$ its null type-I error at 5% was 0.19. The background-filled plug-in stays within 0.90–1.02 in every design, and errs on the conservative side.

##### S3.10b Testing tag enrichment: score-type variance and moment-matched p-values

**Statistic as a quadratic form.** For tag $t$ of the tag-set estimator (3.6), let $\mathbf{A}_t = \mathrm{Diag}(\mathbf{L}^\top \mathbf{e}_t)$ and $\mathbf{S} = \mathrm{Diag}(\mathbf{s})$. Stacking $\mathbf{z} = \mathrm{vec}\,\mathbf{Z}$,

$$\hat{W}_t = \mathbf{z}^\top \mathbf{M}_t \mathbf{z} - \mathrm{const}, \qquad \mathbf{M}_t = \|\mathbf{s}\|^{-2}\,\mathbf{S} \otimes \mathbf{A}_t.$$

For Gaussian $\mathbf{z}$ with covariance $\mathbf{\Sigma} = \mathbf{\Gamma} \otimes \mathbf{R} + \mathbf{T} \otimes \mathbf{G}$, the cumulants of the quadratic form are

$$\kappa_r = 2^{r-1}(r-1)!\ \mathrm{tr}\{(\mathbf{M}_t \mathbf{\Sigma})^r\}. \tag{3.5a}$$

$\kappa_2$ is (3.5). Since $\mathbf{M}_t\mathbf{\Sigma} = \|\mathbf{s}\|^{-2}\{(\mathbf{S\Gamma}) \otimes (\mathbf{A}_t\mathbf{R}) + (\mathbf{ST}) \otimes (\mathbf{A}_t\mathbf{G})\}$ and $\mathrm{tr}(\mathbf{X} \otimes \mathbf{Y}) = \mathrm{tr}\,\mathbf{X}\,\mathrm{tr}\,\mathbf{Y}$, each power expands into products of one trait-side and one SNP-side trace. Up to cyclic rotation there are four distinct words in $\{\mathbf{\Gamma}, \mathbf{T}\}$ of length 3 and six of length 4. So $\kappa_3$ needs four trait-side traces, such as $\mathrm{tr}\{(\mathbf{S\Gamma})^3\}$, computed once, and four SNP-side traces per tag, such as $\mathrm{tr}\{(\mathbf{A}_t\mathbf{R})^3\}$. $\kappa_4$ needs six of each.

**Score-type test.** We test $H_0 : W_t \le W_t^0$ (no more enriched than under uniform enrichment, with $W_t^0$ the tag estimand under $\mathbf{w} \equiv \mathbf{1}$ of §S3.10a) and $H_0 : W_t \le 0$. The test evaluates (3.5a) under the null. The SNPs of tag $t$ get the plug-in value that makes $W_t$ equal to its null value ($W_t^0$ or 0), and every other SNP keeps the background-filled plug-in of §S3.10a. The null configuration $\mathbf{w}^{(0)}$ must itself be a valid model, which needs two conditions. (i) Its implied mean for the tag equals the null value $W_t^{\text{null}}$ ($W_t^0$ or 0) in the estimator's own mean model: $\mathbf{a}_t^\top \mathbf{w}^{(0)} = W_t^{\text{null}}$, with $\mathbf{a}_t$ row $t$ of $(\mathbf{D}_{\cdot\mathcal{T}}^\top \mathbf{D}_{\cdot\mathcal{T}})^{-1}\mathbf{D}_{\cdot\mathcal{T}}^\top \mathbf{D}$. (ii) It is nonnegative, so that the genetic part $\mathbf{R}\,\mathrm{Diag}(\mathbf{w}^{(0)})\mathbf{R}$ of $\mathbf{\Sigma}$ in (3.5a) is positive semidefinite. When $W_t^{\text{null}} \geq 0$, setting the tag's entries to the null value meets both. When $W_t^0 < 0$ (§S3.11) it would make the genetic part indefinite. We then use the nonnegative vector closest in Euclidean distance to the background-filled plug-in $\bar{\mathbf{w}}$ subject to (i):

$$\mathbf{w}^{(0)} = (\bar{\mathbf{w}} - \lambda \mathbf{a}_t)_+, \qquad \lambda \text{ the root of } \mathbf{a}_t^\top (\bar{\mathbf{w}} - \lambda \mathbf{a}_t)_+ = W_t^0.$$

This is the Karush–Kuhn–Tucker solution of the projection. The left side of the root equation is non-increasing in $\lambda$, so the root is found by bisection. The mean in (i) is stated with $\mathbf{D}$ built from the bias-adjusted $\tilde{r}^2$ of §S3.6, not with $\hat{\mathbf{R}} \circ \hat{\mathbf{R}}$. $\tilde{r}^2$ is unbiased for the population $r^2$, which is what the estimator's mean involves. $\hat{r}^2$ from $n_{\text{ref}}$ individuals adds about $1/(n_{\text{ref}} - 2)$ for every unlinked pair, so a constraint stated in $\hat{\mathbf{R}} \circ \hat{\mathbf{R}}$ would treat reference-panel noise as LD. In Pan-UKB ($n_{\text{ref}} = 503$) such a constraint moves the null configuration far from the plug-in: the median ratio of the resulting test SE to that of the configuration with the tag simply set to its null value (no nonnegativity fix) is 1.25, and the 99th percentile is 7 (LRCQ paper, Supplementary Table S8). This null-evaluated variance is used only for the test; the SE reported with $\hat{W}_t$ is the one evaluated at the estimate (Remark in §S3.10a).

**P-values.** Let $z = (\hat{W}_t - W_t^0)/\kappa_2^{1/2}$, with $W_t^0$ replaced by 0 for the second null. With sample overlap and few effective traits the null distribution of $\hat{W}_t$ is right-skewed and heavy-tailed, and a normal reference is anti-conservative in the upper tail. The p-value therefore matches cumulants:

- $z > 2$: kurtosis-matched central chi-square (Lee et al. 2012). With $\ell = 12\kappa_2^2/\kappa_4$, so that $\chi_\ell^2$ has the excess kurtosis $\kappa_4/\kappa_2^2$ of $\hat{W}_t$, $P = \Pr\{\chi_\ell^2 > \ell + z\sqrt{2\ell}\}$.
- $1.5 < z \leq 2$: shifted gamma matched to $\kappa_1, \kappa_2, \kappa_3$ (Pearson type III).
- $z \leq 1.5$: normal. These tags have $P > 0.07$ under any of the references.

A gamma with skewness $\gamma$ has excess kurtosis $1.5\gamma^2$, but the null $\hat{W}_t$ is more heavy-tailed than that (in the calibration simulation below, median skewness 1.4 and median excess kurtosis 4.6, against 2.9 for a gamma with that skewness). This is why the shifted gamma is used only away from the far tail. Inference uses Gaussian moments; non-Gaussian effects add the cumulant terms of §S5.

**Calibration** (LRCQ paper, Supplementary Table S7): Pan-UKB trait structure, 1000 Genomes LD and $\mathbf{w} \equiv \mathbf{1}$, so every tag is null for $H_0 : W_t \leq W_t^0$; 224 replicates, about 225,000 tests. Entries are rejection rates divided by the nominal level (1 is calibrated).

| **score-type test, p-value** | **0.01** | $10^{-3}$ | $3 \times 10^{-4}$ | $10^{-4}$ |
|---|---|---|---|---|
| normal | 2.93 | 12.7 | 29.6 | 66.1 |
| shifted gamma (3 cumulants) | 1.13 | 2.09 | 2.94 | 4.22 |
| kurtosis-matched $\chi^2$ (4 cumulants) | 0.88 | 1.28 | 1.57 | 1.65 |
| Pearson type VI (4 moments) | 0.93 | 1.28 | 1.39 | 1.60 |

Matching four cumulants keeps the far tail within 1.7-fold of nominal down to $P = 10^{-4}$. The residual excess comes from moments above the fourth.

##### S3.11 Tight LD: what is identifiable, ridge versus tag-set parameterisation

In real LD windows (UK Biobank in-sample LD, HapMap3 SNPs) $\mathbf{D}$ is numerically singular. In a 3 Mb window on chromosome 22, $\mathrm{cond}(\mathbf{D}) \approx 10^{13}$ and a median $(\mathbf{D}^{-1})_{ii}$ of 13, falling to 1.4 after pruning to $r^2 < 0.5$. Write $\bar{\mathbf{y}} = \sum_a s_a(\mathbf{z}_a^{\circ 2} - c_{aa}\mathbf{1})/\|\mathbf{s}\|^2$, so $E\bar{\mathbf{y}} = \mathbf{D}\mathbf{w}$ (no local $\rho$). Let $\mathbf{D} = \sum_k d_k \mathbf{v}_k \mathbf{v}_k^\top$.

**What the data identify.** Only $\mathbf{D}\mathbf{w}$ is identified. Directions $\mathbf{v}_k$ with $d_k \approx 0$ are contrasts among tightly linked SNPs. Within a group of $g$ SNPs in near-perfect LD, $\mathbf{D}$ restricted to the group is $\approx \mathbf{1}\mathbf{1}^\top$. The only identified quantity is the group total $\sum_{j\in\text{group}} w_j$, and how it splits among the SNPs is not identified. *Before any inverse,* project $\hat{\mathbf{D}}$ to the PSD cone by clipping negative eigenvalues. The r²-bias correction can create negative eigenvalues; Schur's theorem guarantees PSD only for the uncorrected $\hat{\mathbf{R}} \circ \hat{\mathbf{R}}$.

**(i) Ridge (Tikhonov).** $\hat{\mathbf{w}}_\lambda = (\mathbf{D} + \lambda\mathbf{I})^{-1}\bar{\mathbf{y}}$.

- Estimand: $E\hat{\mathbf{w}}_\lambda = (\mathbf{D} + \lambda\mathbf{I})^{-1}\mathbf{D}\mathbf{w} = \sum_k \frac{d_k}{d_k+\lambda}\mathbf{v}_k\mathbf{v}_k^\top\mathbf{w}$.
- Bias: $-\lambda(\mathbf{D} + \lambda\mathbf{I})^{-1}\mathbf{w}$. Components along well-determined directions (group totals, $d_k \gg \lambda$) are kept. Within-group contrasts ($d_k \ll \lambda$) are shrunk to 0, so each SNP in a tight group receives about the **group mean** of $w$. Ridge "spreads" a causal SNP's enrichment evenly over its perfect proxies.
- Group totals are approximately unbiased.
- Variance: Theorem 3.10 with $\mathbf{A} = \mathrm{Diag}((\mathbf{D} + \lambda\mathbf{I})^{-1}\mathbf{e}_i)$.
- The variant $(\mathbf{D}^2 + \lambda\mathbf{I})^{-1}\mathbf{D}\bar{\mathbf{y}}$, ridge on the stacked regression, has filter factors $d_k^2/(d_k^2 + \lambda)$ and the same qualitative behaviour.

**(ii) Tag-set (clump-level).** Choose tags $\mathcal{T}$ by greedy pruning at $r^2 < r_0^2$, with responses from all SNPs:

$$\hat{\mathbf{w}}_\mathcal{T} = (\mathbf{D}_{\cdot\mathcal{T}}^\top\mathbf{D}_{\cdot\mathcal{T}})^{-1}\mathbf{D}_{\cdot\mathcal{T}}^\top\bar{\mathbf{y}}, \qquad E\hat{\mathbf{w}}_\mathcal{T} = \mathbf{w}_\mathcal{T} + \mathbf{\Pi}\,\mathbf{w}_{\bar{\mathcal{T}}}, \quad \mathbf{\Pi} = (\mathbf{D}_{\cdot\mathcal{T}}^\top\mathbf{D}_{\cdot\mathcal{T}})^{-1}\mathbf{D}_{\cdot\mathcal{T}}^\top\mathbf{D}_{\cdot\bar{\mathcal{T}}} \tag{3.6}$$

Column $j$ of $\mathbf{\Pi}$ is the least-squares representation of untagged SNP $j$'s $r^2$-profile by the tags' profiles. That allocates $w_j$ to tags:

- *Untagged SNP in perfect LD with one tag:* its column of $\mathbf{D}$ equals the tag's, so $\Pi_{tj} = 1$. Its full $w_j$ goes to the tag, and the tag's estimand is the **clump total**.
- *Untagged causal SNP split between two tags* (partial LD with each): $w_j$ is divided by the projection. The shares need not be non-negative and need not sum to 1. In check T1, allocations sum to 0.98–1.00 for SNPs in tight clusters.
- *Causal SNP absent from the panel:* it behaves exactly like an untagged SNP whose $r^2$-profile is that of its proxies, so the same allocation applies.
- *Variance:* Theorem 3.10 with $\mathbf{A} = \mathrm{Diag}(\mathbf{L}^\top\mathbf{e}_t)$, $\mathbf{L} = (\mathbf{D}_{\cdot\mathcal{T}}^\top\mathbf{D}_{\cdot\mathcal{T}})^{-1}\mathbf{D}_{\cdot\mathcal{T}}^\top$. Pruning bounds the $r^2$ of each pair of tags, not the conditioning of the whole tag design: many tags that are all moderately correlated with one another (for example, $m$ tags with common $r = 0.69$, so every $r^2 < 0.5$) still give a large condition number as $m$ grows. In practice tags are sparse within a window and the median $(\mathbf{D}_{\mathcal{T}\mathcal{T}}^{-1})_{ii}$ is 1.4 in Pan-UKB, but the condition number should be checked per window rather than assumed.

- *Tests:* one-sided score-type tests of $H_0 : W_t \le W_t^0$ and $H_0 : W_t \le 0$ with moment-matched p-values (§S3.10b). The null value $W_t^0$ is the tag estimand under $\mathbf{w} \equiv \mathbf{1}$, not the clump size; the two agree only when the tags span their clumps' LD exactly. Because the projection can assign negative shares, $W_t^0$ can be negative for a tag whose profile is partly explained by its neighbours. It is then a reference value of the projected parameter, not a variance or a clump size. For such tags the score-test null is the nonnegative configuration of §S3.10b. In Pan-UKB this applies to 5.0% of tags (14,978 of 301,693).

**Check T1** (`check_tight_ld.py`, 8 clusters of 4 SNPs at $r \approx 0.97$):

- Both expectation formulas hold (max deviation 0.15 and 0.07 in units of $w$).
- Ridge cluster totals are unbiased (12.6 vs 12.8, 6.7 vs 6.8, 9.7 vs 9.8).
- The tag estimands equal the clump totals (12.8, 6.78, 9.78).
- Per-unit SD is 1.80 for tag-set against 2.68 per SNP for ridge.

**Recommendation.** Report LRCQ at the level of **LD clumps** using the tag-set estimator ($r_0^2 = 0.5$), with the LD-projected tag enrichment $W_t$ of (3.6) as the estimand. $W_t$ equals the clump total when the untagged SNPs are in near-perfect LD with a single tag; in general it is a signed projection whose allocation weights need not be non-negative or sum to 1, and it should be called a clump total, or divided by clump size, only where that holds. Use ridge as a sensitivity analysis. Claims about "omni-SNPs" should then be stated as omni-*loci* (clumps). Resolving which SNP in a clump carries the enrichment is fine-mapping and is not identified from summary statistics plus LD alone; it needs annotations or functional priors (cf. §S7).

The same applies to the joint estimator (3.4) and to the LRCP inverses below.

---

#### S4. LRCP: estimating genetic-effect correlations

##### S4.1 Moment equations

By Corollary 2, cross-block products of blocks 1 and 2 satisfy $E[\mathbf{Z}_1\mathbf{M}\mathbf{Z}_2^\top] = \left(\sum_{a,b} M_{ab}t_{ab}\right)\mathbf{R}_1\mathbf{C}_{12}\mathbf{R}_2$ for any $q \times q$ weight matrix $\mathbf{M}$. The natural choices are:

- $\mathbf{M} = \mathbf{S}$: same-trait products only.
- $\mathbf{M} = \mathbf{T}$: all trait pairs, which uses genetic correlation between traits. This is valid because cross-block $c_{ab}r_{kl} = 0$, so sample overlap does not bias cross-block products.

Within a block, (3.4) applies.

##### S4.2 Full cross-block moment estimator (MOM)

$$\hat{\mathbf{C}}_{12} = \frac{1}{\|\mathbf{T}\|_F^2}\mathbf{R}_1^{-1}\mathbf{Z}_1\mathbf{T}\mathbf{Z}_2^\top\mathbf{R}_2^{-1} \qquad (\text{or } \mathbf{S},\ \|\mathbf{s}\|^2 \text{ for same-trait only}). \tag{4.1}$$

This is the OLS solution of the stacked regression of all cross-block products on $t_{ab}(\mathbf{R}_2 \otimes \mathbf{R}_1)$; the proof is as for Theorem 3.1, with Kronecker algebra. Then $\hat{\rho}_{ij} = \hat{C}_{ij}/\sqrt{\hat{w}_i\hat{w}_j}$. The ratio is defined only for prescreened SNPs with $\hat{w}$ bounded away from 0.

##### S4.3 Prescreened regression estimator (REG)

REG estimates a selected set $\mathcal{P}$ of SNP pairs, with all other $C_{ij} = 0$. The regressors are $x^{(ij)}_{kl,a} = s_a r_{ki} r_{lj}$. The normal equations have the Gram matrix $G_{(ij),(i'j')} = \|\mathbf{s}\|^2 (\mathbf{R}_1^2)_{ii'} (\mathbf{R}_2^2)_{jj'}$ and right-hand side $(\mathbf{R}_1 \mathbf{Z}_1 \mathbf{S} \mathbf{Z}_2^\top \mathbf{R}_2)_{ij}$. So

$$\hat{\mathbf{c}}_{\mathcal{P}} = \frac{1}{\|\mathbf{s}\|^2} \left[ (\mathbf{R}_1^2 \otimes \mathbf{R}_2^2)_{\mathcal{P}\mathcal{P}} \right]^{-1} \left[ \mathbf{R}_1 \mathbf{Z}_1 \mathbf{S} \mathbf{Z}_2^\top \mathbf{R}_2 \right]_{\mathcal{P}}. \tag{4.2}$$

For a single pair this is $\hat{C}_{ij} = (\mathbf{R}_1 \mathbf{Z}_1 \mathbf{S} \mathbf{Z}_2^\top \mathbf{R}_2)_{ij} / (\|\mathbf{s}\|^2 \ell_i \ell_j)$. No sum over the $mq$ or $m^2 q$ data points is needed: averaging over GWAS first gives the same estimate.

It is unbiased if $\mathcal{P}$ contains all non-zero cross-block pairs. Otherwise omitted pairs bias it by $G^{-1}_{\mathcal{P}\mathcal{P}} G_{\mathcal{P}\bar{\mathcal{P}}} \mathbf{c}_{\bar{\mathcal{P}}}$. This bias is small when omitted pairs have small $\sqrt{w_i w_j}$, which is the rationale for prescreening on $\hat{w}$.

##### S4.4 Standard errors, efficiency and power

For $\hat{C} = \sum_a s_a x_a y_a / K$ with $x_a = (\mathbf{A}_1 \mathbf{z}_{1a})_i$ and $y_a = (\mathbf{A}_2 \mathbf{z}_{2a})_j$ (REG: $\mathbf{A} = \mathbf{R}$, $K = \|\mathbf{s}\|^2 \ell_i \ell_j$; MOM: $\mathbf{A} = \mathbf{R}^{-1}$, $K = \|\mathbf{s}\|^2$), with independent traits:

$$\mathrm{Var}(\hat{C}_{ij}) = \frac{1}{K^2} \sum_a s_a^2 \Big[ (\mathbf{A}_1 \mathbf{\Sigma}_{1a} \mathbf{A}_1)_{ii} (\mathbf{A}_2 \mathbf{\Sigma}_{2a} \mathbf{A}_2)_{jj} + (\mathbf{A}_1 \mathbf{\Sigma}_{12,a} \mathbf{A}_2)_{ij}^2 \Big], \tag{4.3}$$

where $\mathbf{\Sigma}_{1a} = \mathbf{R}_1 + s_a \mathbf{R}_1 \tilde{\mathbf{\Sigma}}_{11} \mathbf{R}_1$ and $\mathbf{\Sigma}_{12,a} = s_a \mathbf{R}_1 \tilde{\mathbf{\Sigma}}_{12} \mathbf{R}_2$.

Check P1–P2 compares (4.3) with the empirical SDs: 5.13 vs 5.02 (REG) and 5.75 vs 5.76 (MOM) at $w = 50$, $\rho = 0.5$; and 1.38 vs 1.42 (REG) and 1.99 vs 1.99 (MOM) under the null.

- The background polygenic term $s_a \mathbf{R} \tilde{\mathbf{\Sigma}} \mathbf{R}$ must be kept even under the null. A formula ignoring it underestimated the REG null SE by 40%.
- In small-signal limits, $\mathrm{Var}_{\mathrm{REG}} \approx (\mathbf{R}_1^3)_{ii} (\mathbf{R}_2^3)_{jj} / (\|\mathbf{s}\|^2 \ell_i^2 \ell_j^2)$ and $\mathrm{Var}_{\mathrm{MOM}} \approx (\mathbf{R}_1^{-1})_{ii} (\mathbf{R}_2^{-1})_{jj} / \|\mathbf{s}\|^2$. REG is more efficient when $\mathcal{P}$ is correctly prescreened; MOM is robust to unknown pairs.
- *Power.* Ignoring LD, $\mathrm{SE}(\hat{\rho}_{ij}) \approx 1/(\|\mathbf{s}\| \sqrt{w_i w_j})$. With $\|\mathbf{s}\| = 0.75$ as in §S3.6, a pair with $w_i = w_j = 10$ has $\mathrm{SE} \approx 0.13$, while a pair of average SNPs has $\mathrm{SE} \approx 1.3$. **Genetic-effect correlations are estimable essentially only between enriched SNPs.** This is the quantitative justification for Stage 4's prescreening on $\hat{w}$, and it is why LRCQ precedes LRCP.
- *Regression versus likelihood.* Pooled OLS of Z-score products on LD cross components is unbiased but inefficient, because it ignores the LD-induced covariance among products. The likelihood (§S4.5) and the GLS form (4.4) of §S4.6 use that covariance. The variance of MOM grows with block size and LD through $(\mathbf{R}_1^{-1})_{ii} (\mathbf{R}_2^{-1})_{jj}$. Combining per-GWAS estimates by inverse-variance weighting is (4.2) with trait weights $s_a$, so it does not remove the inefficiency; projecting by $\mathbf{R}^{-1}$ does.

##### S4.5 Likelihood (MLE) and tests

**Likelihood.** Under Gaussian effects and independent traits, $\mathbf{z}_a \sim N(\mathbf{0}, \mathbf{\Sigma}_a(\boldsymbol{\theta}))$ with $\mathbf{\Sigma}_a = c_{aa} \mathbf{R} + s_a \mathbf{R} \tilde{\mathbf{\Sigma}}(\boldsymbol{\theta}) \mathbf{R}$, where $\boldsymbol{\theta}$ collects $\mathbf{w}$ and the prescreened $\rho$. The log-likelihood is $\ell(\boldsymbol{\theta}) =$

$-\frac{1}{2}\sum_a[\log|\boldsymbol{\Sigma}_a| + \mathbf{z}_a^\top \boldsymbol{\Sigma}_a^{-1}\mathbf{z}_a]$ (Zhu & Stephens 2017 for the summary-statistic likelihood, Ning et al. 2020 for its LD-eigen form). With $\mathbf{R} = \mathbf{Q}\boldsymbol{\Lambda}\mathbf{Q}^\top$, keep the leading eigenvectors (HDL) and work with $\tilde{\mathbf{z}}_a = \boldsymbol{\Lambda}_p^{-1/2}\mathbf{Q}_p^\top \mathbf{z}_a$, whose covariance is $c_{aa}\mathbf{I} + s_a\boldsymbol{\Lambda}_p^{1/2}\mathbf{Q}_p^\top \tilde{\boldsymbol{\Sigma}}\mathbf{Q}_p\boldsymbol{\Lambda}_p^{1/2}$.

- The score is $\partial\ell/\partial\theta_u = \frac{1}{2}\sum_a \operatorname{tr}\{(\boldsymbol{\Sigma}_a^{-1}\mathbf{z}_a\mathbf{z}_a^\top\boldsymbol{\Sigma}_a^{-1} - \boldsymbol{\Sigma}_a^{-1})\partial_u\boldsymbol{\Sigma}_a\}$.
- The Fisher information is $\mathcal{I}_{uv} = \frac{1}{2}\sum_a \operatorname{tr}(\boldsymbol{\Sigma}_a^{-1}\partial_u\boldsymbol{\Sigma}_a\boldsymbol{\Sigma}_a^{-1}\partial_v\boldsymbol{\Sigma}_a)$, with $\partial\boldsymbol{\Sigma}_a/\partial\rho_{ij} = s_a\sqrt{w_iw_j}\,\mathbf{R}(\mathbf{e}_i\mathbf{e}_j^\top + \mathbf{e}_j\mathbf{e}_i^\top)\mathbf{R}$.
- In the small-signal limit ($\boldsymbol{\Sigma}_a \approx c_{aa}\mathbf{R}$), $\mathcal{I}_{\rho\rho} \approx \sum_a(s_a/c_{aa})^2 w_iw_j(1+r_{ij}^2)$. The power conclusion of §S4.4 applies to the MLE too.
- Maximise with L-BFGS-B under the bounds $|\rho| < 1$ and $w \geq 0$, using the analytic score.
- *Eigenbasis.* The $\rho$ term of $\boldsymbol{\Sigma}_a$ is $\boldsymbol{\Lambda}(\mathbf{q}_i\mathbf{q}_j^\top + \mathbf{q}_j\mathbf{q}_i^\top)\boldsymbol{\Lambda}$ in the eigenbasis of $\mathbf{R}$, with $\mathbf{q}_i = \mathbf{Q}^\top\mathbf{e}_i$. It has rank 2 and is not diagonal, so once $\rho \neq 0$ the likelihood does not separate into a sum over eigenvalues.
- *Null SNPs.* When some $w_i = 0$ the true value lies on the boundary of the parameter space. If effects are Gaussian given $\mathbf{w}$, $\mathbf{z}$ is exactly Gaussian and the bounded MLE handles this well: in check V1 (10% of $w$ equal to 0, $q = 200$) every null SNP is estimated at 0 and $R^2(\mathbf{w}, \hat{\mathbf{w}})$ is 0.91, against 0.85 for OLS.
- *Non-Gaussian effects.* When effects are sparse *within* a SNP's row of $\mathbf{B}$ (point-normal across traits, §S5), the Gaussian likelihood is a quasi-likelihood. It stays consistent, because the score is unbiased whenever second moments are right, but its information-based SEs are too small. Use the sandwich $\mathcal{I}^{-1}\mathcal{J}\mathcal{I}^{-1}$, with $\mathcal{J}$ from Theorem 3.10 plus the §S5 cumulant term. Do not use a trait jackknife under sample overlap (§S3.6).

**Tests.**

- *Lemma A.* Under $H_0 : \mathbf{C} = \mathbf{0}$ (no genetic-effect correlation), $E[\mathbf{z}_a\mathbf{z}_b^\top] = c_{ab}\mathbf{R} + t_{ab}\mathbf{RWR}$. So the expected Z-score correlation between two SNPs equals their $\mathbf{w}$-weighted cross LD score, which explains why proximal SNPs show correlated Z-scores without any functional link. In general, $E[\mathbf{z}_{1a}\mathbf{z}_{2a}^\top] = t_{aa}\mathbf{R}_{1\cdot}\tilde{\boldsymbol{\Sigma}}\mathbf{R}_{\cdot 2}$: the expected cross-product of two SNPs is a weighted sum of the genetic-effect covariances of their LD partners.
- *Lemma B.* For two blocks without LD, $H_0 : \mathbf{C}_{12} = \mathbf{0}$ implies $E[\mathbf{Z}_1\mathbf{M}\mathbf{Z}_2^\top] = \mathbf{0}$ for every $\mathbf{M}$, with no dependence on $\mathbf{w}$, $h^2$ or intercepts: without a functional link and without LD there is no expected association between the blocks' Z-scores.
- *Sign-flip test (exact under Gaussian $\mathbf{z}$ and independent traits).* Under Lemma B's $H_0$, $\mathbf{z}_{1a}$ and $\mathbf{z}_{2a}$ are independent, so flipping the sign of $\mathbf{z}_{2a}$ independently for each trait leaves the null distribution of any statistic $T(\mathbf{Z}_1\mathbf{S}\mathbf{Z}_2^\top)$ unchanged. For example, use $T = \max_{ij}|\hat{C}_{ij}|/\widehat{\mathrm{SE}}_{ij}$ (block-pair test) or a single prespecified pair. This gives a model-free, multiplicity-adjusted test of block-pair genetic-effect correlation.
- With genetically correlated or overlapping traits, the per-trait sign flips are invalid. First decorrelate traits, $\mathbf{Z} \to \mathbf{Z}\hat{\boldsymbol{\Omega}}^{-1/2}$, with $\hat{\boldsymbol{\Omega}}$ the trait correlation of null-SNP Z-scores. This is the whitening used by HOPS (Jordan et al. 2019).
- *Tests* for a prescreened pair use the variance (4.3) or the information $\mathcal{I}$.
- Tests that compare the spread of pair statistics across a block pair, such as a studentised range or a sample-variance test, are not valid here. The pair statistics share Z-scores and have unequal null variances that depend on their LD scores. The sign-flip max-T test above (independent traits) and the conditional test of §S8.2 (any trait dependence) do not need these assumptions.

##### S4.6 Collapsed GLS for distal SNP sets

Take SNP sets $A$ and $B$ in different LD windows ($r_{kl} = 0$), with candidate subsets $P_A$, $P_B$, and let $\bar{\mathbf{Y}} = \sum_a s_a \mathbf{z}_{A,a} \mathbf{z}_{B,a}^\top / \|\mathbf{s}\|^2$. Then $E\bar{\mathbf{Y}} = \mathbf{R}_A \tilde{\mathbf{\Sigma}}_{AB} \mathbf{R}_B$ (1.5). Regress $\operatorname{vec} \bar{\mathbf{Y}}$ on $\mathbf{R}_B[:, P_B] \otimes \mathbf{R}_A[:, P_A]$.

**Proposition 4.6.**

- a. *Exact algebra.* With working covariance $\mathbf{V} \propto \mathbf{R}_B \otimes \mathbf{R}_A$ (the null, small-signal covariance of the cross-products), the GLS estimator is

$$\hat{\mathbf{C}}_{P_A P_B} = \mathbf{R}_A[P_A, P_A]^{-1} \, \bar{\mathbf{Y}}[P_A, P_B] \, \mathbf{R}_B[P_B, P_B]^{-1}. \quad (4.4)$$

  It uses only the Z-products *of the candidate SNPs themselves*. Taking $P_A = A$ and $P_B = B$ gives MOM (4.1). The OLS fit of the same design is REG (4.2).
- b. *Exact mean, whatever the candidate sets.*

$$E\hat{\mathbf{C}}_{P_A P_B} = \mathbf{R}_A[P_A, P_A]^{-1} \mathbf{R}_A[P_A, :] \, \tilde{\mathbf{\Sigma}}_{AB} \, \mathbf{R}_B[:, P_B] \mathbf{R}_B[P_B, P_B]^{-1}.$$

  It is unbiased iff the signal outside $P_A \times P_B$ is invisible to the candidates. A sufficient condition is that every SNP carrying cross-set correlation is a candidate. Otherwise each candidate absorbs the correlation of non-candidates in LD with it, by the population regression coefficient $\mathbf{R}[P, P]^{-1} \mathbf{R}[P, \bar{P}]$.
- c. *Efficiency.* For a single prescreened pair in the small-signal limit, $\operatorname{Var}(\hat{C}_{ij}) = 1/\|\mathbf{s}\|^2$. This equals the inverse Fisher information of the Stage-4 MLE with $\mathbf{w}$ known (§S4.5, $r_{ij} = 0$). REG and MOM are no more efficient: $\ell_i^2 \leq (\mathbf{R}^3)_{ii}$ by Cauchy–Schwarz, and $(\mathbf{R}^{-1})_{ii} \geq 1$. So (4.4) is the small-signal limit of the Stage-4 MLE/GLS.
- d. *With strong signal* (enriched SNPs, $s_a w \ell \not\ll 1$), $\mathbf{V}$ is no longer $\propto \mathbf{R}_B \otimes \mathbf{R}_A$. Equation (4.4) stays unbiased under (b) but is not fully efficient. Its variance follows from Theorem 3.10 with $\mathbf{A}$ the bilinear form selecting $\bar{Y}[P_A, P_B]$ through the two inverses. Use that rather than the GLS working variance.
- e. *Estimated prescreen.* If candidates are chosen from $\hat{\mathbf{w}}$ computed on the same traits, the null statistic stays centred. Under $\tilde{\mathbf{\Sigma}}_{AB} = \mathbf{0}$, $\mathbf{Z}_A$ and $\mathbf{Z}_B$ are independent (Gaussian case), and selection uses functions of each set separately, so the products remain sign-symmetric. Selection can still change the null variance, so calibration after prescreening is checked by simulation (LRCP paper, simulation S4). Under the alternative, selection on large $\sum_a z_{ka}^2$ can induce a winner's-curse bias; simulation found it negligible. Splitting traits into a screening half and an estimation half removes it at the cost of variance.

*Proof.* (a) $\mathbf{X}^\top \mathbf{V}^{-1} \mathbf{X} = (\mathbf{R}_B[:, P]^\top \mathbf{R}_B^{-1} \mathbf{R}_B[:, P]) \otimes (\cdots) = \mathbf{R}_B[P, P] \otimes \mathbf{R}_A[P, P]$ and $\mathbf{X}^\top \mathbf{V}^{-1} \operatorname{vec} \bar{\mathbf{Y}} = \operatorname{vec}(\mathbf{R}_A[:, P]^\top \mathbf{R}_A^{-1} \bar{\mathbf{Y}} \mathbf{R}_B^{-1} \mathbf{R}_B[:, P]) = \operatorname{vec} \bar{\mathbf{Y}}[P_A, P_B]$. (b) Take expectations in (4.4). (c) At the null with small signal, $\operatorname{Var}(\sum_a s_a z_{ia} z_{ja}) = \sum_a s_a^2$ and $\mathbf{R}[P, P] = 1$ for a single SNP. □

**Check D1** (`check_distal_gls.py`, AR(1) LD, $q = 300$):

- (4.4) equals brute-force GLS to $10^{-14}$.

- With the true pair (5, 9) and candidates {6}×{9}, $E\hat{C}_{6,9}$ = 6.94 against a predicted projection of 7.00 (true $C_{6,9} = 0$).
- With a data-driven top-3 prescreen, the null mean is −0.11 (SD 2.6). Under $C = 10$ the mean is 9.81 without trait splitting and 9.93 with it (SD 2.8 vs 4.0).

**Recommendation:** adopt (4.4) as the default Stage-4 point estimator for distal pairs, taking $P_A$ and $P_B$ to include all prescreened SNPs in each window. Take variances from Theorem 3.10. Trait splitting is optional; it is worth its cost in variance only when the winner's curse under the alternative is material.

##### S4.7 Tight LD among LRCP candidates: tag-projected correlations

In (4.4), $\mathbf{R}[P, P]^{-1}$ is unstable when candidates are in tight LD. Restrict candidates to a tag set ($r^2 < 0.5$ within each window). By Proposition 4.6(b), each tag then estimates the covariance of **R-projected tag effects**, $\tilde{\beta}_t = \sum_j (\mathbf{R}_{PP}^{-1}\mathbf{R}_{P\cdot})_{tj}\beta_j$. That is the tag's own effect plus its linked SNPs' effects weighted by population regression coefficients. With perfect LD the coefficient is $\pm 1$, carrying the allele-alignment sign of $r$.

*Consistent denominators.* For $\hat{\rho}_{tu}$ to be a correlation of the same projected effects, the denominators must use the same projection. Use the within-window collapsed estimator on tags,

$$\hat{\mathbf{V}}_{\mathcal{T}} = \mathbf{R}_{\mathcal{TT}}^{-1}\Big[\sum_a s_a(\mathbf{z}_{\mathcal{T}a}\mathbf{z}_{\mathcal{T}a}^\top - c_{aa}\mathbf{R}_{\mathcal{TT}})\Big]\mathbf{R}_{\mathcal{TT}}^{-1}/\|\mathbf{s}\|^2,$$

whose expectation is $\mathbf{R}_{\mathcal{TT}}^{-1}\mathbf{R}_{\mathcal{T}\cdot}\tilde{\mathbf{\Sigma}}\mathbf{R}_{\cdot\mathcal{T}}\mathbf{R}_{\mathcal{TT}}^{-1}$, and set $\hat{\rho}_{tu} = \hat{C}_{tu}/\sqrt{\hat{V}_{tt}\hat{V}_{uu}}$. Its target is then a genuine correlation with $|\rho| \le 1$. The $\mathbf{D}$-based tag $\hat{w}$ of (3.6) projects with $r^2$ instead of $r$, so it matches only under perfect LD.

**Check T2.** Projected target 0.587. With R-projected denominators the estimate is 0.589. With D-based $\hat{w}$ it is 0.581, a small difference at $r \approx 0.9$ that grows with partial LD. The SNP-level $\rho$ is 0.60.

**Recommendation for Stage 4:** prune candidates to tags and use $\hat{\mathbf{V}}_{\mathcal{T}}$ for the denominators. Interpret results as correlations between LD clumps, which feeds directly into gene-level aggregation (§S8).

---

#### S5. Beyond normality: sparse (point-normal) effects

The first-moment identities (1.2)–(1.6) and the unbiasedness of every linear estimator in §S3–S4 need only second moments. Normality affects variances, weights and likelihood-based inference.

**Proposition 5.1.** Let SNP effects be independent across SNPs within a trait, with $\mathrm{Var}(\beta_{sa}) = v_{sa}$ and fourth cumulant $\kappa_{sa} = E\beta_{sa}^4 - 3v_{sa}^2$. Then

$$\mathrm{Cov}(z_{ka}^2, z_{la}^2) = 2\sigma_{ka,la}^2 + n_a^2\sum_s r_{ks}^2 r_{ls}^2\,\kappa_{sa}, \qquad \mathrm{Var}(z_{ka}z_{la}) = \sigma_{ka}^2\sigma_{la}^2 + \sigma_{ka,la}^2 + n_a^2\sum_s r_{ks}^2 r$$

For point-normal effects with causal probability $\pi$, $\kappa = 3v^2(1-\pi)/\pi$.

*Proof.* Write $\mathbf{z}_a = \boldsymbol{\mu}_a + \mathbf{u}_a$ with $\boldsymbol{\mu}_a = \sqrt{n_a}\mathbf{R}\boldsymbol{\beta}_a$. Expand the fourth moments: Gaussian $\mathbf{u}$ and the cumulant expansion of the linear form $\boldsymbol{\mu}$ give the Gaussian terms plus the fourth-cumulant term

$\mathrm{cum}_4(\mu_k, \mu_k, \mu_l, \mu_l) = n^2 \sum_s r_{ks}^2 r_{ls}^2 \kappa_s$. For spike-and-slab, $E\beta^4 = 3\pi\tau^4$ and $v = \pi\tau^2$. $\square$

Check M2 confirms this to 0.7% at $\pi = 0.1$: $\mathrm{Var}(z^2) = 188$ against a Gaussian value of 46.

**Consequences.**

- i. WLS weights from §S3.5 understate the variance of $z^2$ at SNPs near causal variants, by a factor up to $1 + \frac{3}{2}(1-\pi)/\pi \cdot (\cdot)$. WLS remains unbiased but loses some efficiency. Down-weighting is still in the right direction.
- ii. Analytic SEs (3.3) and (4.3) need the $\kappa$ term. A trait jackknife would not need it, but it is invalid under sample overlap (§S3.6). So add the $\kappa$ term, with $\kappa$ estimated from the fourth moments of the Z-scores (§S5 iii), or use a parametric bootstrap from a point-normal fit.
- iii. Under phenome-wide sparsity ($\beta_{sa} = 0$ for most traits $a$ even at an enriched SNP), the excess kurtosis is across traits. This is the "omni-SNP vs specialist SNP" distinction: the same $w_s$ can arise from small effects on many traits or large effects on few. LRCQ estimates $w_s$, the second moment. The phenome-wide polygenicity $\pi_s$ is identified from fourth moments, e.g. $\pi_s = 3v_s^2/(E\beta^4)$ estimated from $\sum_a (z_{sa}^4 - 6z_{sa}^2 + 3)$ in the spirit of S-LD4M (O'Connor et al. 2019). We leave this extension to future work.
- iv. The MLE is a Gaussian quasi-MLE (§S4.5).

---

#### S6. Summary of estimators and defaults

| Quantity | Estimator | Standard error / test | Section |
|---|---|---|---|
| Intercepts $c_{ab}$, heritabilities, genetic covariances | genome-wide (cross-trait) LDSC, held fixed afterwards | — | S1.7, S3.3 |
| Clump enrichment $\mathbf{w}$ (LRCQ) | tag-set OLS (3.6), $r^2 < 0.5$ tags | model SE, Theorem 3.10 (+ §S5 term), background-filled plug-in for $\mathbf{G}$; score-type tests (nonnegative null, mean in $\mathbf{D}$) with kurtosis-matched p-values | S3.1, S3.10, S3.10a, S3.10b, S3.11 |
| Annotation enrichment $E_c$ | clump-level $\hat{E}_c^{\text{clump}}$ or annotation regression | (7.1) | S7.2, S7.2a, S7.3 |
| Phenotype-category contrast | difference of category estimates | parametric bootstrap of the studentised contrast | S7.4, S7.4a |
| Pair covariance $C_{ij}$ (LRCP) | collapsed GLS (4.4) on prescreened tags | conditional test (8.2) | S4.6, S4.7, S8.2 |
| Pair correlation $\rho_{ij}$ | $\hat{C}_{ij}/\sqrt{\hat{V}_{ii}\hat{V}_{jj}}$ with R-projected denominators | ratio-bias correction | S4.7, S8.4 |
| Gene burden covariance $C_{GH}$ | (8.1) | (8.2) with local $\hat{\mathbf{G}}_A$ and conditional bootstrap; max-T within window pairs, BH/BY across | S8.1–S8.3, S8.2a |
| Pairs from one narrow trait domain | class-null version of (8.2) | report $q_{\text{eff}}(\hat{\mathbf{V}}_{\text{class}})$; descriptive if $< 5$ | S8.5 |

Throughout: never use independence-based SEs or a trait-cluster jackknife under sample overlap (§S3.6), and clip $\hat{\mathbf{w}}$ at 0 before building any plug-in covariance (§S7.4a).

#### S7. Category-level LRCQ: annotations and phenotype categories

Per-SNP (or per-clump) power is low (§S3.6), so most of LRCQ's real-data results come from aggregation over functional annotations and phenotype categories. Everything below is linear in $\mathbf{Y}^{(2)}$, so unbiasedness and exact variances follow from §S3.

##### S7.1 A weighted form of Theorem 3.10

For any trait weights $\mathbf{u} \in \mathbb{R}^q$ and symmetric $\mathbf{A}$, under Gaussian $\mathbf{Z}$ with $\mathrm{Cov}(\mathrm{vec}\,\mathbf{Z}) = \mathbf{\Gamma} \otimes \mathbf{R} + \mathbf{T} \otimes \mathbf{G}$:

$$\mathrm{Var}\left(\sum_a u_a \mathbf{z}_a^\top \mathbf{A} \mathbf{z}_a\right) = 2\left[\tau_{RR}\,\mathbf{u}^\top(\mathbf{\Gamma}\circ\mathbf{\Gamma})\mathbf{u} + 2\tau_{RG}\,\mathbf{u}^\top(\mathbf{\Gamma}\circ\mathbf{T})\mathbf{u} + \tau_{GG}\,\mathbf{u}^\top(\mathbf{T}\circ\mathbf{T})\mathbf{u}\right] \quad (7.1)$$

The proof is that of Theorem 3.10. If $\mathbf{w}$ differs between trait groups, replace $\mathbf{G}$ by the pair-specific $\mathbf{G}_{ab}$ inside the sum, $2\sum_{a,b} u_a u_b\,\mathrm{tr}(\mathbf{A}\mathbf{\Sigma}_{ab}\mathbf{A}\mathbf{\Sigma}_{ba})$. Under a null of equal $\mathbf{w}$, (7.1) applies as written.

##### S7.2 Enrichment of a SNP annotation (set average)

Let $\mathbf{1}_c$ indicate the SNPs (or tags, §S3.11) in annotation $c$. Because $\mathrm{avg}\,\mathbf{w} = 1$, the **category enrichment**

$$E_c = \mathbf{1}_c^\top \mathbf{w}/|c|$$

is directly the fold-enrichment of phenome-wide genetic-effect variance in $c$, the cross-trait analogue of s-LDSC's enrichment (Finucane et al. 2015). The estimator $\hat{E}_c = \mathbf{1}_c^\top \hat{\mathbf{w}}_{\mathrm{OLS}}/|c|$ is unbiased whenever $\hat{\mathbf{w}}$ is: no model for how $\mathbf{w}$ depends on annotations is needed. Its variance is (7.1) with $\mathbf{u} = \mathbf{s}/\|\mathbf{s}\|^2$ and $\mathbf{A} = \mathrm{Diag}(\mathbf{D}^{-1}\mathbf{1}_c)/|c|$. In particular $\tau_{RR} = \mathbf{1}_c^\top \mathbf{D}^{-1}\mathbf{1}_c/|c|^2$, which gives an *effective number of independent SNPs* $|c|_{\mathrm{eff}} = |c|^2/\mathbf{1}_c^\top\mathbf{D}^{-1}\mathbf{1}_c$. Summing over a contiguous LD block is well determined, like local heritability, so $|c|_{\mathrm{eff}}$ is not reduced by LD the way per-SNP precision is. **This holds only when $c$ is a union of LD clumps.** When $c$ splits clumps, use §S7.2a.

Unlike per-trait s-LDSC, this pools traits with the exact cross-trait variance. By Proposition 3.2 the point estimate is the $s_a^2$-weighted average of trait-specific enrichments; the equal-weight version uses $\mathbf{u} = \mathbf{1}/(q\,\mathbf{s})$ element-wise.

##### S7.2a Categories that split tight LD clumps

On real windows ($\mathrm{cond}\,\mathbf{D} \sim 10^{13}$, §S3.11), a category almost always contains some SNPs of a clump and not others (a coding SNP and its non-coding proxies, for example). Write $\mathbf{1}_c = \sum_k (\mathbf{1}_c^\top \mathbf{v}_k)\mathbf{v}_k$ in the eigenbasis of $\mathbf{D}$. Then $\mathrm{Var}\,\hat{E}_c \propto \tau_{RR} = \sum_k (\mathbf{1}_c^\top\mathbf{v}_k)^2/(d_k|c|^2)$. The near-null directions $\mathbf{v}_k$ ($d_k \approx 0$) are within-clump contrasts, and $\mathbf{1}_c$ has a component along them exactly when $c$ splits a clump. The estimator in §S7.2 stays unbiased, but its variance is unbounded as $d_k \to 0$. **The within-clump split of $c$ is not identified from Z and LD alone**, for the same reason the per-SNP split is not (§S3.11). There are three ways to proceed. They differ in their estimand.

| Estimator | Formula | Estimand | When it equals $E_c$ |
| --- | --- | --- | --- |
| Clump-level (recommended, model-free) | $\hat{E}_c^{\text{clump}} = \sum_k f_k \hat{W}_k / \sum_k f_k g_k$, with $\hat{W}_k$ the tag-set estimates (3.6), read as clump totals, $g_k$ the clump size and $f_k = \lvert c \cap k \rvert / g_k$ | $E_c^{\text{clump}} = \sum_k f_k W_k / \sum_k f_k g_k$, the overlap-weighted enrichment of the clumps that $c$ touches | $c$ is a union of clumps, or $\mathbf{w}$ is flat within each clump. Otherwise it is attenuated toward the clump mean |
| Annotation regression (§S7.3) | $\hat{\tau}_0 + \hat{\tau}_1$ with $\mathbf{A} = [\mathbf{1}, \mathbf{1}_c, \ldots]$ | $\boldsymbol{\tau}^*$; under $\mathbf{w} = \mathbf{A}\boldsymbol{\tau}$ it is exactly $E_c$ | The model holds. It borrows strength from clumps in which $f_k$ differs, which is s-LDSC's identification. It is not identified if $f_k$ is the same in every clump |
| Ridge | $\mathbf{1}_c^\top (\mathbf{D} + \lambda \mathbf{I})^{-1} \bar{\mathbf{y}} / \lvert c \rvert$ | $\mathbf{1}_c^\top (\mathbf{D} + \lambda \mathbf{I})^{-1} \mathbf{D} \mathbf{w} / \lvert c \rvert$. Depends on $\lambda$, between $E_c^{\text{clump}}$ and $E_c$ | Not recommended for categories |

Variances all come from (7.1). For the clump estimator use $\mathbf{A} = \text{Diag}(\mathbf{L}^\top \mathbf{f}) / \sum_k f_k g_k$, with $\mathbf{L}$ as in §S3.11 and $\mathbf{f}$ the vector of $f_k$ over tags. For annotation regression use $\mathbf{A} = \text{Diag}(\mathbf{D}\mathbf{A}(\mathbf{A}^\top \mathbf{D}^2 \mathbf{A})^{-1} \mathbf{e})$, with $\mathbf{e}$ the contrast of interest.

Under the null $E_c = 1$ with $\mathbf{w}$ flat within clumps, all three are unbiased, so the tests remain valid. A significant $\hat{E}_c^{\text{clump}}$ means "clumps overlapping $c$ are enriched". Attributing the enrichment to the SNPs in $c$ rather than to their proxies needs the annotation model (s-LDSC's assumption) or fine-mapping.

**Check C2** (`check_category_tight_ld.py`):

- *Setup.* 16 clumps of 4 SNPs at $r \approx 0.99$. The category covers 0 to 4 SNPs per clump. $q = 200$, 3000 replicates.
- *(a)* $\mathbf{w}$ *in the annotation span* ($E_c = 1.50$). Exact $\mathbf{D}^{-1}$: 1.52 (SD 0.82). Ridge: 1.26 (SD 0.27). Clump: 1.19, which is its estimand 1.187 (SD 0.15). Annotation regression: 1.50 (SD 0.26).
- *(b)* $\mathbf{w}$ *on single SNPs, so the model is wrong* ($E_c = 1.71$). Annotation regression: 1.51, its estimand $\boldsymbol{\tau}^*$. Clump: 1.18, its estimand.
- *Variance growth.* Here $\text{cond}\,\mathbf{D} = 180$. The exact estimator's SD grows as $\sqrt{\mathbf{1}_c^\top \mathbf{D}^{-1} \mathbf{1}_c}$, so at real-window conditioning it is unusable.

#### S7.3 Annotation regression (conditional effects)

Model $\mathbf{w} = \mathbf{A}\boldsymbol{\tau}$ with an annotation matrix $\mathbf{A}$ ($m \times K$, first column 1). The stacked OLS is

$$\hat{\boldsymbol{\tau}} = (\mathbf{A}^\top \mathbf{D}^2 \mathbf{A})^{-1} \mathbf{A}^\top \mathbf{D}\, \bar{\mathbf{y}} = (\mathbf{A}^\top \mathbf{D}^2 \mathbf{A})^{-1} \mathbf{A}^\top \mathbf{D}^2\, \hat{\mathbf{w}}_{\text{OLS}},$$

the $\mathbf{D}^2$-weighted projection of $\hat{\mathbf{w}}$ onto the annotation span. Sums over windows enter both factors, and the formula needs no $\mathbf{D}^{-1}$, so it is stable under tight LD. Its estimand is $\boldsymbol{\tau}^* = (\mathbf{A}^\top \mathbf{D}^2 \mathbf{A})^{-1} \mathbf{A}^\top \mathbf{D}^2 \mathbf{w}$. That equals $\boldsymbol{\tau}$ only when $\mathbf{w}$ lies in the annotation span; otherwise it is an LD-weighted projection, as with s-LDSC's $\tau$. Use $E_c$ (§S7.2) for "how enriched is $c$" and $\boldsymbol{\tau}$ for "is $c$ enriched conditional on the other annotations".

##### S7.4 Phenotype-category contrasts

Split traits into categories $g$ (e.g. cardiometabolic, neuropsychiatric) and compute $\hat{\mathbf{w}}^{(g)}$ from the traits in $g$. The category-$g$ enrichment of annotation $c$ is $\hat{E}_c^{(g)}$, and a contrast is $\hat{\Delta}_c = \hat{E}_c^{(g_1)} - \hat{E}_c^{(g_2)}$. This is linear with trait weights $u_a = s_a / \|\mathbf{s}_{g_1}\|^2$ for $a \in g_1$ and $-s_a / \|\mathbf{s}_{g_2}\|^2$ for $a \in g_2$, so (7.1) gives its exact null variance. The two categories' estimates are correlated through cross-category $r_g$ and sample overlap, which (7.1) includes. Per SNP or clump, the vector $(\hat{w}_i^{(g)})_g$ distinguishes **omni-loci**, enriched in all categories, from **category-specific** loci. Test this with a $\chi^2$ test on the contrasts, with covariance from (7.1).

##### S7.4a Plug-in variances under sparse, heavy-tailed enrichment

(7.1) needs $\mathbf{G} = \mathbf{R}\tilde{\boldsymbol{\Sigma}}\mathbf{R}$. With a data-based plug-in $\hat{\mathbf{G}} = \mathbf{R}\,\mathrm{Diag}(\hat{\mathbf{w}})\mathbf{R}$ from the same GWAS, simulated null contrasts gave z-scores with SD 0.81 and no rejections. With the true $\mathbf{w}$ the test was calibrated. Three effects are at work:

1. **Bias of the quadratic term.** $\tau_{GG} = \mathrm{tr}(\mathbf{AGAG})$ is quadratic in $\mathbf{w}$, so

$$E\,\tau_{GG}(\hat{\mathbf{G}}) = \tau_{GG}(\mathbf{G}) + \sum_{s,t} \mathrm{Cov}(\hat{w}_s, \hat{w}_t)\,\left[(\mathbf{RAR})_{st}\right]^2.$$

   The plug-in overstates the variance by the noise in $\hat{\mathbf{w}}$. The **debiased plug-in** subtracts the second term, with $\widehat{\mathrm{Cov}}(\hat{\mathbf{w}}) = \mathbf{D}^{-1}\{2[\mathbf{u}^\top(\boldsymbol{\Gamma} \circ \boldsymbol{\Gamma})\mathbf{u}\,\mathbf{D} + 2\mathbf{u}^\top(\boldsymbol{\Gamma} \circ \mathbf{T})\mathbf{u}\,(\mathbf{R} \circ \hat{\mathbf{G}}) + \mathbf{u}^\top(\mathbf{T} \circ \mathbf{T})\mathbf{u}\,(\hat{\mathbf{G}} \circ \hat{\mathbf{G}})]\}\mathbf{D}^{-1}$, $\mathbf{u} = \mathbf{s}/\|\mathbf{s}\|^2$, and floors the corrected $\tau_{GG}$ at 0. The linear term $\tau_{RG}$ needs no correction.
2. **Self-normalisation.** At a dominant locus, both the contrast and $\hat{w}$ are driven by that locus's realised effects in a few trait clusters (small $q_G$, §S9.1). The studentised statistic then behaves like a $t$ statistic whose numerator and denominator are dependent and have few degrees of freedom, so its tails are too light. No plug-in fixes this; only a reference distribution that includes the plug-in step does.
3. **Indefinite $\hat{\mathbf{G}}$.** Negative $\hat{w}$ make $\hat{\mathbf{G}}$ indefinite. The variance can then be near zero or negative at null SNPs, which produces explosive z-scores. **Always clip $\hat{\mathbf{w}}$ at 0 (or project $\hat{\mathbf{G}}$ to the PSD cone) before using it in a variance.**

**Recommended test: parametric bootstrap of the studentised contrast under $H_0$.**

1. Fit the pooled $\hat{\mathbf{w}}$ and clip it at 0. This is the $H_0$ fit, because under $H_0$ all trait groups share $\mathbf{w}$.
2. Draw $\mathbf{Z}^* \sim N(\mathbf{0}, \boldsymbol{\Gamma} \otimes \mathbf{R} + \mathbf{T} \otimes \hat{\mathbf{G}}_+)$, $B$ times per window.
3. Recompute the whole studentised statistic, including $\hat{\mathbf{w}}^*$ and the debiased plug-in variance, for every contrast and category in the window.
4. Set $p = (1 + \#\{|z^*| \geq |z|\})/(B+1)$.

One set of draws serves all categories and SNPs in a window. A trait split (taking $\hat{\mathbf{G}}$ from held-out traits) also removes the dependence for categories, but it halves the data. For per-locus contrasts it is not a fix, because held-out traits do not share the locus's realised effects.

**Check C3** (`check_contrast_plugin.py`):

- *Setup.* $q = 120$ in 4 clusters with full overlap. Sparse, heavy-tailed $\mathbf{w}$: 6 loci with $w$ between 40 and 80, the rest near 0. Null contrast between clusters {1,2} and {3,4}.

| Statistic | true $\mathbf{w}$ | naive plug-in | debiased plug-in | clipped plug-in | bootstrap (`--boot`) |
|---|---|---|---|---|---|
| Category: SD(z), type-I at 5% | 0.97, 0.051 | 0.84, 0.013 | 0.95, 0.033 | 0.79, 0.005 | type-I 0.050 (0.115 at 10%) |
| Dominant locus | 1.00, 0.063 | 0.79, 0.002 | 0.89, 0.009 | 0.76, 0.001 | 0.015 (0.075 at 10%) |
| Null SNP | 1.01, 0.062 | explodes (indefinite $\hat{\mathbf{G}}$), 0.15 | 0.89, 0.032 | 0.76, 0.009 | 0.045 (0.075 at 10%) |

The bootstrap used 200 replicates with $B = 100$, so its Monte Carlo s.e. is about 0.015.

**Conclusions.**

- For **category** contrasts, the main use in practice, the bootstrap is calibrated. The debiased plug-in is not a safe substitute: it was conservative here (0.033) but anti-conservative in a second configuration ($q = 40$, 4 clusters, 6 dominant loci: 0.073, against 0.040 for the bootstrap). Use the bootstrap; the debiased variance serves only as the studentising denominator inside it.
- For contrasts at a **single dominant locus**, every studentised test is conservative when $q_G$ is small. Report such tests as conservative, and gain power by adding genetically distinct trait clusters rather than more traits.
- The uncorrected plug-in should not be used.

**Check C1** (`check_category.py`):

- *Setup.* 8 LD blocks; $q = 120$ traits in 4 $r_g$-clusters with full sample overlap; sparse $\mathbf{w}$ enriched 3-fold in a 20% annotation; 4000 replicates.
- *Category enrichment.* $E_c$ true 2.178, estimated 2.179. SD: empirical 0.492, (7.1) 0.494, independence formula 0.175. The median per-SNP SD within the annotation is 1.93, so aggregation gains roughly 4-fold in SE here.
- *Annotation regression.* $\hat{\boldsymbol{\tau}}$ mean (0.743, 1.198) against target $\boldsymbol{\tau}^*$ (0.738, 1.213). The ordinary least-squares fit of $\mathbf{w}$, (0.792, 1.386), is *not* the target.
- *Phenotype-category contrast under the null.* Mean 0.002; SD empirical 0.890 vs (7.1) 0.902. Type-I error at 5% is 0.052 with (7.1) and **0.428** with independence-based SEs.

---

#### S8. Gene- and pathway-level LRCP, and multiple testing

The biological questions for LRCP concern functionally related SNPs, gene clusters and pathways. Per-pair power is limited to enriched clumps (§S4.4), and the number of distal pairs is huge (about $2 \times 10^8$ gene pairs). LRCP therefore needs aggregation, conditional tests that survive cross-trait dependence, and explicit error control.

##### S8.1 Gene-level estimands

Let gene $G$ have tags $\mathcal{T}_G$ (§S4.7) in window $A$ and gene $H$ have tags $\mathcal{T}_H$ in a distal window $B$. Fix allele coding (e.g. effect allele = minor allele) and weights $\mathbf{v}_G$ over the window's tags, e.g. the indicator of $\mathcal{T}_G$ or $\hat{w}$-weights. The gene's **burden effect** on trait $a$ is $b_{Ga} = \mathbf{v}_G^\top \tilde{\boldsymbol{\beta}}_{\mathcal{T},a}$, built from the R-projected tag effects of

§S4.7. Its phenome-wide covariance with gene $H$ and its variance are

$$C_{GH} = \mathbf{v}_G^\top \mathbf{C}_{\mathcal{T}_A\mathcal{T}_B} \mathbf{v}_H, \qquad V_G = \mathbf{v}_G^\top \mathbf{V}_{\mathcal{T}_A} \mathbf{v}_G, \qquad \rho_{GH} = C_{GH}/\sqrt{V_G V_H}.$$

The estimator follows from (4.4):

$$\hat{C}_{GH} = \frac{1}{\|\mathbf{s}\|^2} \sum_a s_a \,(\mathbf{g}_G^\top \mathbf{z}_{Aa})(\mathbf{h}_H^\top \mathbf{z}_{Ba}), \qquad \mathbf{g}_G = \mathbf{R}_A[\mathcal{T},\mathcal{T}]^{-1}\mathbf{v}_G,\ \mathbf{h}_H = \mathbf{R}_B[\mathcal{T},\mathcal{T}]^{-1}\mathbf{v}_H. \tag{8.1}$$

with $\hat{V}_G$ from the tag-level collapsed variance (§S4.7). $\hat{C}_{GH}$ is unbiased under Proposition 4.6(b).

*Sign and allele coding.* $C_{GH}$ depends on allele orientation within genes. Two SNPs with opposite effect directions cancel in a burden. That is intended when the hypothesis is "the gene's alleles act together" (e.g. eQTLs aligned to expression-increasing alleles, as for the TP53/MDM2 example). For an orientation-free summary, use the Frobenius statistic $Q_{GH} = \|\hat{\mathbf{C}}_{\mathcal{T}_G\mathcal{T}_H}\|_F^2$ with the conditional null below. Its distribution is a weighted $\chi^2$ (Imhof, or the moment matching of Liu, Tang & Zhang 2009).

#### S8.2 A conditional test valid under any cross-trait dependence

Under $H_0 : \tilde{\boldsymbol{\Sigma}}_{AB} = \mathbf{0}$ with Gaussian Z-scores, $\mathbf{Z}_A$ is independent of $\mathbf{Z}_B$. Condition on $\mathbf{Z}_B$ and write $y_{aH} = \mathbf{h}_H^\top \mathbf{z}_{Ba}$. Then every $\hat{C}_{GH}$ is a linear function of $\mathbf{Z}_A$, and

$$\mathrm{Cov}(\hat{C}_{GH}, \hat{C}_{G'H'} \mid \mathbf{Z}_B) = \frac{1}{\|\mathbf{s}\|^4} \sum_{a,b} s_a s_b\, y_{aH} y_{bH'} \left[c_{ab}\, \mathbf{g}_G^\top \mathbf{R}_A \mathbf{g}_{G'} + t_{ab}\, \mathbf{g}_G^\top \mathbf{G}_A \mathbf{g}_{G'}\right]. \tag{8.2}$$

This is exact for any trait correlation, sample overlap and genetic correlation, *provided the weights $\mathbf{g}_G$ are fixed, window A's covariance ($\boldsymbol{\Gamma}$, $\mathbf{T}$, $\mathbf{G}_A$) is known and correct, and the two windows are independent under $H_0$.* When $\mathbf{G}_A$ is estimated, or allele orientation or gene weights are chosen from the same Z-scores, the statistic is no longer a fixed-coefficient Gaussian form, and the conditional bootstrap below must redo those steps in each draw. Under non-separable effects that covariance is locus-class specific (§S8.5). It needs only window $A$'s second moments ($\boldsymbol{\Gamma}$, $\mathbf{T}$, $\mathbf{G}_A = \mathbf{R}_A\tilde{\boldsymbol{\Sigma}}_A\mathbf{R}_A$, the last from tag-level LRCQ), never the cross-window parameter under test. The per-trait sign-flip test of §S4.5 is the special case for independent traits. Equation (8.2) removes that restriction, and with it the need to whiten traits.

*Multiplicity within a window pair.* The vector of all $\hat{C}_{GH}$ (or tag pairs) is conditionally Gaussian with covariance (8.2). A max-$|T|$ critical value simulated from that Gaussian therefore controls the family-wise error rate for the window pair exactly.

**Check G1** (`check_gene_level.py`): two windows × 3 genes × 3 tags; $q = 120$ traits in 4 $r_g$-clusters ($r_g$ 0.6/0.1) with full sample overlap; enriched SNPs; 1500 null replicates. Per-pair type-I error is **0.050** and the max-$T$ FWER is **0.050**. Under the alternative (one gene pair, $C_{GH} = 4$), the mean $\hat{C}$ is 3.93 (MC s.e. 0.09) and other pairs are within ±0.13 of 0 (MC s.e. 0.06–0.09).

#### S8.2a The window covariance $\mathbf{G}_A$ in the burden null: use the local second moment

(8.2) needs $\mathbf{g}^\top \mathbf{G}_A \mathbf{g}$ with $\mathbf{G}_A = \mathbf{R}_A\tilde{\boldsymbol{\Sigma}}_A\mathbf{R}_A$. Under $H_0$ only the *cross*-window covariance is zero. Within-gene genetic-effect correlation is allowed, and along a burden direction it matters: $\mathbf{v}^\top\tilde{\boldsymbol{\Sigma}}\mathbf{v} = \sum_i w_i +$

$2\sum_{i<j} C_{ij}$.

- A plug-in that keeps only the diagonal of the local $\hat{\hat{\boldsymbol\Sigma}}$ drops the second term. With positively correlated causal SNPs in a gene, the burden test is then anti-conservative: simulations in the LRCP paper gave type-I error of 16–17% at a nominal 5%.
- The LRCQ $\hat{\mathbf{w}}$ partly compensates through its own local-$\rho$ bias $\mathbf{D}^{-1}\,\mathrm{diag}(\mathbf{RCR})$ (Theorem 2). That compensation is accidental, not a guarantee.

**Recommended default: the direct local second moment.**

$$\hat{\mathbf{G}}_A = \frac{1}{\|\mathbf{s}\|^2}\sum_a s_a\big(\mathbf{z}_{Aa}\mathbf{z}_{Aa}^\top - c_{aa}\mathbf{R}_A\big), \qquad E\hat{\mathbf{G}}_A = \mathbf{R}_A\tilde{\boldsymbol\Sigma}_A\mathbf{R}_A = \mathbf{G}_A.$$

- It equals $\mathbf{R}\hat{\hat{\boldsymbol\Sigma}}\mathbf{R}$ for the local estimator (3.4). It needs no $\mathbf{R}^{-1}$, so it is stable under tight LD.
- It makes no assumption about local $\rho$.
- Requirements:
  1. Intercepts $c_{aa}$ fixed from Stage 2. An intercept error $\delta$ shifts $\hat{\mathbf{G}}$ by $-\delta\mathbf{R}$.
  2. Projection of $\hat{\mathbf{G}}_A$ to the PSD cone, by clipping eigenvalues.
  3. *No* quadratic debiasing. (8.2) is linear in $\mathbf{G}_A$, so an unbiased $\hat{\mathbf{G}}_A$ gives an unbiased conditional variance. This differs from §S7.4a, where $\tau_{GG}$ is quadratic.
- *Single-signal genes need no fallback.* $\hat{\mathbf{G}}_A$ is a SNP × SNP moment pooled over all $q$ traits. It is not the $q \times q$ locus trait covariance of §S8.5, which is degenerate for one locus.
- A fallback is needed only when few traits carry signal (say $q_{\text{eff}} < 20$). There, shrink toward the LRCQ form: $\hat{\mathbf{G}}_\lambda = (1-\lambda)\hat{\mathbf{G}}_A + \lambda\mathbf{R}\,\mathrm{Diag}(\hat{\mathbf{w}}_+)\mathbf{R}$.

**Calibration: plug-in alone is conservative; use a conditional bootstrap.** $\hat{\mathbf{G}}_A$ and the numerator are built from the same $\mathbf{Z}_A$. When the gene is enriched, $\mathbf{g}^\top\hat{\mathbf{G}}_A\mathbf{g}$ is large exactly when the realised $\mathbf{g}^\top\mathbf{z}_{Aa}$ are, which is the self-normalisation of §S7.4a. The fix is cheap because the test is conditional on $\mathbf{Z}_B$:

1. Draw $\mathbf{Z}_A^* \sim N(\mathbf{0}, \boldsymbol\Gamma\otimes\mathbf{R}_A + \mathbf{T}\otimes\hat{\mathbf{G}}_{A+})$ with $\mathbf{Z}_B$ held fixed. Each draw is only $|A| \times q$.
2. Recompute $\hat{C}^*$ and $\hat{\mathbf{G}}_A^*$, and studentise.
3. Set $p = (1 + \#\{|z^*| \geq |z|\})/(B+1)$. One set of draws serves every gene pair, and the max-$T$ statistic, in the window pair.

**Check G3** (`check_burden_null.py`):

- *Setup.* AR(1) windows of 9 tags. One gene per window with 3 causal SNPs ($w = 30$) and within-gene correlation 0.6. $q = 100$ and 300 traits in 4 clusters with overlap. 2000 replicates; 300 for the bootstrap.

| $\mathbf{G}_A$ used in (8.2) | type-I at 5%, $q = 100$ / 300 | power at $\rho = 0.5$, $q = 100$ / 300 |
|---|---|---|
| diagonal of local $\hat{\hat{\boldsymbol\Sigma}}$ | **0.101 / 0.100** | 0.78 / 0.80 |
| $\mathbf{R}\,\mathrm{Diag}(\hat{\mathbf{w}}_{\text{LRCQ}})\mathbf{R}$ | 0.040 / 0.039 | 0.72 / 0.75 |
| oracle $\mathbf{G}_A$ | 0.050 / 0.051 | 0.57 / 0.61 |
| local moment $\hat{\mathbf{G}}_A$, plug-in | 0.016 / 0.009 | 0.61 / 0.65 |
| local moment $\hat{\mathbf{G}}_A$, conditional bootstrap | 0.050 / 0.073 | 0.79 / 0.78 |

The bootstrap rows have Monte Carlo s.e. ≈ 0.013.

**Recommendation.** Use $\hat{\mathbf{G}}_A$ from the local second moment (PSD-clipped) in (8.2) for gene, pathway and max-$T$ tests, with p-values from the conditional bootstrap. Under the null this run is calibrated. Under the alternative it was more powerful than the oracle plug-in here, because the oracle's variance includes the signal-inflated $\mathbf{G}_A$ under $H_1$. Never use the diagonal-only projection. Run the test in both orientations ($A|B$ and $B|A$) and report the larger p-value.

##### S8.3 Genome-wide error control: a three-tier design

1. **Screen** genes or clumps on phenome-wide enrichment: tag-level LRCQ with gene-level $\hat{E}_G$ (§S7.2) above a threshold or FDR cut. By Proposition 4.6(e), screening on squares keeps the null of the cross-window products centred and sign-symmetric. Symmetry fixes the centre only: selecting loci with large profiles from the same data can change the spread of the null statistic in either direction, so the calibration after screening has to be checked for the actual pipeline rather than assumed. For the released pipeline this was done by simulation (LRCP paper, simulation S4 and Fig. S2), and the multiplicity correction is then applied to the tests carried forward. Modules and gene pairs defined from the same screen are exploratory localisation, to be confirmed in an independent panel. With $K$ screened genes, the distal pairs number about $K^2/2$, i.e. $10^3$–$10^5$ for $K$ of 50–500, against about $2 \times 10^8$ unscreened.
2. **Test pairs** with (8.2): window-pair max-$T$ inside each window pair, then FDR across window pairs. Window pairs sharing a window are dependent: positively for Gaussian two-sided statistics sharing $\mathbf{Z}_A$. Use Benjamini–Hochberg (1995) as the primary procedure and Benjamini–Yekutieli (2001) as the conservative check.
3. **Pathways (competitive).** For a gene set $\mathcal{S}$, use $S_{\mathcal{S}} = \sum_{G,H\in\mathcal{S},\,\text{distal}} \hat{C}_{GH}/\widehat{\mathrm{SE}}_{GH}$, or the orientation-free $\sum Q_{GH}$. The *self-contained* null $C_{GH} = 0$ uses (8.2), summing the conditional covariances over the window pairs involved. Biology needs a *competitive* null: is $\mathcal{S}$ more co-regulated than random gene sets with the same enrichment and LD profile? Calibrate against matched random sets, sampling genes within strata of $\hat{E}_G$, number of tags and LD score, as MAGMA's competitive test does (de Leeuw et al. 2015). Without matching, any set of enriched genes looks "co-regulated" simply because enriched genes carry more power.

##### S8.4 Clusters of genes or clumps

The gene-by-gene matrix $\hat{\boldsymbol{\rho}}$ among screened genes gives "gene clusters", for example by hierarchical or spectral clustering of $\hat{\boldsymbol{\rho}}$, or community detection on the graph of FDR-significant edges.

- *Ratio bias.* $\hat{\rho} = \hat{C}/\sqrt{\hat{V}_G\hat{V}_H}$ is a ratio of noisy estimates. With independent numerator and denominators, the second-order bias is $E\hat{\rho} - \rho \approx \rho \cdot \frac{3}{8}(\mathrm{CV}_G^2 + \mathrm{CV}_H^2)$, where CV is the coefficient of variation of $\hat{V}$. Denominators near 0 make $\hat{\rho}$ explode. Restrict to genes with $\mathrm{CV}(\hat{V}_G) < 0.3$, about $\hat{E}_G/\,\mathrm{SE} > 3.3$, and report the analytically bias-corrected $\hat{\rho}/[1 + \frac{3}{8}(\widehat{\mathrm{CV}}_G^2 + \widehat{\mathrm{CV}}_H^2)]$, with CVs from Theorem 3.10. A trait-cluster jackknife is not valid under sample overlap (§S3.6).
- *Clustering on* $\hat{\mathbf{C}}$ standardised by its SEs is more stable than clustering on $\hat{\rho}$. Report cluster stability by resampling traits.

**Summary of the gene-level workflow.**

- Screen on LRCQ.
- Estimate gene-level burden covariances with (8.1).
- Test with the conditional Gaussian (8.2): exact under any trait dependence, with max-$T$ inside window pairs and BH/BY across them.
- Run competitive, LD- and enrichment-matched pathway tests.
- Cluster on standardised $\hat{\mathbf{C}}$, with $\hat{\rho}$ reported only for well-estimated genes.

##### S8.5 Non-separable loci: why the GLS and conditional tests disagree, and what to use

In the targeted Pan-UKB analysis of the LRCP paper, the two model-based routes disagree for lipid loci: the conditional test (8.2) gives $z$ about 2.8 times larger than the GLS null SE (PCSK9 × APOB −7.3 vs −2.6).

**(i) What both tests assume.** Write $x_a$ and $y_a$ for the two candidates' projected Z-scores on trait $a$, and $\hat{C} = \mathbf{x}^\top \mathbf{S}\mathbf{y}/K$. Under $H_0$ (the two loci's effects are independent), with $\boldsymbol{\Sigma}_x = \mathrm{Cov}(\mathbf{x})$ and $\boldsymbol{\Sigma}_y = \mathrm{Cov}(\mathbf{y})$ the trait-by-trait covariances:

$$\mathrm{Var}(\hat{C} \mid \mathbf{y}) = \frac{(\mathbf{S}\mathbf{y})^\top \boldsymbol{\Sigma}_x (\mathbf{S}\mathbf{y})}{K^2} \ (\text{conditional, 8.2}), \qquad \mathrm{Var}(\hat{C}) = \frac{\mathrm{tr}(\mathbf{S}\boldsymbol{\Sigma}_x \mathbf{S}\boldsymbol{\Sigma}_y)}{K^2} \ (\text{GLS nul}$$

Under (A2), $\boldsymbol{\Sigma}_x = \tau_R \boldsymbol{\Gamma} + \tau_G \mathbf{T}$: the locus's trait covariance is the genome-wide genetic covariance $\mathbf{T}$, scaled by its enrichment.

- The **conditional** test uses the model for side $A$ only. Side $B$ enters as data.
- The **GLS** test uses the model for both sides, including $\hat{w}_B$.

**(ii) Non-separability makes both anti-conservative.** Suppose a locus acts through a narrow set of mediators, as lipid loci act through LDL. Its trait covariance is then a locus- or class-specific $\mathbf{V}_{\text{class}}$, not $\mathbf{T}$. Two *independent* loci of the same class have random-effect $\rho = 0$. Their realised trait profiles are nevertheless nearly collinear, because $\mathbf{V}_{\text{class}}$ has only $q_{\text{eff}}(\mathbf{V}_{\text{class}}) = (\mathrm{tr}\,\mathbf{V})^2 / \mathrm{tr}\,\mathbf{V}^2$ effective dimensions. This is the finite-$q$ variance of §S2.4, $\mathrm{Var}\,\hat{\rho}^{(q)} \approx 1/q_{\text{eff}}$, with $q_{\text{eff}}$ taken from the *locus class*. A test calibrated with $\mathbf{T}$, where $q_{\text{eff}}(\mathbf{T})$ is about 11 for the 444 Pan-UKB traits, reads that collinearity as strong evidence of correlation. Simulations in the LRCP paper show exactly this: two independent loci on the same trait cluster are rejected 9–31% of the time by both tests, against 2–4% when they hit different clusters.

**(iii) Why the two tests then disagree.** In expectation over $\mathbf{y}$ under the model, the two variances are equal. Their ratio in data is

$$\mathcal{R}_{AB} = \frac{(\mathbf{S}\mathbf{y})^\top \boldsymbol{\Sigma}_x (\mathbf{S}\mathbf{y})}{\mathrm{tr}(\mathbf{S}\boldsymbol{\Sigma}_x \mathbf{S}\boldsymbol{\Sigma}_y)}, \qquad z_{\text{cond}}/z_{\text{GLS}} = \mathcal{R}_{AB}^{-1/2}.$$

Under the model $\mathcal{R} \approx 1 \pm \sqrt{2/q_{\text{eff}}}$. A 2.8-fold gap means $\mathcal{R} \approx 0.13$; the observed values for the lipid pairs are 0.12–0.16, far off-model. The GLS denominator contains $\boldsymbol{\Sigma}_y$, and so $\hat{w}_B$ (or $\hat{\mathbf{V}}_{\mathcal{T},B}$). Two things inflate it relative to the realised $\mathbf{y}$:

- A noisy, up-clipped $\hat{w}_B$, such as $\max(\hat{w}_{\text{raw}}, 0)$ with $\hat{w} = 321$ and SE 744 at rs78378222.
- The $r^2$-versus-$r$ projection mismatch of §S4.7.

The conditional test uses the realised $\mathbf{y}$ and is immune to both. Non-separability also enters: the conditional variance is too small whenever $\mathbf{y}$ lies in a direction where $\mathbf{T}$ understates the locus-class variance. So the gap is a diagnostic of misspecification. It is not evidence that one route is right.

**Diagnostic to report for every pair:**

- $\mathcal{R}_{AB}$, and the side-wise signal check $e_B = (\mathbf{y}^\top \mathbf{S}\mathbf{y} - \operatorname{tr} \mathbf{S}\mathbf{\Gamma}_{\text{proj}}) / \operatorname{tr}(\mathbf{S}\, \tau_{G,B} \mathbf{T})$, the realised over the model-predicted signal energy. $e_B \ll 1$ means $\hat{w}_B$ is overstated.
- The **class effective trait count** $q_{\text{eff}}(\hat{\mathbf{V}}_{\text{class}})$ (below).

**(iv) Locus-specific covariance: what works and what does not.**

- *From the block's own profile.* $\hat{\mathbf{V}}_A \propto \mathbf{x}\mathbf{x}^\top - \mathbf{\Gamma}$ is rank one when the block carries one signal. The conditional variance then becomes $\approx (\mathbf{x}^\top \mathbf{S}\mathbf{y})^2$, so $|z| \leq 1$ by Cauchy–Schwarz and the test has no power. A block's own tags help only if they carry several *independent* signals, and then $\hat{\mathbf{V}}_A$ has that many degrees of freedom.
- *A per-locus effective trait count.* For the same reason, $q_G$ computed from one block's own trait covariance is always about 1. It is not informative.
- **From held-out loci of the same class (recommended).** Cluster the enriched tags genome-wide by their normalised trait profiles, for example by hierarchical clustering of $\operatorname{sign}$-free profile correlations $|\operatorname{corr}(\mathbf{x}_t, \mathbf{x}_{t'})|$. For a pair $(A, B)$:
  1. Estimate $\hat{\mathbf{V}}_{\text{class}} = \operatorname{avg}_{t \in \text{class} \setminus \{A,B,\text{their blocks}\}} (\mathbf{x}_t \mathbf{x}_t^\top - \tau_{R,t} \mathbf{\Gamma}) / \tau_{G,t}$.
  2. Use $\mathbf{\Sigma}_x = \tau_R \mathbf{\Gamma} + \kappa_A \hat{\mathbf{V}}_{\text{class}}$, with $\kappa_A$ trace-matched to the realised signal of $\mathbf{x}$, in (8.2). Do the same on both sides for the GLS SE. This tests the random-effect $\rho = 0$ *relative to the class*: "are A and B more aligned than two independent loci of the same trait domain?". When $A$ and $B$ fall in different classes, or the class is the genome-wide one, it reduces to the current tests.
- **Interpretation when $q_{\text{eff}}(\hat{\mathbf{V}}_{\text{class}})$ is small** (below about 5). Profile alignment within the class is then not testable against the class null, whatever the method. The realised $\hat{\rho}^{(q)} \approx \pm 1$ is the mediator-model signature of the two loci acting through the same mediator (§S2.3). It should be reported as such, descriptively: "both loci act on the LDL axis, with opposite (or equal) directions". It should not be reported as a significant random-effect correlation. The sign concordance with known biology is the substantive result.

**Check G2** (`check_nonseparable.py`):

- *Setup.* Mediator model with 25 mediators. The genome-wide $\mathbf{T}$ has $q_{\text{eff}} = 18.5$; a two-mediator "lipid class" has $q_{\text{eff}} = 1.9$. $q = 100$ traits with overlap, 2000 replicates.
- *Separable test (null covariance from* $\mathbf{T}$*).* Under $H_0$ (two independent class loci; median realised |profile corr| 0.73) it rejects **0.70** at 5% (median $|z|$ 3.1).
- *Class-null test* ($\hat{\mathbf{V}}_{\text{class}}$ from 40 held-out class loci). It rejects 0.004 under $H_0$. Under $\rho = 1$ and $\rho = 0.7$ within the class it rejects 0.005 and 0.003: valid, but with no power, as $q_{\text{eff}} = 1.9$ predicts.
- *Own-block covariance.* $\max |z| = 1.00$ in every scenario, as the Cauchy–Schwarz bound predicts.

**Recommendations.**

1. Keep the conditional test (8.2) as the primary per-pair test. It is the more robust of the two, because it does not depend on $\hat{w}_B$. Report $\mathcal{R}_{AB}$ alongside it.
2. Add the class-null version (8.2 with $\hat{\mathbf{V}}_{\text{class}}$) as the test of record for loci whose profile class is not the genome-wide one. Report $q_{\text{eff}}(\hat{\mathbf{V}}_{\text{class}})$ with each pair.
3. When $q_{\text{eff}} < 5$, present $\hat{\rho}$ descriptively as shared-mediator alignment. The lipid pairs fall here.
4. Never use a diagonal or Cheverud-style genetic covariance. Use the genome-wide LDSC $\mathbf{T}$ (rg-based). A diagonal $\mathbf{T}$ produced 2,166 false positives in a Pan-UKB test run.
5. Drop the trait-cluster jackknife for targeted pairs. It needs many comparable clusters, and here one cluster carries the signal.

---

#### S9. Power and sample size

The tables below are produced by `theory/power/power_tables.py`, with a Monte Carlo spot-check in `theory/power/spotcheck_power.py`.

Every number below follows from Theorem 3.10 / (7.1). Trait heterogeneity enters through $s_a = n_a h_a^2/m$, with $m = 1.2 \times 10^6$ fixing the per-SNP scale. LD enters through $\kappa_{\text{LD}} = (\mathbf{D}^{-1})_{ii}$ for tags, which is 1.4 after $r^2 < 0.5$ pruning (§S3.11).

##### S9.1 Three effective numbers of traits

With equal $s$, Theorem 3.10 reduces to

$$\operatorname{Var}\hat{w}_i = \frac{2\kappa_{\text{LD}}}{s^2}\left[\frac{1}{q_P} + \frac{2sw_i}{q_{PG}} + \frac{(sw_i)^2}{q_G}\right], \qquad \operatorname{Var}\hat{C}_{ij} = \frac{1}{s^2}\left[\frac{1}{q_P} + \frac{s(w_i+w_j)}{q_{PG}} + \frac{s^2(w_i w_j \text{[illegible]}}{q_G} \right. \tag{9.1}$$

with $q_P = q^2/\sum_{ab}\Gamma_{ab}^2$ (sample overlap × phenotypic correlation), $q_G = q^2/\sum_{ab} r_{g,ab}^2$ and $q_{PG} = q^2/\sum_{ab}\Gamma_{ab} r_{g,ab}$.

The null SE depends only on $q_P$. A large SNP effect makes the $q_G$ term dominate, so power at large $w$ is governed by how many *genetically* distinct traits there are. With $q$ traits in $K$ equal clusters, within-cluster correlation $r_w$ and between-cluster $r_b$, $q_{\text{eff}} = q^2/\{q[1+(q/K-1)r_w^2] + q(q-q/K)r_b^2\}$. For $q = 300$:

| **Clusters $K$** | $r_w = 0.2$ | $r_w = 0.4$ | $r_w = 0.6$ | $r_w = 0.4, r_b = 0.1$ |
|---|---|---|---|---|
| 5 | 89 | 29 | 14 | 23 |
| 10 | 139 | 53 | 26 | 36 |
| 20 | 192 | 93 | 50 | 49 |
| 50 | 250 | 167 | 107 | 63 |

A small between-cluster correlation caps $q_{\text{eff}}$ near $1/r_b^2$, whatever $q$ is. The simulated trait structures of the LRCP paper (16 to 120 of 300 traits) sit inside this range; in Pan-UKB, $q_{\text{eff}}$ is 58 of 452 traits for the sampling-noise correlation and about 11 of 444 for the genetic covariance. **Collecting more GWAS of correlated traits buys little. Adding genetically distinct trait domains is what raises power.**

**Check S9** (`spotcheck_power.py`):

- *Setup.* $q = 200$ in 10 overlap clusters, $r_w = 0.4$, so $q_P = 49.5$ and $q_G = 200$. 4000 replicates.
- *SD of* $\hat{w}$. At $w = 1$: simulated 4.09, (9.1) 4.07. At $w = 20$: simulated 5.29, (9.1) 5.31. The single-$q_{\text{eff}}$ shortcut gives 8.04.
- $\hat{C}$. Mean 10.03 for true 10.0; SD simulated 3.74, (9.1) 3.82.

The tables below set $q_P = q_G = q_{PG} = q_{\text{eff}}$, which is conservative when genetic correlations are weaker than phenotypic ones.

##### S9.2 Per-clump LRCQ

One-sided test of $w = 0$ at $5 \times 10^{-7}$ (about $10^5$ clumps), $h^2 = 0.1$. Each cell gives the null SE, then power at $w = 10$ / 20 / 50.

| $n$ $q_{\text{eff}}$ | 50 | 100 | 300 | 1000 |
|---|---|---|---|---|
| $10^5$ | 28.4: 0 / 0 / .01 | 20.1: 0 / 0 / .05 | 11.6: 0 / 0 / .34 | 6.4: 0 / .07 / .98 |
| $4 \times 10^5$ | 7.1: 0 / .11 / .79 | 5.0: .02 / .29 / .97 | 2.9: .14 / .89 / 1 | 1.6: .85 / 1 / 1 |
| $10^6$ | 2.8: .23 / .79 / .99 | 2.0: .52 / .97 / 1 | 1.2: .98 / 1 / 1 | 0.6: 1 / 1 / 1 |

The SE scales as $1/(\sqrt{q_{\text{eff}}}\, nh^2)$, so **doubling $n$ is worth quadrupling $q_{\text{eff}}$**. With UK Biobank-sized GWAS ($n \approx 4 \times 10^5$) and $q_{\text{eff}} \approx 100$, only omni-loci with $w \gtrsim 30$ are detectable genome-wide. Per-clump LRCQ is a discovery tool for the strongest loci only. This confirms §S3.6 and is the case for category-level analysis and prescreened LRCP.

*Two regimes.* When per-SNP signal $s$ is large, the third term of (9.1), $2\kappa w^2/q_G$, dominates; it does not involve $n$, so precision is then limited by the number of genetically distinct traits rather than by GWAS sample size. At realistic genome sizes $s \propto 1/m$ is small, the first term dominates, and the null variance grows as $m^2$. Small simulations with $m$ equal to the block size are therefore far more optimistic than genome-scale analyses.

##### S9.3 Category LRCQ

Test of $E_c = 1$ at $5 \times 10^{-4}$ (100 annotations), $n = 4 \times 10^5$, $h^2 = 0.1$. Each cell gives power at $E_c = 1.5$ / 2 / 3. Here $|c|_{\text{eff}}$ is the number of effectively independent clumps (§S7.2).

| $q_{\text{eff}}$ $\lvert c\rvert_{\text{eff}}$ | 100 | 1000 | 10000 |
|---|---|---|---|
| 50 | .01 / .05 / .48 | .23 / .96 / 1 | 1 / 1 / 1 |
| 100 | .02 / .16 / .88 | .62 / 1 / 1 | 1 / 1 / 1 |
| 300 | .10 / .74 / 1 | 1 / 1 / 1 | 1 / 1 / 1 |

Annotations covering ≥1000 effectively independent clumps (about 1% of the genome) reach full power for 2-fold enrichment even at $q_{\text{eff}} = 50$. Category LRCQ is therefore the best-powered use of LRCQ.

##### S9.4 Pair and gene-level LRCP

Two-sided test of $\rho = 0$ between two distal tags with equal enrichment $w$. Bonferroni over the $K(K-1)/2$ pairs among $K$ prescreened clumps, $h^2 = 0.1$. Each cell gives the null SE of $\hat{\rho}$, then power at $\rho = 0.5$ for $K = 100$ / 1000.

| | $w = 10$ | $w = 20$ | $w = 50$ |
|---|---|---|---|
| $n = 4 \times 10^5$, $q_{\text{eff}} = 300$ | 0.23: .01 / 0 | 0.14: .18 / .03 | 0.09: .83 / .53 |
| $n = 4 \times 10^5$, $q_{\text{eff}} = 1000$ | 0.13: .32 / .09 | 0.08: .97 / .84 | 0.05: 1 / 1 |
| $n = 10^6$, $q_{\text{eff}} = 300$ | 0.13: .32 / .09 | 0.09: .83 / .53 | 0.07: .99 / .94 |

At $\rho = 0.2$, power exceeds 0.5 only when $n = 10^6$, $q_{\text{eff}} = 1000$ and $w = 50$.

- **Gene and pathway level (§S8).** A burden $C_{GH} = \mathbf{v}_G^\top \mathbf{C} \mathbf{v}_H$ behaves as one pair with $w$ replaced by the gene totals $W_G = \mathbf{v}_G^\top \mathbf{w}$, provided the correlations within the pair of genes share a sign. Aggregating $g$ clumps per gene with coherent $\rho$ multiplies the noncentrality by about $g$. Mixed-sign correlation cancels.
- **Design rules:**
    1. Prescreen to $K \leq 100$ clumps with $\hat{w} \geq 10$ from LRCQ. This costs no power, because the alternative needs large $w$ anyway, and the multiple-testing burden falls by $K^2$.
    2. Report the $\rho$ detectable at 80% power for the realised $(q_{\text{eff}}, n, w)$.
    3. Treat $|\rho| < 0.3$ as below resolution for the current Pan-UKB data.

---

#### Appendix A. Numerical verification

Each check is a Python script (run with `python3 <script>.py`). The scripts are available from the corresponding author on request and will be made publicly available upon journal publication. The estimators themselves are implemented in lrcpq (https://github.com/unfated/lrcpq).

| ID | Script | What is checked | Result |
|---|---|---|---|
| M1 | `check_master_identity.py` | (1.2) from individual-level data: 6 SNPs with LD, 2 traits, $n_a = 3000$, $n_b = 2000$, $n_{ab} = 1200$, residual correlation 0.4, $r_g = 0.5$, non-normal effects, one correlated pair; 6000 replicates | median relative error of the genetic part 3.5% (expected $O(h^2)$ at per-SNP $h^2 \approx 0.7\%$); $Ez^2$: 52.2 empirical, 52.8 theory, 59.1 double-counted form |
| F1 | `check_factor2.py` | cross-block regression design of §S1.5, 400 reps | slope ≈ ρ with pairs counted once; exactly ρ/2 with a double-counted regressor (table in §S1.5) |
| L1 | `check_lrcq.py` | fast OLS (3.1) vs stacked OLS | max difference $1.6 \times 10^{-14}$ |
| L2 | `check_lrcq.py` | fast WLS (3.2) vs stacked WLS | max difference $1.3 \times 10^{-13}$ |
| L3 | `check_lrcq.py` | bias (1.6) with local ρ = 0.6 between nearby causal SNPs | corr(observed, predicted) 0.999, slope 1.02; no bias without local ρ |
| L4 | `check_lrcq.py` | Proposition 3.2, trait-specific $\mathbf{w}_a$ | OLS = $s^2$-weighted mean (corr 1.000); differs from simple mean by up to 0.80 |
| L5 | `check_lrcq.py` | Proposition 3.3 | rank 99 of 100; null direction $(\mathbf{D}^{-1}\mathbf{1}, -\mathbf{s})$ |

| ID | Script | What is checked | Result |
|---|---|---|---|
| P1–P2 | `check_lrcp.py` | REG and MOM unbiased; SE (4.3) | $C = 25$: REG 24.82, MOM 24.96; SD vs (4.3): 5.02/5.13, 5.76/5.75; null 1.42/1.38, 1.99/1.99 |
| P3 | `check_lrcp.py` | (3.4) within-block $\tilde{\Sigma}$ | $\hat{w}$ = 50.0, 49.9 (true 50); rest mean 1.00 |
| M2 | `check_moments.py` | Isserlis and Proposition 5.1 | max relative error 0.5% (normal), 0.7% (π = 0.1) |
| X1 | `check_crosstrait.py` | Theorem 3.10, correlated and overlapping traits | SD 18.6 vs (3.5) 18.3 (LRCQ), 14.8 vs 14.7 (LRCP); independence formula 6.7, 5.3 ( $q_{\text{eff}} \approx 16$) |
| D1 | `check_distal_gls.py` | Proposition 4.6 | GLS identity to $10^{-14}$; projection bias 6.94 vs 7.00 predicted; prescreen null mean −0.11, alternative 9.81 (true 10) |
| T1 | `check_tight_ld.py` | §S3.11 ridge and tag-set estimands | formulas hold (max dev 0.15, 0.07); tag estimand = clump totals 12.8/6.78/9.78; SD 1.80 tag vs 2.68 ridge |
| T2 | `check_tight_ld.py` | §S4.7 tag-projected ρ | target 0.587; R-projected 0.589; D-based 0.581 |
| C1 | `check_category.py` | §S7 category enrichment, annotation regression, category contrast | $E_c$ 2.179 (true 2.178), SD 0.492 vs 0.494; $\hat{\tau}$ = target; contrast type-I 0.052 (independence SEs: 0.428) |
| C2 | `check_category_tight_ld.py` | §S7.2a categories splitting tight clumps | clump estimator 1.188 = estimand 1.187, SD 0.15 vs 0.82 for exact $\mathbf{D}^{-1}$; annotation regression 1.501 for $E_c$ 1.500 when $\mathbf{w} = \mathbf{A}\boldsymbol{\tau}$ |
| C3 | `check_contrast_plugin.py` | §S7.4a plug-in variance for contrasts under sparse $\mathbf{w}$ | category type-I: naive 0.013, debiased 0.033, bootstrap 0.050 (true-$\mathbf{w}$ 0.051) |
| G2 | `check_nonseparable.py` | §S8.5 pair tests under non-separable (mediator-class) effects | separable-null type-I 0.70; class-null 0.004 (no power at $q_{\text{eff}} = 1.9$); own-block $\lvert z \rvert \leq 1$ |
| G3 | `check_burden_null.py` | §S8.2a window covariance in the burden null | diagonal-only 0.10; local moment + conditional bootstrap 0.050/0.073, power 0.79 |
| J1 | `check_jackknife_overlap.py` | §S3.6 trait-cluster jackknife under overlap | jackknife SE 2.55 vs empirical 7.31 at between-cluster overlap correlation 0.2; model (3.5) 7.30 |
| S9 | `power/spotcheck_power.py` | §S9 variance (9.1) with distinct $q_P$, $q_G$ | SD $\hat{w}$ 5.29 vs 5.31 at $w = 20$ (single $q_{\text{eff}}$: 8.04); SD $\hat{C}$ 3.74 vs 3.82 |

| ID | Script | What is checked | Result |
|---|---|---|---|
| G1 | `check_gene_level.py` | §S8 conditional Gaussian test and gene-level burden | per-pair type-I 0.050, max-T FWER 0.050 with correlated, overlapping traits; $\hat{C}_{GH}$ 3.93 (true 4, MC s.e. 0.09) |
| V1 | `check_mle_null_w.py` | §S4.5: bounded MLE of $\mathbf{w}$ with 10% $w = 0$ (20 SNPs, $q = 200$, Gaussian effects), 50 reps | all converge; every null at $\hat{w} < 0.05$ (mean 0.000); $R^2$ MLE 0.905, OLS 0.854 |
| P4 | `check_se_plugin.py` | §S3.10a enrichment plug-in in the model SE of $\hat{w}_i$, dense to very sparse $\mathbf{w}$, mean LD score 2.8–15.7 | null z SD: method A up to 1.68 (type-I 0.19); background-filled method A 0.90–1.02; true $\mathbf{w}$ 1.00–1.02 |
| M3 | `check_moments.py` | Proposition 2.3 | Cov(vec B) = U⊗V to 0.003; $EB^4$ = 35.8 (theory 36; MN 12) |


#### Acknowledgements

We thank Justin D. Tubbs for contributions to an earlier version of this work.

*Relation to existing methods.*

- **s-LDSC** estimates trait-specific enrichment of annotation categories. LRCQ estimates SNP-level enrichment pooled across traits, without annotations (§S1.4).
- **HOPS** scores per-SNP pleiotropy from whitened Z-scores but does not deconvolve LD. LRCQ removes LD through $\mathbf{D}^{-1}$ and is unbiased for $\mathbf{w}$ (§S3.1).
- **Local genetic correlation** methods (SUPERGNOVA, LAVA) estimate correlation between *traits* within a locus. LRCP estimates correlation between *SNPs* across traits, the transpose problem (§S2.4).
- **DeGAs** factorises the Z-score matrix without modelling LD. The mediator model (§S2.3) gives the matching generative interpretation.
